# Origin and Evolution of Pseudomurein Biosynthetic Gene Clusters

**DOI:** 10.1101/2022.11.30.518518

**Authors:** Valérian Lupo, Célyne Roomans, Edmée Royen, Loïc Ongena, Olivier Jacquemin, Frédéric Kerff, Denis Baurain

## Abstract

The peptidoglycan (PG; or murein) is a mesh-like structure, which is made of glycan polymers connected by short peptides and surrounds the cell membrane of nearly all bacterial species. In contrast, there is no PG counterpart that would be universally found in Archaea, but rather various polymers that are specific to some lineages. Methanopyrales and Methanobacteriales are two orders of Euryarchaeota that harbor pseudomurein (PM) in their cell-wall, a structural analogue of the bacterial PG. Owing to the differences between PG and PM biosynthesis, some have argued that the origin of both polymers is not connected. However, recents studies have revealed that the genomes of PM-containing Archaea encode homologues of the bacterial genes involved in PG biosynthesis, even though neither their specific functions nor the relationships within the corresponding inter-domain phylogenies have been investigated so far. In this work, we devised a bioinformatic pipeline to identify all potential proteins for PM biosynthesis in Archaea without relying on a candidate gene approach. After an *in silico* characterization of their functional domains, the taxonomic distribution and evolutionary relationships of the collected proteins were studied in detail in Archaea and Bacteria through HMM similarity searches and phylogenetic inference of the Mur domain-containing family, the ATP-grasp superfamily and the MraY-like family. Our results notably show that the extant archaeal muramyl ligases are ultimately of bacterial origin, but likely diversified through a mixture of horizontal gene transfer and gene duplication. Moreover, structural modeling of these enzymes allowed us to propose a tentative function for each of them in pentapeptide elongation. While our work clarifies the genetic determinants behind PM biosynthesis in Archaea, it also raises the question of the architecture of the cell wall in the last universal common ancestor.

## Introduction

The cell wall is a complex structure that surrounds most prokaryotic cells, protects them against the environment and maintains their internal turgor pressure (Pazos and Peters 2019; Meyer and Albers 2020). It also constitutes one of the striking phenotypic differences between Archaea and Bacteria. Indeed, while most archaeal species possess a paracrystalline protein surface layer (S-layer; Rodrigues-Oliveira et al. 2017), other species harbor a large variety of cell-wall polymers (e.g., sulfated heteropolysaccharides, glutaminylglycan, methanochondroitin) (Albers and Meyer 2011; Meyer and Albers 2020), whereas nearly all bacterial cell walls contain a single common polymer termed peptidoglycan (PG; also known as murein) (Vollmer et al. 2008; Pazos and Peters 2019). PG is a net-like polymer (Fig. 1) formed by long glycosidic chains of alternating N-acetylglucosamine (GlcNAc) and N-acetylmuramic acid (MurNAc) units linked by a β-(1→4) bond. To MurNAc is attached a short peptide, from three to five amino acids (AA) long, usually composed of L-alanine (L-Ala), D-glutamic acid (D-Glu), meso-diaminopimelic acid (meso-DAP) or L-lysine (L-Lys), and two D-alanines (D-Ala). This short peptide serves as a bridge between two glycosidic chains and is built at the final stage of PG biosynthesis (Vollmer et al. 2008; Pazos and Peters 2019). Interestingly, there exists an archaeal cell wall polymer that shows a three-dimensional structure similar to PG, hence named pseudopeptidoglycan or pseudomurein (PM). Compared to PG, PM (Fig. 1) contains N-acetyl-L-talosaminuronic acid (NAT) units linked to GlcNAc through a β-(1→3) bond, instead of MurNAc, and only has L-amino acids attached to NAT (König et al. 1982; König et al. 1993; Meyer and Albers 2020). Depending on the species, both PG and PM can show variation in their amino acids and glucidic composition (König et al. 1982; Vollmer et al. 2008; Pazos and Peters 2019; Meyer and Albers 2020). In the early 1990s, a PM biosynthesis pathway was proposed (Hartmann and König 1990; König et al. 1993; Hartmann and König 1994) and, due to differences between PG and PM biosynthesis, it was concluded that both polymers had evolved independently (Kandler and Konig 1993; Scheffers and Pinho 2005; Albers and Meyer 2011). In contrast to the ubiquity of PG, PM is found only in two orders of Euryarchaeota: Methanopyrales and Methanobacteriales. In recent phylogenomic reconstructions, Methanopyrales and Methanobacteriales are both monophyletic and further form a clade with Methanococcales as an outgroup, all three orders being collectively termed class I methanogens (CIM) (Bapteste et al. 2005; Williams et al. 2020). Unlike Methanopyrales and Methanobacteriales, the cell wall of Methanococcales is composed of an S-layer and does not contain PM. This restricted taxonomic distribution suggests that PM has appeared in the last common ancestor (LCA) of these two orders of methanogens, after their separation from the Methanococcales lineage, and thus that PM was not a feature of a more ancient archaeal ancestor. In other studies, Methanopyrales are basal to the whole clade of CIM (Williams et al. 2020; Aouad et al. 2022), which would point to a loss of PM in Methanococcales. However, it has been proposed that the latter topology might be caused by a long-branch attraction (LBA) artifact (Gribaldo et al. 2006; Da Cunha et al. 2018).

**Figure 1.**
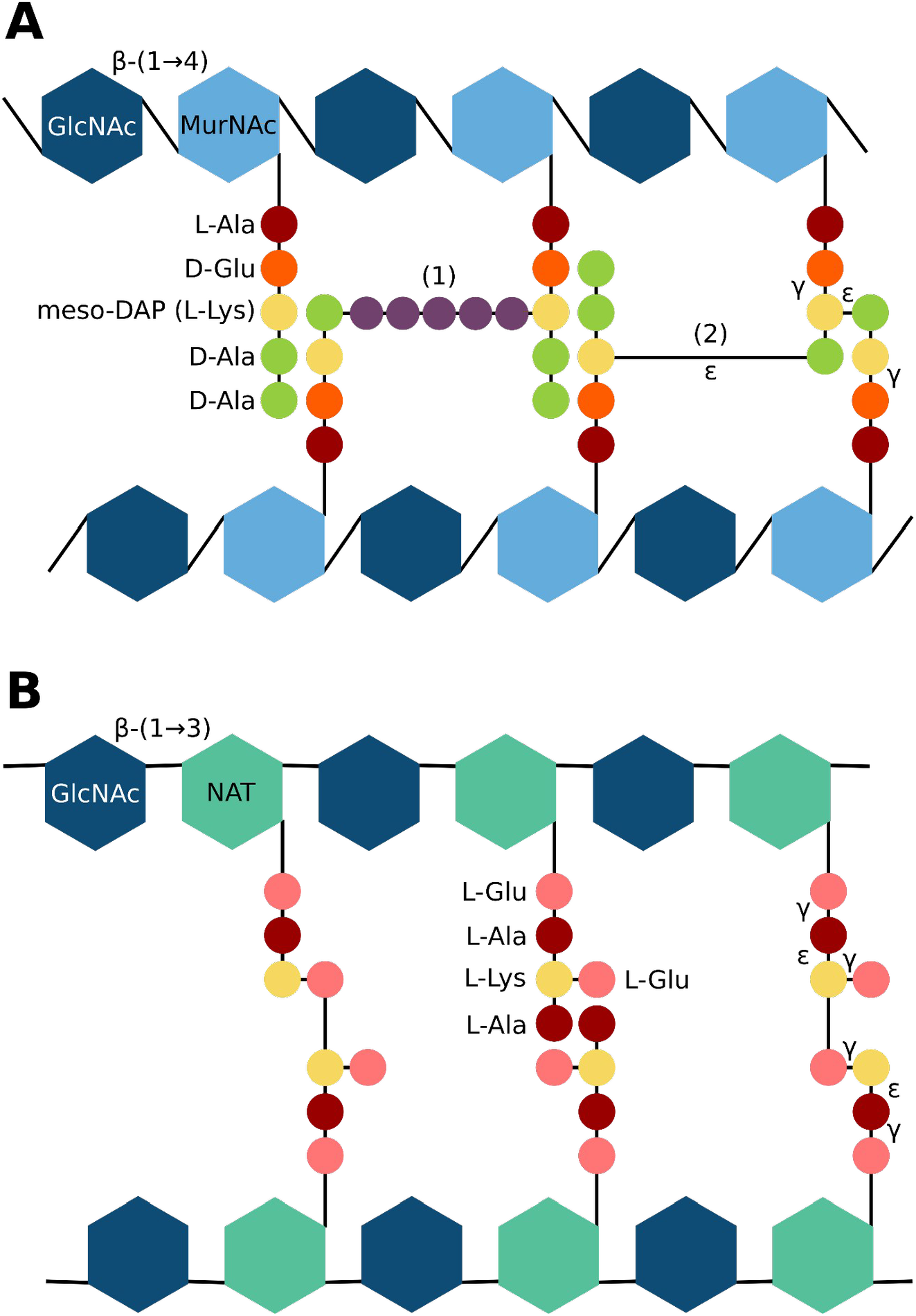
Structure comparison of the bacterial peptidoglycan (PG) and the archaeal pseudomurein (PM). **(A)** The glycosidic chain of PG is composed of alternating N-acetylglucosamine (GlcNAc) and N-acetylmuramic acid (MurNAc) units linked by a β-(1→4) bond. In most bacterial species, the pentapeptide attached to MurNAc is composed of L-alanine (L-Ala), D-glutamic acid (D-Glu), meso-diaminopimelic acid (meso-DAP; in *Escherichia coli*) or L-lysine (L-Lys; in *Staphylococcus aureus*), and two D-alanines (D-Ala). Interchain cross-linking usually occurs between the third amino acid (AA) of the first chain and the fourth AA of the second chain, accompanied by the loss of the D-Ala in position five. This cross-linking is either (1) indirect, through a pentaglycine bridge in *S. aureus*, or (2) direct in *E. coli*. **(B)** Instead of MurNAc, PM contains N-acetyl-L-talosaminuronic acid (NAT) units linked through β-(1→3) bonds to GlcNAc units. To NAT is attached a pentapeptide composed of L-Glu, L-Ala and L-Lys. Beyond the lack of D-AA, the archaeal pentapeptide bears more ε- and γ-peptide bonds than its bacterial counterpart.

Regarding PG, it is so crucial for cell survival and growth that even bacteria once thought to lack PG, like Planctomycetes or Chlamydiae, were actually shown to synthesize a thin layer of PG, notably during septal division (Liechti et al. 2014; Jeske et al. 2015; Packiam et al. 2015; van Teeseling et al. 2015; Liechti et al. 2016). Therefore, the proteins involved in PG biosynthesis have been extensively studied over the last years, in particular as potential targets for antimicrobial agents (Bhattacharjee 2016). Usually, many genes involved in PG biosynthesis lie in the *dcw* (division and cell-wall synthesis) gene cluster. The order of the genes within this cluster is relatively well conserved across the different bacterial lineages (Tamames 2001; Mingorance and Tamames 2004; Real and Henriques 2006), even if some species lack one or more PG biosynthesis genes in their genome (Pilhofer et al. 2008; Martínez-Torró et al. 2021). A recent reconstruction of the ancestral state of the *dcw* cluster showed that the last bacterial common ancestor (LBCA) had a complete *dcw* cluster, composed of 17 genes (Léonard et al. 2022).

Among the proteins encoded by *dcw* cluster genes, the four muramyl ligase enzymes, MurC, MurD, MurE, MurF, and the D-alanine--D-alanine ligase, Ddl, are critical for PG biosynthesis. The four muramyl ligase add, respectively and successively, L-Ala, D-Glu, meso-DAP (or L-Lys) and D-Ala-D-Ala to UDP-MurNAc, while Ddl binds two D-Ala to yield the D-Ala-D-Ala dipeptide (Pazos and Peters 2019; Egan et al. 2020). Inhibiting one of those genes leads to lysis of the bacterial cell (Zawadzke et al. 2008; Kouidmi et al. 2014). The muramyl ligases belong to the ATP-dependent Mur domain-containing family, which further includes four other enzymes: 1) MurT, which forms a complex with GatD to catalyze the amidation of D-Glu to D-glutamine (D-Gln) in *Staphylococcus* species (Münch et al. 2012; Nöldeke et al. 2018), 2) CapB, which plays a role in the formation of the poly-γ-glutamic acid capsule in *Bacillus* (Makino et al. 1989; Ashiuchi 2013; Hsueh et al. 2017), 3) cyanophycin synthetase (CphA), which catalyzes the polymerisation of L-arginine (L-Arg) and L-aspartate (L-Asp) into cyanophycin, a polymer that constitutes a nitrogen reserve in Cyanobacteria (Aboulmagd et al. 2001; Sharon et al. 2021), 4) folylpolyglutamate synthase (FPGS), which is responsible for the addition of polyglutamate to folate. The FPGS enzyme is found in the three domains of life: Archaea, Bacteria and Eukarya, but not in methanogenic archaea (Levin et al. 2004; Gorelova et al. 2019; Kordus and Baughn 2019; Kordus and Baughn 2019). Ddl is part of the ATP-grasp superfamily, including at least 21 groups of enzymes (Fawaz et al. 2011). Among those, the synthetase domain of carbamoylphosphate synthetase (CPS; Shi et al. 2018), CarB, is a well-studied enzyme that has been used to root the tree of life because it results from an internal gene duplication that occurred before the Last Universal Common Ancestor (LUCA) (Lawson et al. 1996; Philippe and Forterre 1999; Cammarano et al. 2002).

With the advances in genome sequencing, homologues of genes involved in PG biosynthesis, including muramyl ligases, have been identified in Methanopyrales and Methanobacteriales (Smith et al. 1997; Slesarev et al. 2002; Samuel et al. 2007; Leahy et al. 2010). Consequently, it was suggested that, despite the difference between the two biosynthetic pathways, the evolution of PG and PM are connected. More precisely, archaeal PM could have arisen from horizontal transfers (HGTs) of PG genes from Bacteria (Graham and Huse 2008; Subedi et al. 2021; Ithurbide et al. 2022). Last year, Subedi et al. 2021 re-investigated the PM biosynthetic pathway proposed by (Leahy et al. 2010) and resolved the first structure of an archaeal muramyl ligase, which they named pMurC, after its supposed homology with bacterial MurC. These recent studies have thus led to an increase in the number of candidate genes for PM biosynthesis. However, their function and exact role in the different steps of PM biosynthesis have still to be experimentally validated.

In the present work, we used a *de novo in-silico* approach to identify candidate genes for PM biosynthesis, characterized their functional domains using various prediction software and assessed the taxonomic distribution of their homologs in both bacterial and archaeal domains. We also investigated the evolutionary origins of PM by performing phylogenetic analyses of the Mur domain-containing family, the ATP-grasp superfamily and the MraY-like family using multiple variations of the taxon sampling and different AA substitution models. Our results reveal a bacterial origin of the four main archaeal muramyl ligases, which probably traces back to two HGT events in an ancestor of Methanopyrales and Methanobacteriales, followed by one or two rounds of gene duplication, depending on the considered gene. Moreover, *in silico* structural characterization of the muramyl ligases from two model archaea allowed us to tease apart their potential functions in PM biosynthesis.

## Results

### Collection of potential proteins for pseudomurein biosynthesis

For the identification of candidate genes for pseudomurein (PM) biosynthesis following an approach independent of already identified genes, we used the whole proteomes of ten archaeal organisms, corresponding to five PM-containing archaea (i.e., four Methanobacteriales and one Methanopyrales) and five non-PM Euryarchaeota (i.e., one Methanococcales, two representatives from different orders of Methanomicrobia, one Archaeoglobales and one Thermoplasmatales). The protein sequences of the ten archaeal assemblies were first clustered into 6,321 orthologous groups (OGs; clusters named from OG0000001 to OG0006321). A taxonomic filter allowed us to select 82 OGs specific to the PM-containing archaea, among which 26 OGs contained sequences of all five PM-containing archaea, whereas 56 OGs contained sequences of the only Methanopyrales and three Methanobacteriales (retained to maximize the sensitivity of our search). No OG was specific to the four Methanobacteriales. The paralogue-targeting approach (see Material and Methods) allowed us to identify 20 additional OGs. In parallel, eight OGs were selected using three pseudomurein-related HMM profiles downloaded from the NCBI CDD (Conserved Domain Database) (see Material and Methods). In total, 110 OGs were thus identified as candidates for PM biosynthesis (Fig. S1).

### Genetic environment of candidate proteins

Synteny analysis revealed that 22 out of 110 OGs are encoded by genes clustered in five regions of the genomes of PM-containing archaea, which we termed clusters A to E (Fig. S2). *In silico* functional analysis indicates (Table S1; sheet 1 to 3) that proteins of cluster A and B may be involved in PM biosynthesis while proteins of clusters C, D and E are probably not. Cluster C is a bidirectional cluster, where annotated proteins belong to different pathways. Indeed, OG0001177 and OG0001178 proteins are associated with pilus assembly proteins and/or surface proteins, while OG0001176 and OG0000094 can be associated with cell shape or gene regulation (the latter is not identified in our pipeline but its gene is always located downstream of the OG0001176 gene). Cluster D is related to nucleic acid metabolism or cellular signal transduction (Braun et al. 2021), whereas cluster E code for the four proteins that compose the methyl-coenzyme M reductase, which is implied in methane formation (Chen et al. 2020). Very recently, two potential clusters for PM biosynthesis were identified using bacterial proteins from PG biosynthesis as BLAST queries (Subedi et al. 2021). Those clusters correspond to our clusters A and B. Cluster A is composed of five genes: 1) OG0001014, which was experimentally characterized as the smallest CPS (Popa et al. 2012), 2) OG0001163, a type 4 glycosyltransferase homologue to MraY, 3) OG0001473, a Mur domain-containing protein, 4) OG0001162 and 5) OG0001472, two hypothetical proteins. Regarding cluster B, it is composed of three genes: 1) OG0001150, a Mur domain-containing protein, 2) OG0001147, a hypothetical protein and 3) OG0001146, a MobA-like NTP transferase domain-containing protein. In addition, two genes of Mur domain-containing proteins (i.e., OG0001148 and OG0001149) can be located either in cluster A or cluster B, and even outside any cluster, depending on the PM-containing species considered. Furthermore, another PM-specific gene (OG0000796, coding for a hypothetical protein) is located just downstream of the OG0001472 gene in the genome of *Methanopyrus sp. KOL6*, while a second one (OG0000169, coding for a Zn peptidase) is only three genes away from the OG0001146 gene in *Methanothermobacter thermautotrophicus str. Delta*. Based on the genetic environment of clusters A and B, we attempted to identify a conserved regulon for PM biosynthesis by phylogenetic footprinting (Cristianini and Hahn 2006; Anderssen et al. 2022). However, unlike in Bacteria (Anderssen et al. 2022), such analyses were unsuccessful on our archaeal dataset (Supplementary data).

Taking into account OG0000094, identified by its conserved localisation within cluster C, our pipeline recovered 23 syntenic genes (out of 111 OGs), of which half are likely to be involved in PM biosynthesis (Table 1). For clarity, in the following, the four Mur domain-containing proteins OG0001148, OG0001149, OG0001150 and OG0001473 will be arbitrary called Murα, Murß, Murγ and Murδ, respectively, without considering any specific homology with bacterial MurCDEF. Most of the proteins encoded in clusters A and B have no predicted signal peptide (SP) and are either cytoplasmic or transmembrane (TM) proteins. TM segment prediction was used as a complement to SP prediction. It allowed us to distinguish between cytoplasmic and transmembrane proteins, and revealed that only OG0000796, OG0001163 and OG0001472 are TM proteins. OG0000169 and OG0001162 feature a Sec SP and are thus the only exported proteins of these gene clusters. In PM-containing archaea, the synteny of the two genes of OG0001472 and murδ is highly conserved. However, in *Methanothermobacter thermautotrophicus str. Delta*, both genes were annotated as pseudogenes and thus not predicted as proteins.

**Table 1.** Overview of the proteins identified in our search for genes involved in PM biosynthesis. Orthologous Groups (OGs) composing the identified gene clusters, named clusters A to E are listed. For each OG, there is the functional prediction of InterProScan (if any), the predicted signal peptide type (SP) and the number of predicted transmembrane (TM) segments (0 = cytoplasmic, 1 = monotopic, >1 = polytopic).

| Cluster | Orthologous Groups | InterProScan Prediction | Signal peptide | # TM |
| --- | --- | --- | --- | --- |
| A | OG0001014 | CPS | Other | 0 |
|  | OG0001163 | MraY-like | Other | >1 |
| | OG0001473 | Muramyl ligase (= Murδ) | Other | 0 |
|  | OG0001162 | / | Sec | 0 |
|  | OG0001472 | / | Other | 1 |
|  | OG0000796 | / | Other | >1 |
| B | OG0001150 | Muramyl ligase (= Murγ) | Other | 0 |
|  | OG0001147 | / | Other | 0 |
|  | OG0001146 | MobA-like NTP transferase domain | Other | 0 |
|  | OG0000169 | Zn peptidase | Sec | 0 |
| A-B | OG0001148 | Muramyl ligase (= Murα) | Other | 0 |
| | OG0001149 | Muramyl ligase (= Murβ) | Other | 0 |
| C | OG0001210 | Aminotransferases class-I pyridoxal-phosphate attachment site | Other | 0 |
|  | OG0000094 | MreB/DnaK-like | Other | 0 |
|  | OG0001176 | Coiled coil protein | Other | 0 |
|  | OG0001177 | Flp pilus assembly protein<br>RcpC/CpaB | Other | 1 |
|  | OG0001178 | Sortase E | Other | >1 |
| D | OG0001213 | Zc3h12a-like Ribonuclease NYN<br>domain | Other | 0 |
|  | OG0001214 | Nucleotide cyclase | Other | >1 |
| E | OG0000266 | Methyl-coenzyme M reductase, beta<br>subunit | Other | 0 |
|  | OG0000231 | Methyl-coenzyme M reductase<br>operon protein D | Other | 0 |
|  | OG0000230 | Methyl-coenzyme M reductase,<br>gamma subunit | Other | 0 |
|  | OG0000229 | Methyl-coenzyme M reductase,<br>alpha subunit | Other | 0 |

### Taxonomic distribution of candidate proteins and their homologues

To ensure the completeness of the selected OGs, we looked for corresponding pseudogenes or mispredicted proteins in the genomes of the five PM-containing archaea (see Material and Methods). After completing the OGs, we retained only those containing protein sequences from all five PM-containing archaea, decreasing the number of OGs from 111 to 49. Interestingly, no OG from the five syntenic regions was discarded. Similarity searches in three local databases showed that 15 OGs are widespread (though not universal) among Bacteria and Archaea, 9 OGs have homologues only in bacteria, while 25 OGs are exclusive to archaea, among which 15 to PM-containing archaea (Fig. 2; for details see Table S2). In clusters A and B, which likely encode proteins involved in PM biosynthesis, 6 OGs are exclusive to Methanopyrales and Methanobacteriales whereas 7 OGs share homology with bacterial proteins. We also noticed that our HMM profiles of the four muramyl ligases (i.e., Murα, Murß, Murγ and Murδ) recovered a common set of sequences, indicating that Murαßγδ are specifically related. According to this taxonomic distribution, we further investigated the origin of CPS, the MraY-like and the four muramyl ligases Murαßγδ. The MobA-like NTP transferase, OG0001146, was not considered for phylogenetic analysis because, compared to the aforementioned proteins, no homologous protein was identified in the representative bacterial database (nor for OG0001215 and OG0000138). However, some bacterial homologues were identified when we determined the taxonomic distribution of the 49 OGs using the (much larger) prokaryotic database.

**Figure 2.**
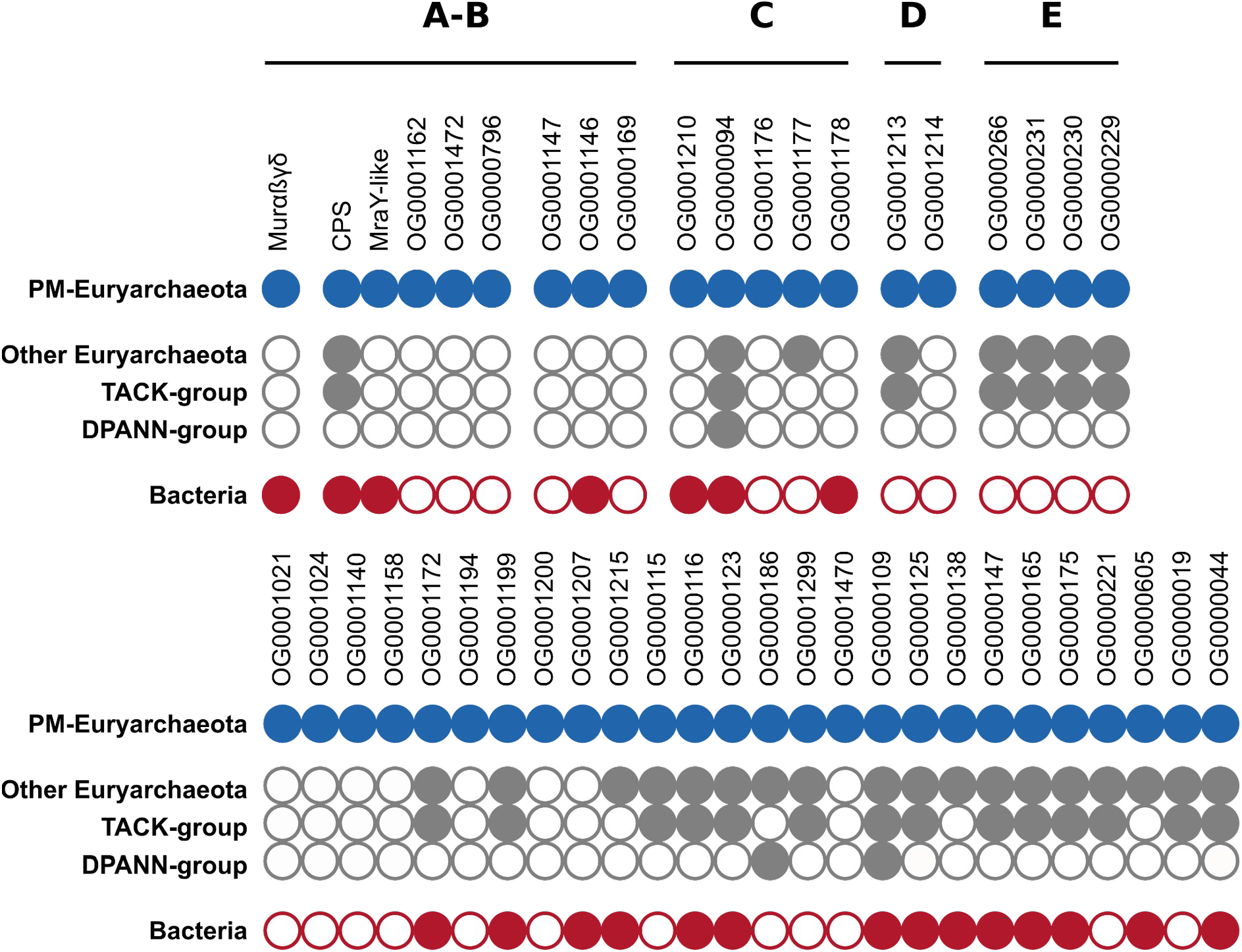
Taxonomic distribution patterns of the 49 retained orthologous groups (OGs). The four OGs OG0001148, OG0001149, OG0001150 and OG0001473 are considered together and referred to as Murαßγδ, OG0001014 is referred to as CPS and OG0001163 as MraY-like. Black lines delineate gene clusters in the genomes of PM-containing archaea (clusters A to E). Full circle = gene present in the taxonomic group; empty circle = gene absent from the taxonomic group.

### Phylogenetic trees

#### ATP-grasp superfamily

The CPS from the cluster A of PM-containing archaea, as well as the Ddl from the *dcw* cluster of bacteria, are member proteins of the ATP-grasp superfamily. Due to the large number of protein functions and architectures within the ATP-grasp superfamily (Fawaz et al. 2011), we focused our phylogenetic analyses on the ATP-grasp domain. Furthermore, we wanted to investigate whether CPS is closely related to Ddl (through HGT for instance). Thus, we excluded eukaryotic ATP-grasp proteins from our analyses. In the local databases, we identified 8,013 unique protein sequences containing at least one ATP-grasp domain, which are distributed across 1387 prokaryotic organisms. ATP-grasp domains were spliced out of full-length proteins, yielding a total of 12,074 domain sequences, then sequence deduplication led to 2344 sequences from which 149 highly divergent sequences were removed. Annotation showed that 1788 domain sequences correspond to 17 members of the ATP-grasp superfamily, while 406 sequences have no similarity with reference ATP-grasp sequences (see Material and Methods). We also observed that PyC, PccA and AccC reference sequences annotate sequences belonging to the same monophyletic group. These three enzymes use hydrogenocarbonate as a substrate (Diesterhaft and Freese 1973; Shen et al. 2006; Hou et al. 2015), which could explain the phylogenetic proximity of their ATP-grasp domain sequences. Accordingly, we decided to indistinctly tag the whole group with the three annotations. A similar observation and decision were made for PurK and PurT proteins, though the former uses hydrogenocarbonate as its substrate, while the latter uses formate (Mueller et al. 1994; Marolewski et al. 1997).

Due to an internal gene duplication that occurred before LUCA (Lawson et al. 1996; Philippe and Forterre 1999; Cammarano et al. 2002), the seven phylogenetic trees (see Material and Methods) were rooted on CarB, the monophyly of which is supported by high statistical values. Despite a low topology conservation between the different evolutionary models and number of tree search iterations, some recurring patterns can be observed (Fig. 3 and Fig S3 to S8). RimK ATP-grasp domain sequences are always paraphyletic, due to the inclusion of GshB, GshAB and CphA, the latter two clustering into a smaller clan. The monophyly of Acetate--CoA ligases AcD (Musfeldt and Schönheit 2002) is maximally supported and a long branch is present at the base of the group. Except for the C40 model (Fig S5 and S6), AcD forms a clan with the Succinate--CoA ligase SucC (Joyce et al. 1999). The position of the other members of the ATP-grasp superfamily is much more elusive. For example, Pur2 (Cheng et al. 1990) emerges somewhat alone in the LG4X tree (Fig. 3), whereas it forms a clan with either AcD and SucC in the four C20 and C60 trees (Fig S3-4 and S7-8) or only with SucC in the two C40 trees (Fig S5 and S6). Similarly, albeit Ddl and CPS branch together in one C20 tree with a branch support of 63 (Fig S3), their respective positions within the ATP-grasp superfamily are unstable (Fig 3 and Fig S3 to S8). Therefore, there is no strong phylogenetic evidence for a specific relationship between the Ddl and CPS proteins. In contrast, CPS is never close to CarB, which is at odds with the less extensive phylogenetic analyses of Popa et al. 2012.

**Figure 3.**
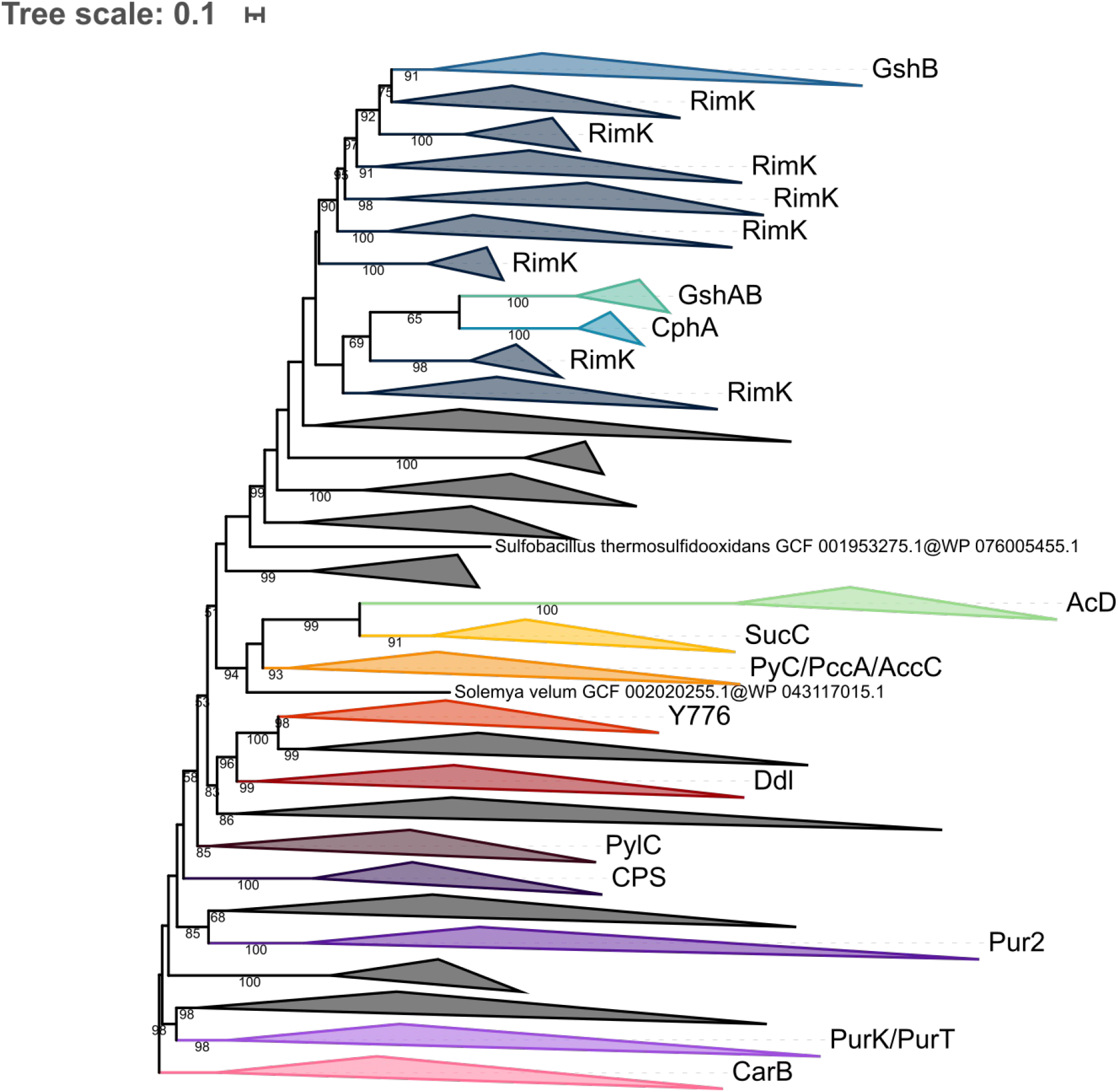
Phylogenetic tree of the ATP-grasp superfamily rooted on CarB. The tree was inferred from a matrix of 2,194 sequences x 180 unambiguously aligned AAs using IQ-TREE under the LG4X+R4 model. Tree visualization was performed using iTOL. Bootstrap support values are shown if greater or equal to 50. Branches were collapsed on sequence annotation based on reference sequences. Black collapsed branches correspond to unannotated sequences.

#### MraY-like family

Homology searches revealed that the bacterial homologue of OG0001163 is the glycosyltransferase 4 (GT4) MraY. According to the NCBI CDD (Lu et al. 2020), MraY is part of the MraY-like family, which further includes WecA (Amer and Valvano 2001), WbpL (Campbell et al. 1997; Price and Momany 2005), and eukaryotic and archaeal GPT (Dal Nogare et al. 1998). In addition to the MraY-like OG0001163, our pipeline has highlighted transmembrane proteins in OG0001207 (Fig. 2), for which the only bacterial homologue also has a MraY/WecA-like GT4 domain. Therefore, we decided to add the sequences of OG0001207 to the phylogenetic analysis of the MraY-like family. Although only one sequence similar to OG0001207 had been identified in the bacterial database, 62 additional bacterial OG0001207 homologues were identified in the (larger) prokaryotic database. According to the study of Lupo et al. 2021, none of the genomes coding for those protein sequences are considered as contaminated, which suggests that OG0001207 homologues genuinely exist in these bacteria. Overall, a total of 1267 sequences from the MraY-like family were identified in our databases, corresponding to 1071 unique sequences. Interestingly, 773 sequences among 1267 were identified by two or more HMM profiles of the individual members of the MraY-like family. During the annotation pipeline, six bacterial sequences remained unannotated due to their ambiguous position within the preliminary guide tree (see Material and Methods). Moreover, reference sequences of WecA and WbpL annotated putative sequences from the same monophyletic group and thus, the whole group was considered as WecA/WbpL.

Due to this non-universal taxonomic distribution and lack of an ancestral gene that could be present in the genome of LUCA, the three MraY-like family trees (see Material and Methods) were left unrooted. Phylogenetic analysis showed that each of the five members of the MraY-like family are monophyletic and all supported by high bootstrap values. Moreover, MraY formed a clan with WecA/WbpL while GPT formed a clan with OG0001163 and OG0001207 (Fig. 4). Those results are similar for the three evolutionary models LG4X, C20 and C40. Regarding the six unannotated sequences, the sequence of *Syntrophaceticus schinkii* is always basal to OG0001207, whereas the group composed of two sequences of *Ruminococcaceae* sp. and two sequences of *Treponema* sp. is always basal to MraY. The last sequence from *Ruminococcus* sp. is basal to MraY in the LG4X tree, while it is basal to WecA/WbpL in the C20 and C40 trees (Fig. S9 and S10). Taxonomic analysis revealed that MraY and WecA/WbpL are exclusive to bacteria, while GPT is only found in archaea. Regarding OG0001163, it is exclusive to PM-containing archaea, as would be OG0001207, ignoring the few exceptions discussed above.

**Figure 4.**
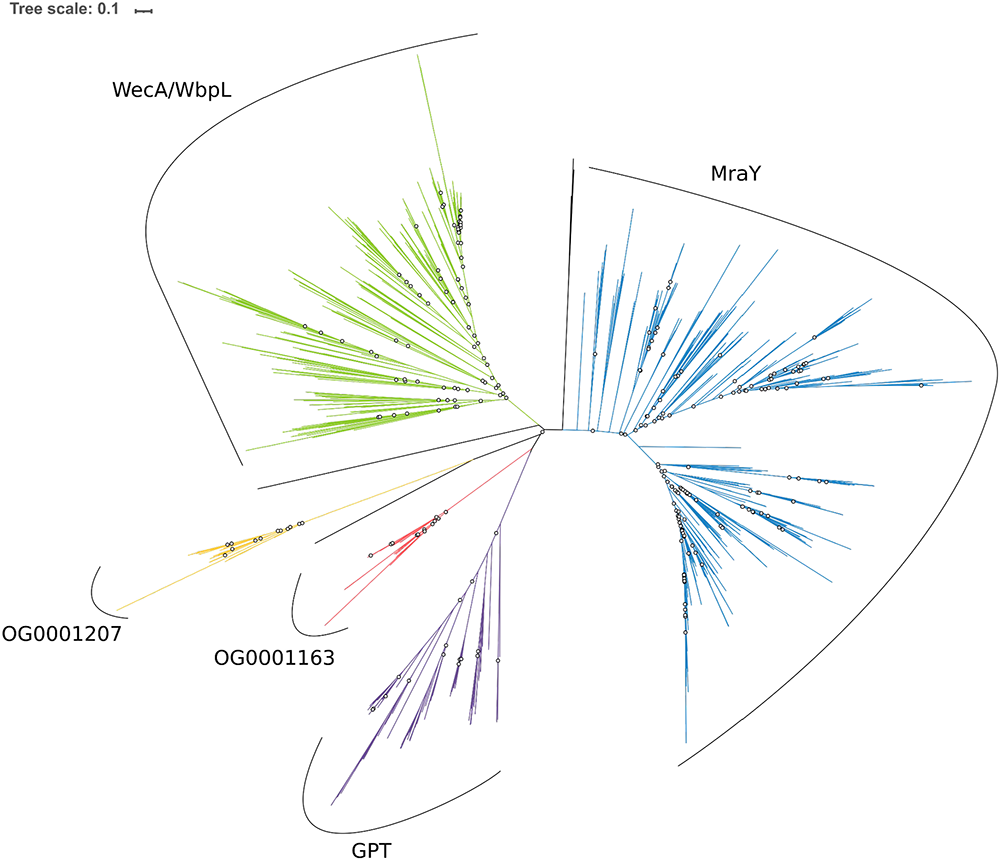
Unrooted phylogenetic tree of the MraY-like family. The tree was constructed from a matrix of 1,070 sequences x 408 unambiguously aligned AAs using IQ-TREE under the C40+G4 model. Open circles correspond to bootstrap support values under 90. Blue sequences correspond to a MraY annotation, green to WecA/WbpL, red to OG0001163 (MraY-like), yellow to OG0001207, purple to GPT, and black to unannotated bacterial sequences.

#### Mur domain-containing family

Homology searches allowed us to identify 3398 unique sequences distributed across 755 prokaryotic organisms. These sequences correspond to 12 members of the Mur domain-containing family, which are the four bacterial MurCDEF, the four archaeal Murαßγδ, MurT, CapB, CphA and FPGS. Taxonomic distribution within each member protein group revealed that MurCDEF and CphA are specific to bacteria, Murαßγδ are specific to PM-containing archaea, while MurT, CapB and FPGS are found both in Bacteria and Archaea, albeit not universally. According to the function and ubiquity of FPGS, we assumed that a FPGS protein was already present in LUCA, and trees were rooted on the corresponding clan. The phylogenetic trees, inferred with three models from a matrix of 3407 sequences x 550 AAs including the 12 members of the Mur domain-containing family, showed that each member group is monophyletic and supported by high statistical values (bootstrap values around 100; Fig S11 to S13), except for the long-branched sequence of *Francisella noatunensis*, tagged as MurE, which is positioned basal to the CphA clan (except in the C20 tree). In spite of the solid monophyly of each Mur domain-containing family member, the recovered relationships between these members (i.e., the topology of the family tree) depend on the evolutionary model (LG4X, C20 or C40). We made the same observation for phylogenetic reconstructions based on a smaller matrix restricted to the most conserved AAs over the full-length sequence (3386 sequences x 228 AAs) (Fig. S14 to S16).

In order to investigate the orthology relationships between the four bacterial muramyl ligases MurCDEF and their uncharacterized archaeal homologues Murαßγδ, we performed phylogenetic analyses using only one out of four potential outgroups among MurT, CapB, CphA and FPGS, under the three models (Fig. 5a and Fig S17 to S27). In these trees, Murα and Murß always group together, and further form a clan with Murγ and MurD in 11 trees out of 12. Murδ groups with MurC in eight of the single-outgroup trees. Furthermore, MurαßγD and MurδC form a clan in five trees, while this larger clan further includes MurT in the three trees where the latter is present. Interestingly, the sequence of *Francisella noatunensis*, tagged as MurE, groups with CphA instead of MurE when CphA is considered during phylogenetic inference. In the CapB and CphA outgroup trees computed with the C40 model, Murδ branches inside the MurE clan, within Firmicutes. Even though such an alternative relationship would fit the structure of *Methanothermus fervidus* Murδ (PDB codes 6VR8 and 7JT8), described as a ‘type E peptide ligase’ (Subedi et al. 2022), the analysis of the two matrices under the more sophisticated PMSF LG+C60+G4 model (Fig S28 and S29) did not return that topology, and instead supported the first solution. Besides, two phylogenetic trees focusing on indels, with FPGS as the only outgroup (see Material and Methods), tend to confirm the first topology too (Fig 5b and S30). Indeed, when using a binary encoding, we also observe a clan formed by MurαßγD and MurδC, which is supported by a bootstrap value of 100, while MurE and MurF are paraphyletic. However, in those indel trees, Murß forms a clan with Murγ rather than Murα.

**Figure 5.**
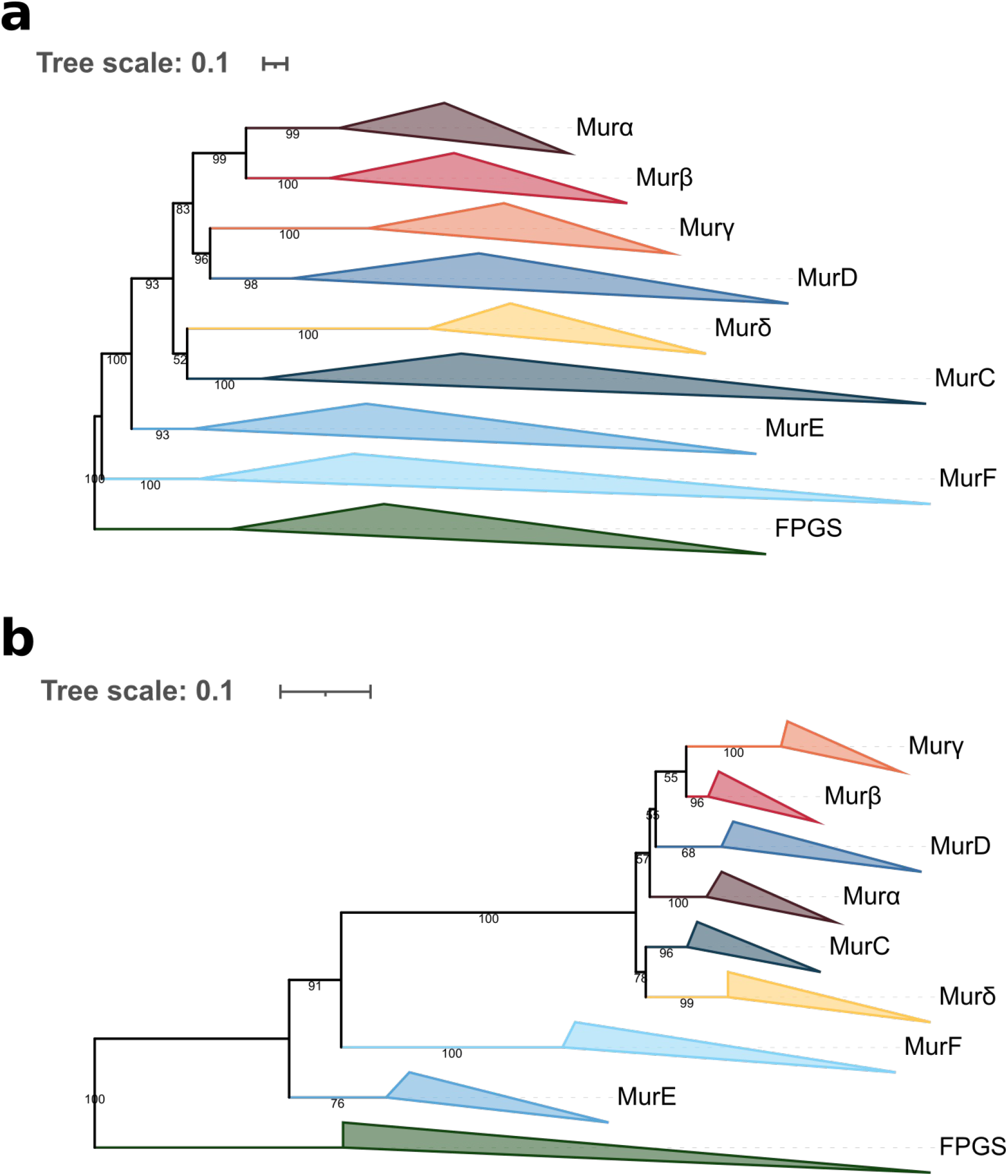
Phylogenetic trees of the Mur domain-containing family rooted on FPGS. (**a**) The tree was inferred from a matrix of 3,046 sequences x 543 unambiguously aligned AAs using IQ-TREE under the C40+G4 model. (**b**) “Indels” tree inferred from a matrix of 2997 sequences x 2243 unambiguously aligned AAs using RAxML under the BINGAMMAX model. Tree visualization was performed using iTOL. Bootstrap support values are shown if greater or equal to 50. Branches were collapsed on sequence annotation.

In parallel, jackknife support values from species resampling analyses (Table 2; see Table S3 for complete results and Material and Methods for details) confirmed the monophyly of each of MurC, MurD, Murα, Murß, Murγ, Murδ, CapB and FolC with jackknife support ranging between 99.7% and 100% under the three evolutionary models. Support for MurE, MurF and CphA is slightly lower and lies between 89.5 and 95.7%, whereas support for MurT is really low, with values ranging from 37.5 to 51.1% (Table 2 and Table S3). ASTRAL trees (Fig. S32 to S34) showed that the sequences of *Francisella noatunensis* (tagged as MurE) and *Solemya velum* gill symbiont (tagged as MurF) both group with CphA, which explains the lower jackknife support for the latter. When these two sequences are instead considered as belonging to CphA, support increases to 100% under the three evolutionary models. Support for MurE and MurF also increases (Table 2), which suggests that both sequences were mistagged by the annotation pipeline and rather are (divergent) CphA proteins. Furthermore, LG4X and C40 species trees revealed that MurT is polyphyletic and split into two distinct clans: 1) a large one composed of bacterial and Methanobacteriales sequences, and 2) a smaller one composed of sequences of Methanopyrales and Methanobacteriales, which we termed MurT-like. Indeed, support for MurT increases to 99.7% when MurT-like sequences are considered as a separate clan (Table S3). ASTRAL trees (Fig. S32 to S34) also confirmed the relationships between the eight muramyl ligases observed in the single-outgroup trees, even if those are blurred by the unstable positions of MurT and MurT-like. Murα and Murß are clustered in the three trees with a jackknife support of 86%, 71.3% and 66.7%, under LG4X, C20 and C40 models, respectively (Table 2). Regarding Murγ, it groups with MurT-like in LG4X (jackknife support of 39.0%) and C40 (38.9%) trees, which further form a clan with MurD (27.9% and 31.3%), whereas Murγ forms a clan with only MurD in the C20 tree (37.3%). Moreover, Murδ and MurC form a clan in the LG4X ASTRAL tree (47.5%), but are paraphyletic in the C20 (30.9%) and C40 (29.6%) trees. Murαßγ, MurD and MurT-like are grouped in the LG4X (27.5%) and C40 (15.9%) ASTRAL trees. In addition, MurαßγD, MurδC, MurT and MurT-like are grouped in the C20 (22.6%) and C40 (29.7%) trees. Symmetrically, these analyses revealed that MurE forms a clan either with MurF (30.7%, 16.4% and 14.7%) or CphA (36.8%, 34.1% and 30.6%) (Tables 2 and S3).

**Table 2.** Jackknife support values computed from the 1000 replicates of species resampling under three phylogenetic models: LG4X+R4, C20+G4 and C40+G4. Specific clans are shown if the support value reaches 200‰ in at least one of the three models. Here, the two misclassified sequences of MurE and MurF are considered as CphA sequences. For complete results, see Table S3.

| Clan | Support value (‰) |  |  |
| --- | --- | --- | --- |
|  | LG4X | C20 | C40 |
| CapB | 1000 | 1000 | 1000 |
| CphA | 1000 | 1000 | 1000 |
| FPGS | 1000 | 998 | 997 |
| MurC | 1000 | 1000 | 1000 |
| MurD | 1000 | 1000 | 1000 |
| MurE | 993 | 930 | 935 |
| MurF | 990 | 998 | 994 |
| MurT | 357 | 511 | 510 |
| Murα | 1000 | 1000 | 1000 |
| Murβ | 1000 | 1000 | 1000 |
| Murγ | 1000 | 1000 | 999 |
| Murδ | 1000 | 1000 | 1000 |
| α-β | 860 | 713 | 667 |
| D-γ | 379 | 373 | 356 |
| C-δ | 475 | 309 | 296 |
| E-F | 307 | 164 | 147 |
| CapB-FPGS | 269 | 621 | 655 |
| CapB-δ | 208 | 108 | 163 |
| CphA-FPGS | 322 | 109 | 110 |
| CphA-E | 368 | 341 | 306 |
| T-γ | 87 | 211 | 261 |
| CphA-E-F | 242 | 141 | 145 |
| CphA-FPGS-E | 258 | 72 | 67 |
| CphA-FPGS-E-F | 243 | 84 | 70 |
| D-T-α-β-γ | 136 | 159 | 208 |
| CapB-CphA-FPGS-C-E-F-δ | 136 | 159 | 208 |
| CapB-CphA-FPGS-E-F-δ | 188 | 324 | 404 |
| C-D-T-α-β-γ | 188 | 324 | 404 |
| C-D-T-α-β-γ-δ | 94 | 226 | 297 |
| CapB-CphA-FPGS-E-F | 94 | 226 | 297 |
| C-D-E-F-T-α-β-γ-δ | 58 | 189 | 258 |
| CapB-CphA-FPGS | 58 | 189 | 258 |
| CapB-FPGS-δ | 143 | 204 | 270 |
| T-α-β | 115 | 200 | 143 |

Moreover, CapB appears to be closely related to FPGS in C20 and C40 ASTRAL trees, with a jackknife support of 62.1% and 65.5%, respectively. As expected, the clan formed by MurE, MurF, CphA, FPGS and CapB has the same jackknife support as its counterpart (MurαßγDδCTT-like) in C20 (22.6%) and C40 (29.7%) trees (Tables 2 and S3). Therefore, it appears that the primary sequences of MurEF proteins are quite distinct from the six other muramyl ligases MurαßγδCD.

Overall, our analyses showed that neither MurT nor CphA should be considered as an outgroup for the Mur domain-containing family. Indeed, we observe that MurT sequences form either one or two (MurT + MurT-like) clans, which emerge from within the larger clan formed by the six muramyl ligases MurαßγδCD. In spite of the difficulty to determine the exact positions of MurT and MurT-like, topology and jackknife support tend to indicate that MurT sequences derive from the same ancestral gene as MurαßγδCD. In contrast to the other members of the Mur domain-containing family, CphA originates from the fusion of two functional domains: 1) an ATP-grasp domain at the N-terminal region (see ATP-grasp superfamily) and 2) the Mur ligase domain at the C-terminal region. This C-terminal region appears to be closely related to MurE and MurF in our phylogenetic inferences. Regarding CapB, species resampling showed that it is not related to the four bacterial muramyl ligases MurCDEF nor to the four archaeal muramyl ligases Murαßγδ, but more likely to FPGS (Table 2), thus indicating that it can be used as an outgroup to study the relationships between MurCDEF and Murαßγδ. However, unlike FPGS, CapB distribution is more restricted, the gene being found only in Gammaproteobacteria, Bacilli, Synergistetes, Halobacteria and a few Methanosarcinales and Korarchaota, according to our taxonomic analyses.

### 3D models of the archaeal Mur ligases

As the four archaeal muramyl ligases do not have straightforward orthology relationships with their four bacterial counterparts, phylogeny alone cannot help determining the origin of those enzymes. However, the 3D structure of proteins can be used as a complement to unravel the evolution of muramyl ligases (Chang et al. 2004; Illergård et al. 2009). The structures of “Murα” (PDB code 6VR7) and “Murδ” (PDB codes 6VR8 and 7JT8) from *Methanothermus fervidus* are available in the Protein Data Bank (PDB). This data was complemented by the 3D models of the Murαßγδ ligases from *M. fervidus* and *Methanothermus smithii* obtained with the AlphaFold software (Jumper et al. 2021). Importantly, the Murα and Murδ models were obtained using a version of the PDB reference database predating the release of the corresponding structures to assess the accuracy of AlphaFold on this type of protein. The overall quality of all the models is very good, with average pLDDT (predicted local-distance difference test) values of the best model superior to 90% and only a few loops with significantly lower pLDDT values (Fig. S35). For Murα from *M. fervidus*, the rms (root-mean-square) deviation between the crystallographic structure and the AlphaFold model calculated for the Cα is 2.1Å, while it is below 0.7Å when calculated separately for each of the three domains. For Murδ, these values are 2.43Å and below 1.0Å, respectively. This shows that the AlphaFold models are of very high accuracy for the individual domains but with some slight movements observed between the domains.

As the nature of the AAs transferred to the pseudomurein precursors depends on the structural features of the C-terminal domain of the various Mur ligases, the 3D structures of Murαßγδ were compared with those of MurCDEF to identify their respective role in PM biosynthesis. For Murδ, a clear homology was observed with the structure of the C-terminal domain of MurC (Mol Clifford D. et al. 2003), which adds L-Ala to MurNAc in Bacteria (Fig. 6a). The residues surrounding the L-Ala moiety are either strictly conserved (H198, R377, A459, H348 in MurC from *Haemophilus influenzae*) or substituted by an identical AA from a different structural element (R380 in *H. influenzae*) or substituted by residues with similar properties (H376 by a glutamine and Y346 by a phenylalanine). The AA added by Murδ to the archaeal PM peptide will therefore likely be an L-Ala as well, further strengthening the phylogenetic link identified between Murδ and MurC. However, as recently reported, the N-terminal domain of Murδ is more closely related to the corresponding MurE domain (both the primary and secondary structures) than to the MurC domain (Subedi et al. 2022).

**Figure 6.**
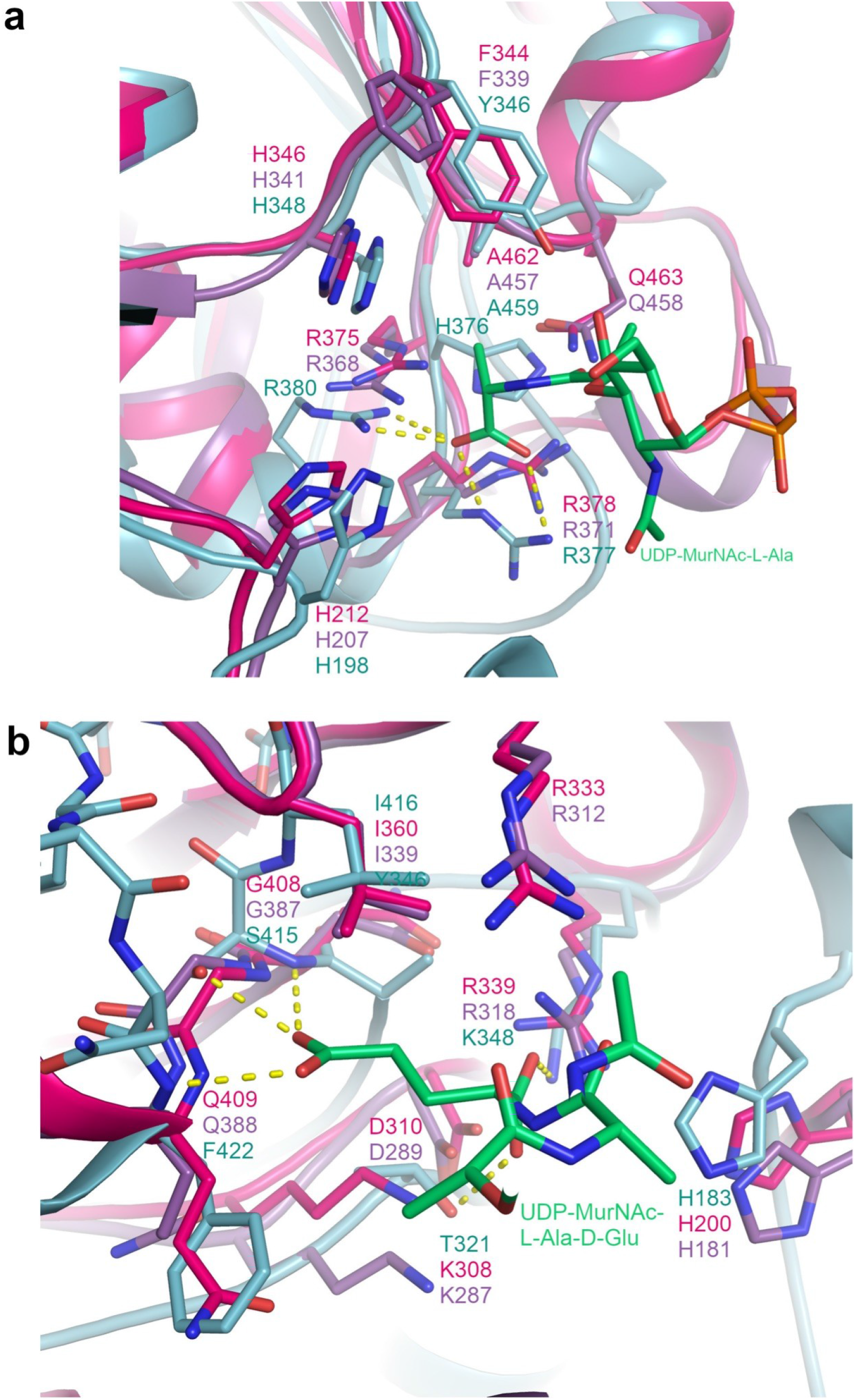
Identification of the amino acid recognized by the C-terminal domain of Murδ and Murγ. (**a**) Superimposition of the Murδ structure from *M. fervidus* (PDB code 6VR8; purple) with the AlphaFold model of Murδ from *M. smithii* (pink) and the MurC structure from *Haemophilus influenzae* (PDB code 1P3D; light cyan) in complex with UDP-MurNAc-L-Ala (green). H-bonds between the L-Ala moiety and MurC are shown as yellow dashed lines. (**b**) Superimposition of the AlphaFold models of Murγ from *M. fervidus* (purple) and *M. smithii* (pink) and the MurD structure from *E. coli* (PDB code 4UAG; light cyan) in complex with UDP-MurNAc-L-Ala-D-Glu (green). H-bonds between the D-Glu moiety and MurD are shown as yellow dashed lines.

A second significant match was observed between the structure of C-terminal domains of Murγ and MurD (Bertrand et al. 1999), which is responsible for the addition of D-Glu in Bacteria (Fig. 6b). The conservation is less strict in this case (only I416 of MurD from *E. coli* is conserved in Murγ), but the functionality of other AAs surrounding the D-Glu substrate is maintained. S415 and F422, which stabilize the γ-carboxylic acid through their backbone nitrogen and serine hydroxyl, are replaced by the backbone nitrogen of a glycine and a subsequent glutamine. In PM, the only AA with a carboxylic group away from the reaction center is the L-Glu added at the fifth position through its γ-carboxylic acid. This reaction must however involve a significant modification in the vicinity of the reaction center, as the functional groups of the stem peptide and AA added are inverted (bond between the γ carboxylic acid of L-Glu and ε amine of L-Lys at the third position of the peptide). In this context, it is therefore difficult to interpret the replacement of K348 and T321, which stabilize the α-carboxylic acid in MurD, by an arginine and a lysine, respectively, as well as the presence of an arginine and an aspartic acid (R312 and D289 in *M. fervidus*) close to the reaction center. While the ligation of L-Glu to the L-Lys in third position by Murγ is not fully validated by the comparison with MurD, it remains the most likely role of this enzyme.

For Murα and Murß, the comparison with the structure of the C-terminal domain of bacterial Mur enzymes did not reveal obvious similarities. However, in the Murα structure from *M. fervidus* and the model from *M. smithii*, two glutamic acids are conserved in the cavity usually accommodating the substrate, suggesting a role in the ligation of the L-Lys rather than the second L-Ala. This would leave Murß for the addition of the other L-Ala of the PM stem peptide, but it is difficult to verify because the two AlphaFold models of Murß analyzed are not congruent in this region.

## Discussion

Our phylogenetic analyses of the Mur domain-containing family show that each member of the Mur family is monophyletic. However, the relationships between those members are hard to establish owing to the low phylogenetic signal within the family and because phylogenetic artifacts, such as LBA (Gouy et al. 2015), probably affect phylogenetic reconstruction, especially for the trees including all non-Mur “outgroups”. Indeed, compared to MurCDEF, archaeal muramyl ligases (here termed Murαßγδ) are characterized by very long branches, and particularly Murδ, which has experienced more than one substitution per site since its probable separation from MurC. When focussing on Mur trees with only one outgroup, the topology is quite robust to different evolutionary models and species resampling within each member of the Mur domain-containing family. In this topology, MurD forms a clan with Murα+Murß+Murγ, MurC a clan with Murδ, and MurE a clan with MurF, a result that is also compatible with unrooted trees devoid of any outgroup (Fig S36 to S38). Moreover, structural analyses of the C-terminal domain of the four archaeal muramyl ligases allowed us to assign them a putative function in PM biosynthesis (Fig. 7). Indeed, due to some similarities between MurC and Murδ and between MurD and Murγ, we assume that Murδ adds one of the two L-Ala and Murγ adds L-Glu to the stem peptide. Although there are no obvious similarities between Murα and Murß and bacterial muramyl ligases, some clues suggest that Murα is responsible for the addition of L-Lys. Therefore, the second L-Ala of the stem peptide is probably added by Murß.

**Figure 7.**
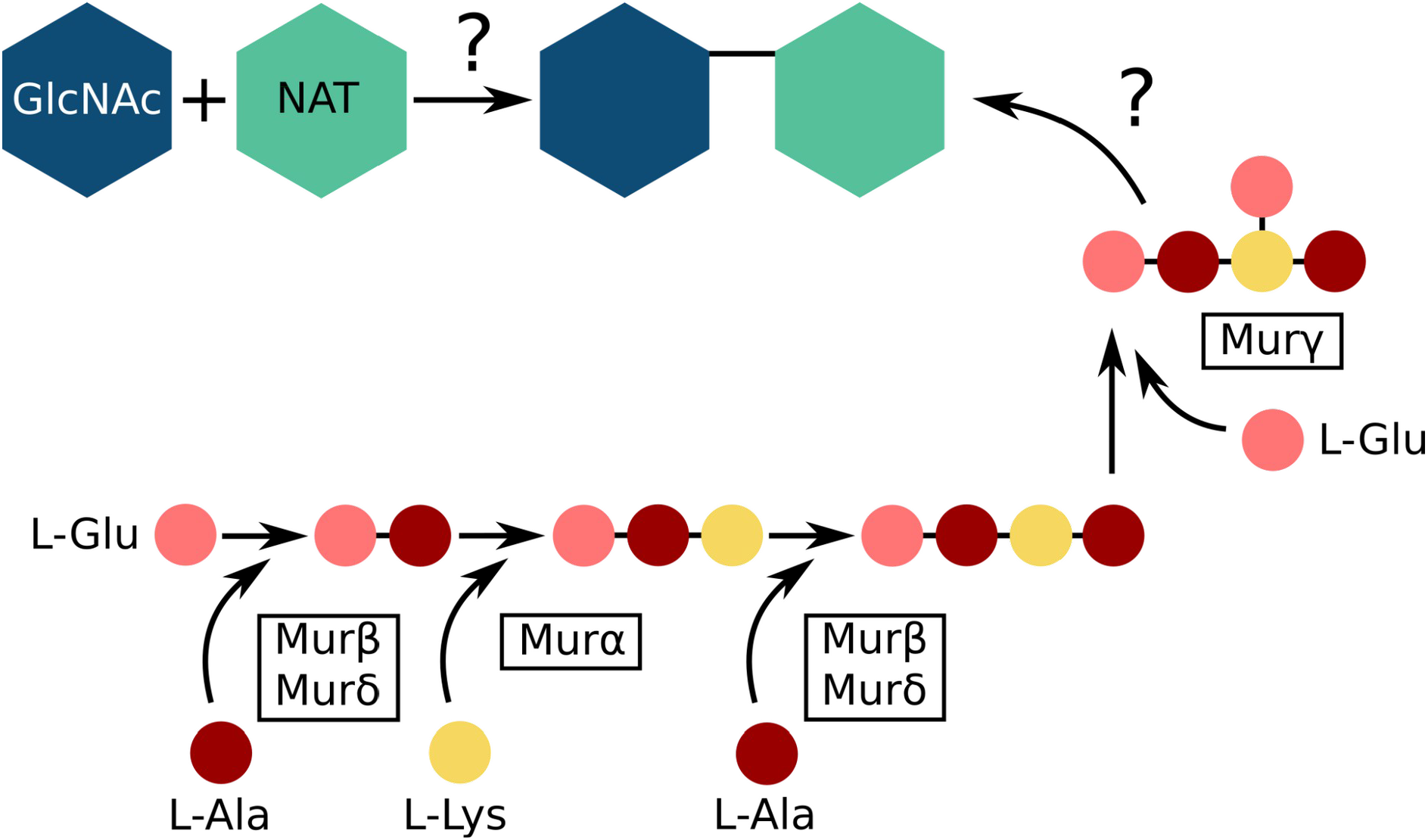
Putative functions proposed for the four archaeal muramyl ligases (Murαßγδ) based on 3D structure comparisons. The pathway presented here is the scenario proposed by Evamarie Hartmann, Helmut König and Uwe Kärcher (Hartmann and König 1990; König et al. 1993; Hartmann and König 1994). Although a specific function has been attributed to each archaeal muramyl ligase, we could not determine which one between Murβ and Murδ adds the L-Ala in position 2 and the L-Ala in position 4 of the stem peptide.

As previously stated, early analyses of their biosynthetic pathways have suggested that neither PG nor PM were a feature of LUCA (Scheffers and Pinho 2005; Albers and Meyer 2011; Subedi et al. 2021; Ithurbide et al. 2022). Therefore, LUCA probably did not possess the various muramyl ligases presently involved in cell-wall biosynthesis. However, FPGS is found in the three domains of life (Levin et al. 2004; Gorelova et al. 2019; Kordus and Baughn 2019), indicating that the gene was already part of the genome of LUCA. Thus, muramyl ligases emerged in Bacteria from a duplication of an ancestral version of FPGS and then were transferred to the other domain. In our phylogenetic trees, archaeal muramyl ligases (Murαßγδ) never branch within bacterial muramyl ligases (MurCDEF), and those trees do not give clues about the direction of the transfers. However, this topology could also be an artifact due to fast-evolving sequences in archaeal species. This kind of artifact has already been reported, e.g., with plastidial genes in eukaryotes, which rarely branch within (and rather sister to) Cyanobacteria (Sato 2021), although the endosymbiotic origin of the plastid is widely accepted (Ponce-Toledo et al. 2019). Because the LBCA already possessed a complete *dcw* gene cluster (Léonard et al. 2022), and given that PM is restricted to Methanopyrales and Methanobacteriales (Meyer and Albers 2020), we propose a scenario for the evolution of archaeal muramyl ligases through HGT (Fig. 8).

**Figure 8.**
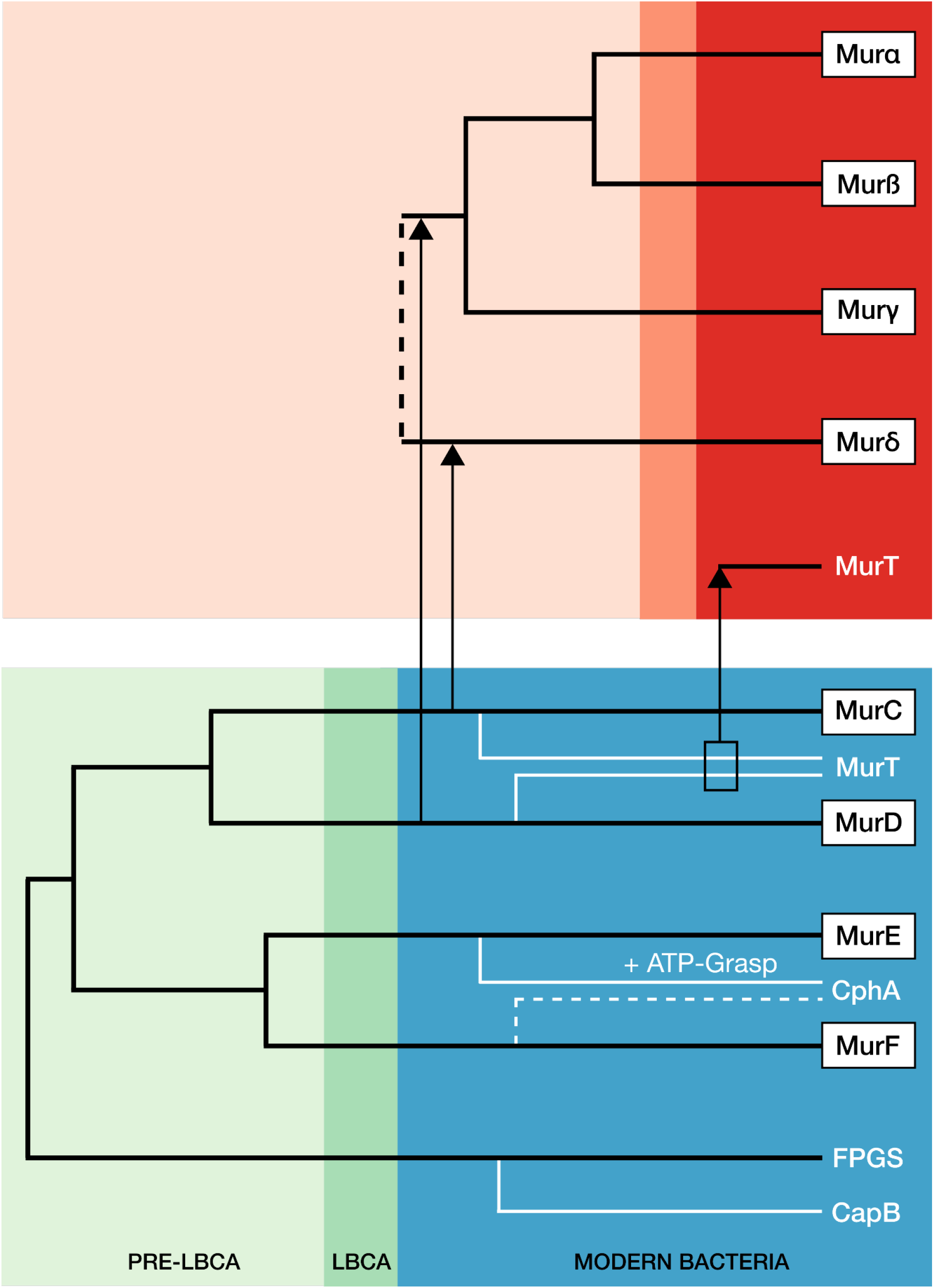
Proposed scenario for the duplication events and horizontal gene transfers from Bacteria to Archaea having led to the extant organization of the Mur domain-containing family. In this figure, only one possible origin is represented for Murδ, the hypothesis where it stems from MurC.

In this scenario, the ancestral gene of *murCDEF* was duplicated a first time in the pre-LBCA lineage to yield the ancestral genes of *murCD* and *murEF*, followed by a second round of duplications, which led to the current four bacterial muramyl ligases. Some evidence indicates that the duplication of the *murEF* ancestral gene to yield *murE* and *murF* could have occurred later than the duplication of the *murCD* ancestral gene. In fact, *murE* and *murF* genes are always in tandem in the *dcw* cluster of most bacterial species, as well as in the reconstruction of the LBCA *dcw* cluster (Léonard et al. 2022), and can even be expressed as a single fusion protein MurE-MurF (Laddomada et al. 2019). Moreover, in the majority of our Mur domain-containing family trees, MurE and MurF have slightly shorter branches than those of MurC and MurD. Early after the diversification of the LBCA, the *murD* gene was transferred to the common ancestor of Methanopyrales and Methanobacteriales, then *murD* experienced two duplications that yielded murα, murß and murγ (our nomenclature). In addition, *M*urα, Murß and Murγ exhibit a 3D fold similar to MurC/MurD for each of its three domains except for the presence of insertions in some loops (Fig. S39). In contrast, there is strong evidence that *murα* and *murß* arose from a gene duplication. These two muramyl ligases group together in almost all phylogenetic reconstructions (in both rooted and unrooted trees) and, as for MurF and MurE, their genes are in tandem in the genome of the majority of PM-containing archaea. Moreover, some Methanobrevibacter and Methanothermobacter genomes (two genera of Methanobacteriales) code for a Murα-Murß fusion protein (Subedi et al. 2021). As for the first, older, duplication of *murD*, leading to *murγ* and the *murαß* ancestor, it is visible in unrooted trees, where *murα*, *murß* and *murγ* form a clan.

However, the origin of the *murδ* gene remains unclear: while most of the phylogenetic trees and conserved residues in the C-terminal domain associate Murδ with MurC, the 3D structure of the N-terminal domain suggests that Murδ is rather related to MurE (Subedi et al. 2022). This inconsistency between phylogeny and structure can be due to different phenomena that are still to be untangled. First, the phylogenetic models struggle to exactly position the Murδ clan, probably due to its long basal branch. In most of the cases, Murδ forms a clan with MurC, while two phylogenetic trees using a C40 model (Fig S28 and S29) show Murδ emerging from within the MurE clan. Second, one cannot exclude evolutionary convergence, where a *murC* gene was first transferred and then its 3D structure gradually shifted to a MurE-like fold, or conversely, a *murE* gene was transferred and its key AAs converged to a MurC-like sequence. Finally, a more complex scenario would be the transfer of both *murC* and *murE* genes, followed by their recombination at the domain level, leading to the current Murδ.

Species resampling allowed us to complete this scenario with the three remaining proteins from the Mur domain-containing family: MurT, CphA and CapB. Hence, our analyses showed that MurT is clearly related to the clan formed by MurαßγDδC, CphA related to the MurEF clan, while CapB appears close to the outgroup, FPGS. In contrast to FPGS and MurCDEF, which are ubiquitous in Bacteria, MurT, CphA and CapB have a patchy distribution. Thus, they have probably arisen in a specific lineage, followed by HGT, instead of being a feature of the LBCA. In such a context, we assume that MurT could be derived from MurC or MurD, while CphA would originate from the fusion of an ATP-grasp containing gene, similar to the Glutathione biosynthesis GshAB, and a MurE or MurF gene. Regarding CapB, its origin is less clear but, like FPGS, CapB uses L-Glu as a substrate (Hsueh et al. 2017; Gorelova et al. 2019). Therefore, CapB could have been recruited from a duplicated FPGS gene, which suggests that it was indeed a suitable outgroup to study the relationships among the eight muramyl ligases.

Further insight about the transfers between Bacteria and PM-containing archaea can be obtained from the phylogeny of MurT. Previously, MurT has been described in *Staphylococcus* spp*, Streptococcus pneumoniae* and *Mycobacterium tuberculosis* (Münch et al. 2012; Morlot et al. 2018; Nöldeke et al. 2018; Maitra et al. 2021). Our analyses revealed that MurT is not ubiquitous in Bacteria, being only found in Firmicutes, Actinobacteria, *Caldisericum exile* (Caldiserica) and *Thermobaculum terrenum* (Chloroflexi). We also identified homologues in Archaea, specifically in Methanopyrales and Methanobacteriales. Interestingly, almost all bacteria have one copy of the *murT* gene while some PM-containing archaea have two copies, which we named *murT* and *murT-like*. Surprisingly, Methanobacteriales can possess only MurT or only MurT-like or both, while the few available Methanopyrales solely have one MurT-like gene. Moreover, archaeal MurT sequences are monophyletic and emerge from within Firmicutes (as sometimes observed for Murδ; Fig S28 and S29), while bacterial MurT sequences are consequently paraphyletic. Regarding MurT-like, the clan is monophyletic and basal to the MurT clan. In genomes of *Staphylococcus* species, *murT* and *gatD* genes are clustered in an operon (Münch et al. 2012; Morlot et al. 2018). Methanobacteriales and bacterial species that harbor a MurT homolog also have a GatD homolog while no GatD homologs are found in archaeal species bearing only MurT-like. This pattern suggests that MurT and GatD genes were transferred together to Methanobacteriales from a Terrabacteria lineage, probably Firmicutes. According to the taxonomic distribution of archaeal MurT/GatD and Murαßγδ, we can assume that the gene transfers of *murT*/*gatD* and the two ancestor genes of murαßγδ both occurred before the diversification of PM-containing archaea. In contrast, the origin of the *murT-like* gene is enigmatic, even though one possible explanation would be a duplication of *murT* in the LCA of Methanopyrales and Methanobacteriales, followed by differential loss of either *murT*/*gatD* or murT-like in some recent lineages.

In any case, those scenarios assume that the LBCA is older than the LCA of Methanopyrales and Methanobacteriales. However, molecular dating of prokaryotes is challenging since there are only a few microbial fossils or traces for which a meaningful taxonomy was proposed. The oldest evidence for microbial life has been identified in the Nuvvuagittuq belt in Quebec, Canada, which is between 3.75 and 4.28 billion years old (Gy) (Dodd et al. 2017; Papineau et al. 2022). There are also Archean rocks from up to 3.5 Gy containing chemical traces of microbial methanogenesis and sulfate reduction (Shen et al. 2001; Ueno et al. 2006; Aoyama and Ueno 2018; Catling and Zahnle 2020; Mißbach et al. 2021), thereby indicating that methanogenesis could be one of the most ancient biochemical pathways. Moreover, methanogenesis is a metabolism specific to the archaeal lineage (Gribaldo et al. 2006; Sorokin et al. 2017; Spang and Ettema 2017; Drake and Reiners 2021). Regarding bacterial microfossils, only three are unambiguously identified, all affiliated with the cyanobacterial lineage, of which *Eoentophysalis*, the oldest one, has been described from 1.9 Gy stromatolites (Hofmann 1976). Most scientists agree on the idea that the Great Oxidation Event (GOE) that occurred 2.4 Gy ago was due to the rise of oxygenic photosynthesis by Cyanobacteria. Using the GOE and the cyanobacterial fossil record as constraints for molecular clocks, it has been estimated that the cyanobacterial lineage appeared slightly before the GOE, as reviewed in Demoulin et al. 2019. Two recent molecular clock studies used horizontal gene transfers between archaeal methanogens and the LCA of Cyanobacteria, along with the cyanobacterial fossil record and the GOE, to date the origin of euryarchaeotal methanogens. They estimate the divergence between Euryarchaeota and the TACK group to have occurred around 4.1 and 3.8 Gy ago. Within Euryarchaeota, the LCA of class I methanogens (CIM) and class II methanogens (CIIM; Bapteste et al. 2005) (i.e., Methanomicrobiales and Methanosarcinales) originated 3.66 Gy ago (Gribaldo et al. 2006; Wolfe and Fournier 2018). As we hypothesize above, murT/gatD and murαßγδ genes were transferred from one or more bacterial lineages to the ancestor of Methanopyrales and Methanobacteriales. In present-day microbial communities, methanogens and sulfate-reducing bacteria (e.g., Deltaproteobacteria or Firmicutes) share the same ecological niche and can live in syntrophy under certain conditions (Lin et al. 2006; Muyzer and Stams 2008; Ozuolmez et al. 2015; Zouch et al. 2017). Evidence of such associations between sulfur-reducing and methanogens organisms were also identified in geological fluid inclusions from 3.5 Gy ago (Mißbach et al. 2021). Therefore, gene transfers between sulfur-reducing bacteria and methanogenic archaea could have occurred in that kind of environment and led to the origin of PM * containing archaea.

In one of the two putative gene clusters for PM biosynthesis, there is an ATP-grasp domain-containing gene that is always located upstream of the *mraY-like* and *murδ* genes. Moreover, this ATP-grasp domain-containing gene is exclusive to Methanopyrales and Methanobacteriales species. Thus, it has been proposed that it is probably involved in PM biosynthesis (Subedi et al. 2021). However, this gene was previously experimentally characterized by Popa et al. 2012, who concluded that it was a small (actually the smallest) carbamoyl phosphate synthetase (CPS) closely related to the “true” CPS, CarB. Given its putative function in cell-wall biosynthesis and its restricted taxonomic distribution, we hypothesized that this small CPS was not related to CarB but to Ddl instead and, as the muramyl ligases, had been transferred from a bacterial lineage to PM-containing archaea. However, our extensive phylogenetic analyses of the ATP-grasp superfamily remained inconclusive about the origin of the small CPS. Indeed, in our seven trees, it never clusters with CarB, nor with Ddl (except in the C20 tree). Moreover, the whole group is supported by a long branch, which can explain the difficulty to position the small CPS (i.e., LBA artifact). Although our phylogeny of the small CPS is inconclusive, its genetic environment suggests that it is indeed involved in PM biosynthesis. Accordingly, we postulate that the reported CPS function of this enzyme might be non-specific. If so, its real function in PM biosynthesis still has to be experimentally determined.

Located right downstream of the ATP-grasp domain-containing gene, the *mraY-like* (OG0001163) gene codes for a transmembrane protein that shows homology with the bacterial MraY. However, this archaeal MraY-like does not appear to have evolved from the bacterial MraY (i.e., through HGT). Indeed, bacterial and archeal proteins are clearly separated in all unrooted phylogenetic trees, although archaeal monophyletic groups are characterized by long branches, especially OG0001207, which could lead to strong phylogenetic artifacts (i.e, LBA). In contrast to Mur domain-containing family and ATP-grasp superfamily trees, MraY-like family trees were left unrooted. In fact, none of the MraY-like family members is found in both Bacteria and Archaea. As shown in the Results section, MraY and WecA/WbpL are only present in bacterial species, GTP is ubiquitous to archaea while MraY-like and OG0001207 are exclusive to PM-containing archaea. In addition, WecA/WbpL is the only monophyletic group where some organisms bear two sequences, which could indicate that WecA and WbpL are two paralogues. The position of the monophyletic group composed of the four bacterial unannotated sequences revealed that they could be divergent MraY sequences. According to the taxonomic distribution of MraY, WecA/WbpL and GPT, we propose a scenario where an ancestral GT4 gene found in LUCA was vertically transmitted to both Archaea (GPT) and Bacteria (the ancestral gene of MraY and WecA/WbpL). The bacterial gene was then duplicated once to yield *mraY* and *wecA/WbpL*, and the latter experienced a second duplication in some bacterial species. Thus, GPT would be the orthologue of MraY and WecA/WbpL, while MraY and WecA/WbpL would be paralogous. For this phylogenetic analysis of the MraY-like family, we followed the family as defined in the NCBI CDD (https://www.ncbi.nlm.nih.gov/Structure/cdd/cddsrv.cgi?uid=264002) to increase the sequence sampling. In theory, it is possible that we undersampled the family. Indeed, the GT4 domain is also present in other proteins, like MurG (Mengin-Lecreulx et al. 1991; Laddomada et al. 2019), which are not part of the MraY-like family. A proper way to study the origin of the MraY-like family would be to infer a phylogenetic tree of the GT4 domain. However, such an analysis would be very time-consuming due to the large number of GT sequences (Lombard et al. 2014). For now, overlapping HMM search results starting from the different family members do not suggest any undersampling issue. Furthermore, although bacterial homologues of OG0001207 have a MraY/WecA-like GT4 domain, the long branch of the monophyletic group could indicate that OG0001207 and homologues are probably not part of the MraY-like family.

The current architecture of PG and PM are well-known, but it is clear that both polymers were different in their early evolutionary state, i.e., before acquisition and diversification of their respective muramyl ligases. However, inferring the ancestral states of PG and PM is almost impossible because those evolved in the stem branch of Bacteria or CIM Archaea, respectively, before the LCAs of extant organisms. As other Mur-ligase family proteins, like CapB, FPGS, MurT or CphA, bind AAs with an α-carboxylic acid group (i.e., aspartic acid and glutamic acid), we can speculate that the first muramyl ligase proteins were also associated with those AAs. Moreover, glutamic acid is one of the most abundant AAs in many organisms, and it participates in a wide array of metabolisms (Walker and van der Donk 2016).

Therefore, glutamic acid could be one of the first AAs to have been selected by muramyl ligases. In *Bacillus*, the complex formed by CapB, CapC, CapA and CapE recruits L-Glu or D-Glu to synthesize the poly-γ-glutamic acid capsule. This kind of cell wall has been suggested to occur in *Haloquadratum walsbyi*, based on genomic analyses. *H. walsbyi* is classified in Halobacteria, a class of Euryarchaeota characterized by a diverse variety of cell walls: S-layer, sulfated heteropolysaccharides, halomucin and a glutaminylglycan. The latter is composed of poly-γ-L-glutamate, to which are linked two types of oligosaccharides (Meyer and Albers 2020). Analyses showed that CapB is ubiquitous in Halobacteria, indicating that CapB could be involved in glutaminylglycan biosynthesis. Consequently, we suggest that this simpler cell wall could resemble the ancient forms of PG and/or PM.

## Material and Methods

### Data availability

Publicly available datasets, including all detailed YAML configuration files used with Forty-Two (Irisarri et al. 2017; Simion et al. 2017) and classify-ali.pl (D. Baurain; https://metacpan.org/dist/Bio-MUST-Core), and a detailed command line log file can be found here: https://doi.org/10.6084/m9.figshare.21641612.

### Protein sequence databases

Three local mirrors of NCBI RefSeq were used during this study: 1) an archaeal database composed of the 819 whole genomes that were available on March 7, 2019, 2) a bacterial database of 598 representative genomes selected by the ToRQuEMaDA pipeline (Léonard et al. 2021) and 3) a prokaryotic database of 80,490 genomes, already used in (Lupo et al. 2022). To assemble the bacterial database, ToRQuEMaDA was run in June 2018, according to a ‘direct’ strategy and using the following parameters: dist-metric set to JI (Jaccard Index), dist-threshold set to 0.86, clustering-mode set to ‘loose’, and pack size set to 200.

### Identification of candidate proteins for pseudomurein biosynthesis

Protein orthologous groups (OGs) were built from the conceptual translations of ten archaeal whole genomes using OrthoFinder v2.2.1 (Emms and Kelly 2015) with default parameters. These archaeal genomes correspond to five organisms having pseudomurein (PM) (GCF_000008645.1, GCF_000016525.1, GCF_000166095.1, GCF_002201915.1, GCF_900095295.1) and five without PM (GCF_000011185.1, GCF_000013445.1, GCF_000017165.1, GCF_000025285.1, GCF_000251105.1) and were downloaded from the NCBI RefSeq database on March 7th, 2019. Then, taxonomic filters were applied to the OGs using classify-ali.pl v0.212670 in order to select candidate proteins for PM biosynthesis. Hence, we first looked for OGs with protein sequences from all five PM-containing archaea or from one Methanopyrales and three Methanobacteriales or from four Methanobacteriales. To identify OGs corresponding to a widespread gene that would also include a paralogue potentially specific to PM-containing archaea, we used the same taxonomic criteria but set the ‘min_copy_mean’ option to 1.75 for PM-containing archaea and to 1.25 for other species (see YAML configuration files for details). In addition, three HMM profiles from NCBI CDD (Conserved Domain Database) (Lu et al. 2020) featuring ‘pseudomurein’ in their annotation were downloaded on December 18th, 2020. Then the profiles were used to identify homologues in the conceptual translations of the five PM-containing archaea with hmmsearch from the HMMER package v3.3 (Mistry et al. 2013) with default parameters. Matching protein sequences were graphically selected using the Ompa-Pa v0.211430 interactive software package (A. Bertrand and D. Baurain; https://metacpan.org/dist/Bio-MUST-Apps-OmpaPa) with the ‘max_copy’ option set to 20 and ‘min_cov’ to 0.7. Finally, the corresponding OGs were added to the selection.

### Genetic environment analysis of candidate proteins and *in-silico* characterization of their domains

Genetic environment databases were built for the genes of the selected OGs using the “3 in 1” module of GeneSpy (Garcia et al. 2019). Functional domains were predicted using InterProScan v5.37-76.0 (Jones et al. 2014), along with SignalP v5.0b (Almagro Armenteros et al. 2019) and TMHMM v2.0c (Krogh et al. 2001). InterProScan was used with default parameters and we disabled the precalculated match lookup, while the SignalP organism option was set to ‘arch’. To avoid misprediction by TMHMM, the signal peptide was first removed from the original sequences when the cleavage site prediction probability was greater than or equal to 0.1.

### Filtering of candidate proteins

To rescue potential pseudogenes or mistranslated proteins missing in selected OGs with protein sequences from only four (out of five) PM-containing archaea, Forty-Two v0.213470 was run in TBLASTN mode on the whole genomic sequences of the five PM-containing archaea. Then, classify-ali.pl was used again to retain only the OGs having sequences from all five PM-containing archaea. To enrich OGs with further archaeal orthologues, a second round of forty-two.pl in BLASTP mode was performed using the archaeal database of 819 whole genomes (see YAML configuration files for details). Each enriched OG was aligned using MAFFT L-INS-i v7.273 (Katoh and Standley 2013). From those alignments, HMM profiles were built using the HMMER package and bacterial homologues were identified separately in the bacterial and the prokaryotic databases. Protein sequences were graphically selected using Ompa-Pa with ‘max_copy’ and ‘min_cov’ options set to 20 and 0.7, respectively. For each OG, identical length and e-value thresholds were used for both databases when selecting homologous proteins.

### Phylogenetic analyses

#### ATP-grasp superfamily

In order to select a set of representative sequences containing the ATP-grasp domain, we first built a HMM profile from the alignment of the OG containing archaeal ATP-grasp domain proteins using the HMMER package. This profile was uploaded to the HMMER website (https://www.ebi.ac.uk/Tools/hmmer/search/hmmsearch) from which we retrieved homologous sequences (excluding eukaryotes) from the Swiss-Prot database (Poux et al. 2017). From those sequences, homologues were identified in our local bacterial databases using the HMM profile and Ompa-Pa. In parallel, the archaeal OGs (see Identification of candidate proteins for pseudomurein biosynthesis) homologous to the Swiss-Prot proteins were identified using NCBI BLASTp v2.2.28+ (Camacho et al. 2009) and enriched using Forty-Two with the archaeal database as ‘bank’. Finally, all archaeal and bacterial homologous sequences were merged into one single file.

To identify most ATP-grasp-containing domain proteins in our local databases, the merged file was aligned using MAFFT L-INS-i and the alignment was masked using the mask-ali.pl perl script (D. Baurain; https://metacpan.org/dist/Bio-MUST-Core) to isolate the ATP-grasp domain. From this domain alignment, an HMM profile was built using the HMMER package to identify ATP-grasp domain-containing homologues in our archaeal and bacterial databases, and homologous sequences were selected using Ompa-Pa. Protein sequences with two ATP-grasp domains (i.e., CarB) were cut at half-length, then both complete and half-sequences were aligned using MAFFT and their ATP-grasp domain again isolated using mask-ali.pl. Protein sequences were deduplicated using cdhit-tax-filter.pl perl script (V. Lupo and D. Baurain; https://metacpan.org/dist/Bio-MUST-Drivers) with the ‘keep-all’ option enabled and the identity threshold set to 0.65, then tagged using a BLAST-based annotation script (part of Bio-MUST-Drivers) and highly divergent sequences were removed using prune-outliers.pl v0.213470 with the ‘evalue’ option set to 1e-3, ‘min-hits’ to 1, ‘min_ident’ to 0.01 and ‘max_ident’ to 0.2. Finally, sequences were realigned with MAFFT L-INS-i. Conserved sites were selected using ali2phylip.pl v0.212670 (D. Baurain; https://metacpan.org/dist/Bio-MUST-Core) with the ‘min’ and ‘max’ options set to 0.3. The resulting matrix of 2,194 sequences x 180 AAs was used to infer phylogenetic domain trees using IQ-TREE v1.6.12 (Nguyen et al. 2015) with 1000 ultrafast bootstrap (UFBoot) replicates (Hoang et al. 2018) and under four models: LG4X+R4, C20+G4, C40+G4 and PMSF LG+C60+G4. In total, seven trees were computed because we tested the effect of increasing the number of iterations from 1000 to 3000 for the C20 and C40 models, and from 3000 to 5000 for the PMSF model.

#### MraY-like family

The two OGs (see Identification of candidate proteins for pseudomurein biosynthesis) containing proteins predicted with a domain glycosyltransferase 4 were enriched in bacterial homologues using Forty-Two in BLASTP mode. In parallel, representative sequences from other members of the MraY-like family (https://www.ncbi.nlm.nih.gov/Structure/cdd/cddsrv.cgi?uid=264002) were downloaded from the UniProtKB (The UniProt Consortium 2021) database: WecA (P0AC78, P0AC80, Q8Z38), GPT (P96000, B5IDH8) and WbpL (G3XD50, A0A379IBB8). The three files were then enriched in bacterial and archaeal (if any) homologues using Forty-Two. Finally, the five files were aligned using MAFFT L-INS-i.

To better explore the diversity of the MraY-like family, HMM profiles were built from those alignments and homologous sequences were selected from HMMER hits on the bacterial database using Ompa-Pa. All homologous protein sequences were merged into one file and tagged using a BLAST-based annotation script (part of Bio-MUST-Drivers) and aligned using MAFFT L-INS-i. Conserved sites were selected using ali2phylip.pl with the ‘min’ and ‘max’ options set to 0.2. A first guide tree was computed from the resulting matrix of 1070 sequences x 410 AAs using IQ-TREE with 1000 UFBoot under the LG4X+R4 model. From this guide tree and automated annotation, all sequences were manually tagged using ‘treeplot’ from the MUST software package (Philippe 1993). According to their annotation, protein sequences of each member of the MraY-like family were aligned using MAFFT L-INS-i, then all members were realigned using Two-Scalp v0.211710 (A. Bertrand, V. Lupo and D. Baurain; https://metacpan.org/dist/Bio-MUST-Apps-TwoScalp) with the ‘linsi’ option enabled. Finally, ali2phylip.pl was used to select conserved sites with the ‘min’ and ‘max’ options set to 0.2 and the resulting matrix of 1070 sequences x 408 AAs was used to infer phylogenetic trees with IQ-TREE under three models (i.e., LG4X+R4, C20+G4, C40+G4) and 1000 UFBoot.

#### Mur domain-containing family

After enrichment of the OGs with archaeal and bacterial homologues, the multiple OGs corresponding to the Mur domain-containing family were merged into one single (unaligned) file. In parallel, reference protein sequences from additional members of the Mur domain-containing family were downloaded into three separated files using the command-line version of the ‘efetch’ tool v10.4 from the NCBI Entrez Programming Utilities (E-utilities): CapB (P96736), MurT (Q8DNZ9, A0A0H3JUU7, A0A0H2WZQ7) and CphA (P56947, O86109, P58572). Forty-Two in BLASTP mode was run, in two rounds, on the four files, using both bacterial and archaeal databases as ‘bank’, in a final effort to sample the diversity of Mur domain-containing proteins. Then, fusion proteins were cut between the two protein domains and half-sequences with no Mur ligase domain were discarded. The enriched files were merged and protein sequences were deduplicated using the cdhit-tax-filter.pl with the ‘keep-all’ option enabled and the identity threshold set to 1. Mur domain-containing family proteins were tagged using a BLAST-based annotation script (part of Bio-MUST-Drivers) with an e-value threshold of 1e-20. Protein sequences were aligned using MAFFT (default mode) and conserved sites were selected using ali2phylip.pl with the ‘max’ option set to 0.3. A first guide tree was computed with IQ-TREE under the LG4X+R4 model with 1000 UFBoot. Based on the automatic annotation, all protein sequences were manually tagged following the guide tree using ‘treeplot’ from the MUST software package.

In order to improve phylogenetic analysis, the alignment of the Mur domain-containing family was refined as follows: 1) sequences from the different members of the family were exported to distinct files and aligned using MAFFT L-INS-i, 2) using the ‘ed’ programme from the MUST software package, misaligned sequences were manually transferred to a ‘.non’ file, and then, reduced files were realigned using MAFFT L-INS-i, 3) realigned files and ‘.non’ files were merged and all sequences were aligned using Two-Scalp with the ‘linsi’ and ‘keep-length’ options enabled. Conserved sites were selected using ali2phylip.pl with the ‘max’ and ‘min’ option set to 0.3. Phylogenetic analysis was performed on the resulting matrix of 3407 sequences x 550 AAs using IQ-TREE with 1000 UFBoot under three models of sequence evolution: LG4X+R4, C20+G4 and C40+G4.

From the alignment of the four bacterial muramyl ligases (MurCDEF), the four archaeal muramyl ligases (Murαßγδ) and the FGPS protein sequences, we have produced two more alignments: one where the N-ter and the C-ter domains of the protein sequences were trimmed, and another where we kept only the most conserved AAs. Fusion proteins were removed from those three alignments and protein sequences converted to a binary encoding to analyze indels (i.e., 0 for a gap or a missing character state and 1 for any AA). Short sequences were removed using ali2phylip.pl with the ‘min’ option set to 0.6. The three resulting matrices of 2997 sequences x 2243 AAs, 3001 sequences x 1799 AAs and 3004 sequences x 281 AAs, respectively, were used to infer phylogenetic trees with with RAxML v8.1.17 (Stamatakis 2014) under the BINGAMMAX model.

The jackknife.pl perl script (part of Bio-MUST-Drivers) was used for species resampling analysis with the ‘linsi’ option enabled, ‘min’ and ‘max’ set to 0.3 and ‘n-process’ to 1000. The one thousand resulting alignments were used to infer phylogenetic trees using IQ-TREE with 1000 UFBoot under the LG4X+R4, C20+G4 and C40+G4 models. Clan support values were assessed using the parse_consense_out.pl perl script (Baurain et al. 2010) with the ‘mode’ option set to ‘tree’. Consensus trees were computed from the 1000 replicate trees using ASTRAL v5.7.7 (Zhang et al. 2018) with default options.

## Acknowledgements

VL is supported by a FRIA (Fonds pour la Formation à la Recherche dans l’Industrie et dans l’Agriculture) fellowship of the F.R.S.-FNRS. FK is a research associate of the F.R.S.-FNRS. Computational resources were provided through two grants to DB (University of Liège “Crédit de démarrage 2012” SFRD-12/04; F.R.S.-FNRS “Crédit de recherche 2014” CDR J.0080.15), and by the Consortium des Équipements de Calcul Intensif (CÉCI) founded by the F.R.S.-FNRS. The authors thank Rosa Gago for her help with the graphical design of the figures.

## Authors’ Contributions

VL conceived the study and designed experiments, performed experiments, analyzed the data, drafted and drew the figures, wrote the manuscript and approved the final manuscript. DB conceived the study and designed experiments, analyzed the data, wrote and reviewed the manuscript and approved the final manuscript. FK conceived the study and designed experiments, performed experiments, analyzed the data, drew the figures, wrote and reviewed the manuscript and approved the final manuscript. CR, ER, LO and OJ performed experiments and approved the final manuscript.

## Notes

https://doi.org/10.6084/m9.figshare.21641612

### Competing Interest Statement

The authors have declared no competing interest.

