## Supplemental Figures for "Origin and Evolution of Pseudomurein Biosynthetic Gene Clusters"

(a)

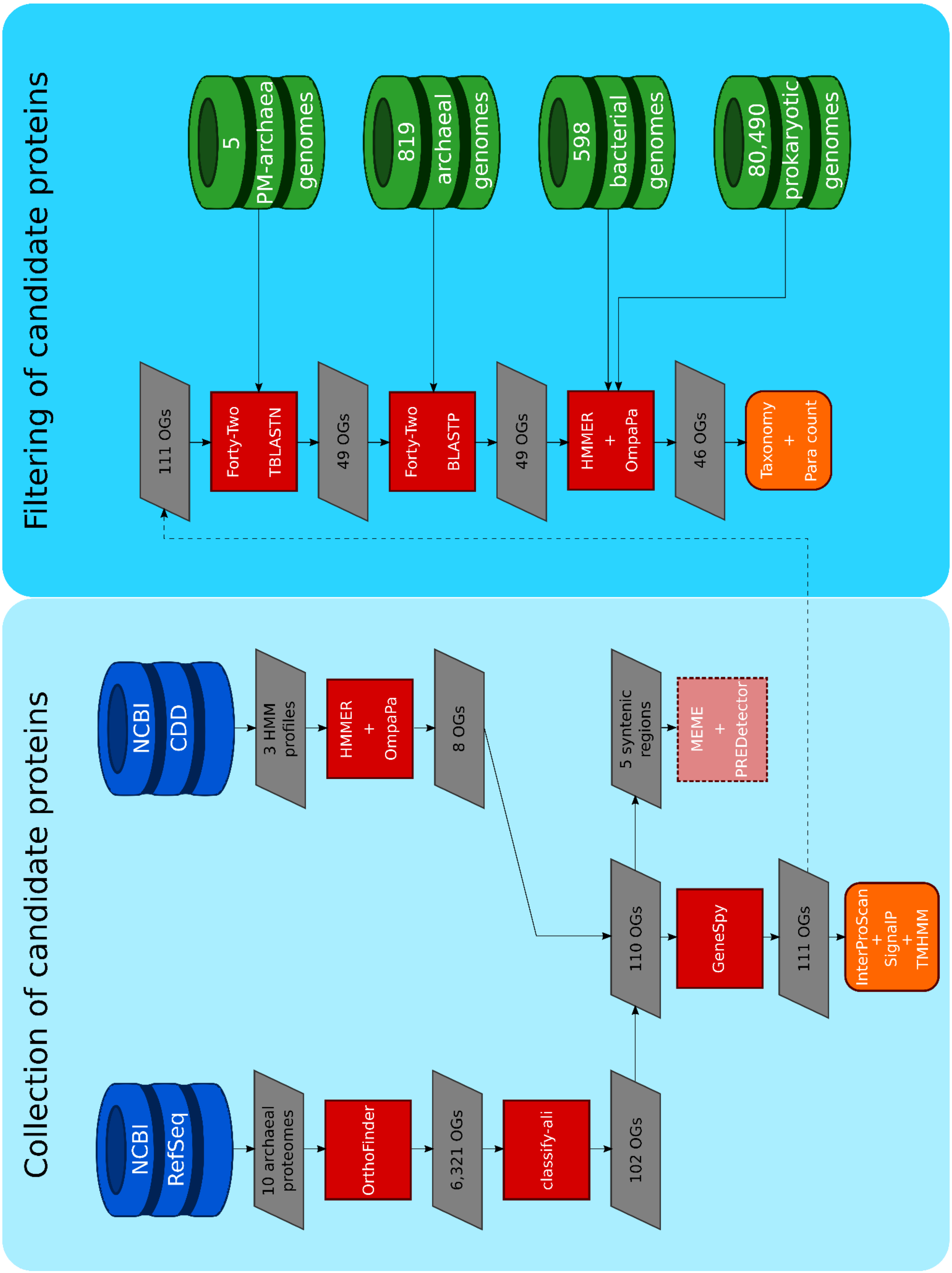


**Figure S1. Overview of the methods used for the bioinformatic analyses carried out during this study (see Material and Methods of the main text for**

**details). (a, previous page)** Main steps of the pipeline for the identification and filtering of the OGs potentially involved in PM biosynthesis. **(b, this page)** Details of the datasets and AA substitution models used for phylogenetic analyses.

**(b)**

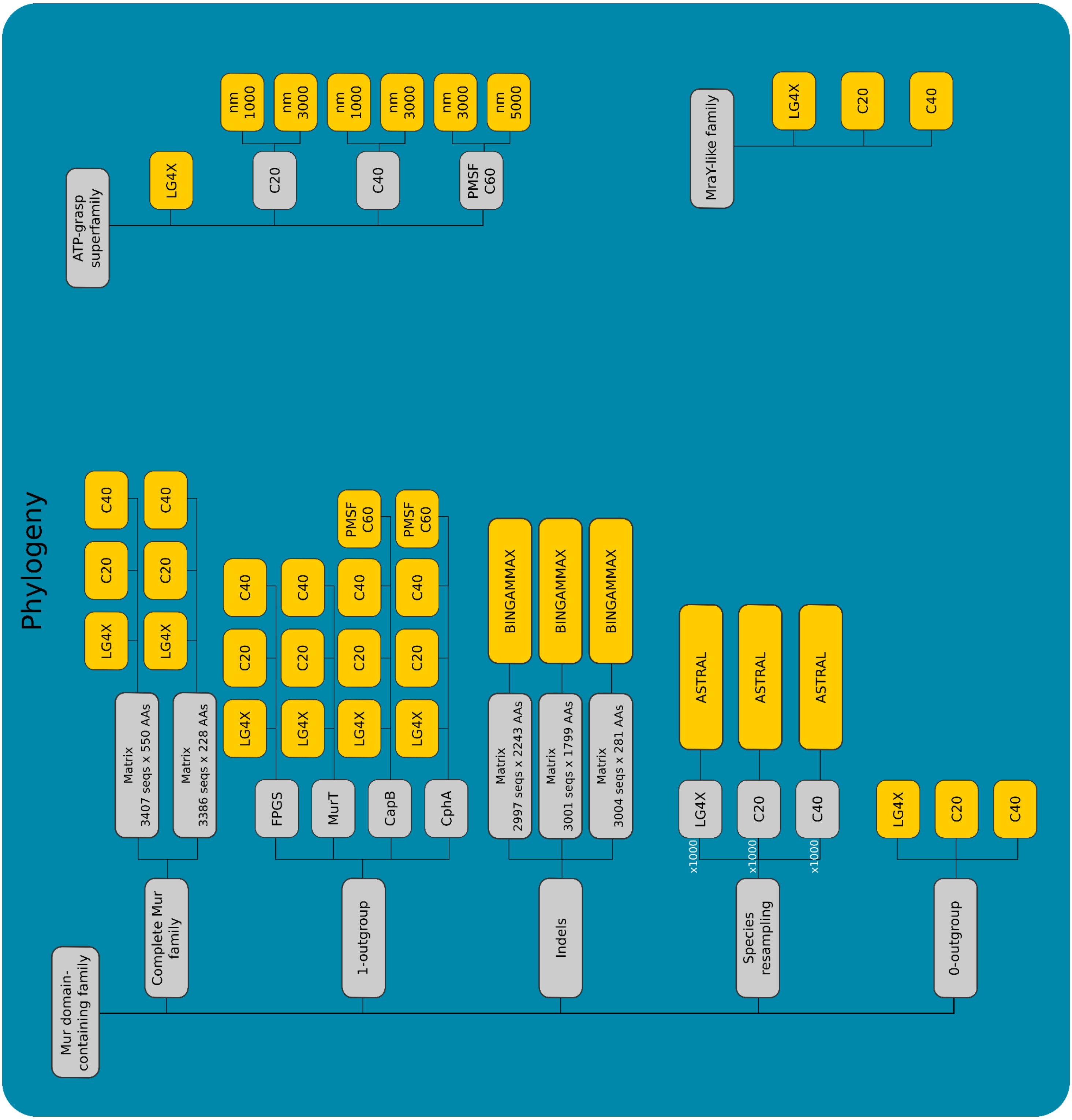

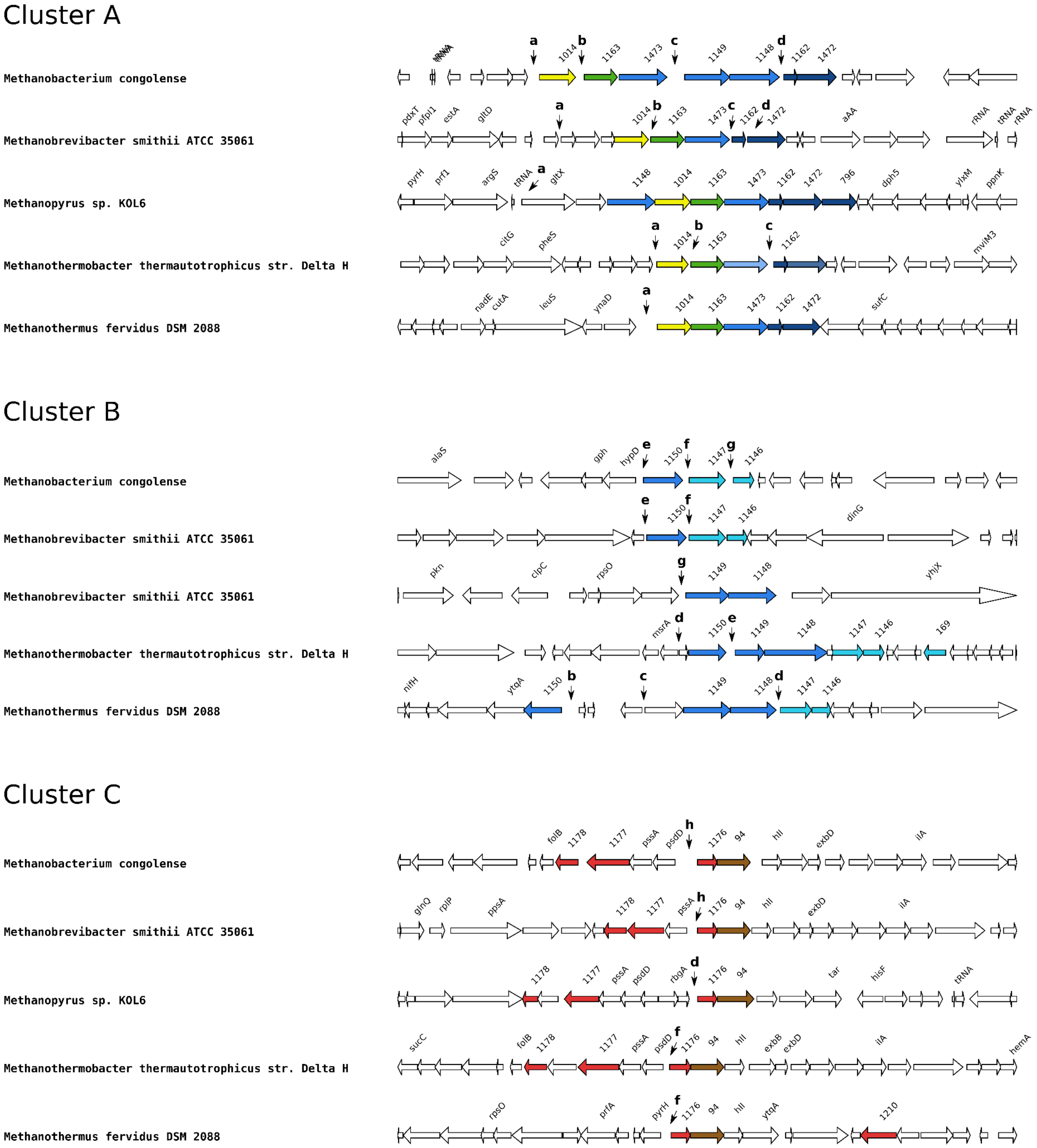

Cluster A
Methanobacterium congolense
a b c d
1014 1163 1473 1149 1148 1162 1472
Methanobrevibacter smithii ATCC 35061
pdxT pfpI1 estA gltD
1014 1163 1473 1162 1472
aAA rRNA tRNA rRNA
Methanopyrus sp. KOL6
pyrH prf1 argS tRNA gltX
1148 1014 1163 1473 1162 1472 796 dph5 ylxM ppnK
Methanothermobacter thermautotrophicus str. Delta H
citG pheS
1014 1163 1162 mviM3
Methanothermus fervidus DSM 2088
nadE cutA leuS ynaD
1014 1163 1473 1162 1472 sufC
Cluster B
Methanobacterium congolense
alaS gph hypD
e f g
1150 1147 1146
Methanobrevibacter smithii ATCC 35061
1150 1147 1146 dinG
Methanobrevibacter smithii ATCC 35061
pkn clpC rpsO
1149 1148 yhjX
Methanothermobacter thermautotrophicus str. Delta H
msrA 1150 1149 1148 1147 1146 169
Methanothermus fervidus DSM 2088
nifH ytqA 1150
1149 1148 1147 1146
Cluster C
Methanobacterium congolense
folB 1178 1177 pssA psdD
h
1176 94 hll exbD ilA
Methanobrevibacter smithii ATCC 35061
glnQ rplP ppsA 1178 1177 pssA 1176 94 hll exbD ilA
Methanopyrus sp. KOL6
1178 1177 pssA psdD rbgA 1176 94 tar hisF tRNA
Methanothermobacter thermautotrophicus str. Delta H
sucC folB 1178 1177 pssA psdD 1176 94 hll exbB exbD ilA hemA
Methanothermus fervidus DSM 2088
rpsO prfA pyrH 1176 94 hll ytqA 1210

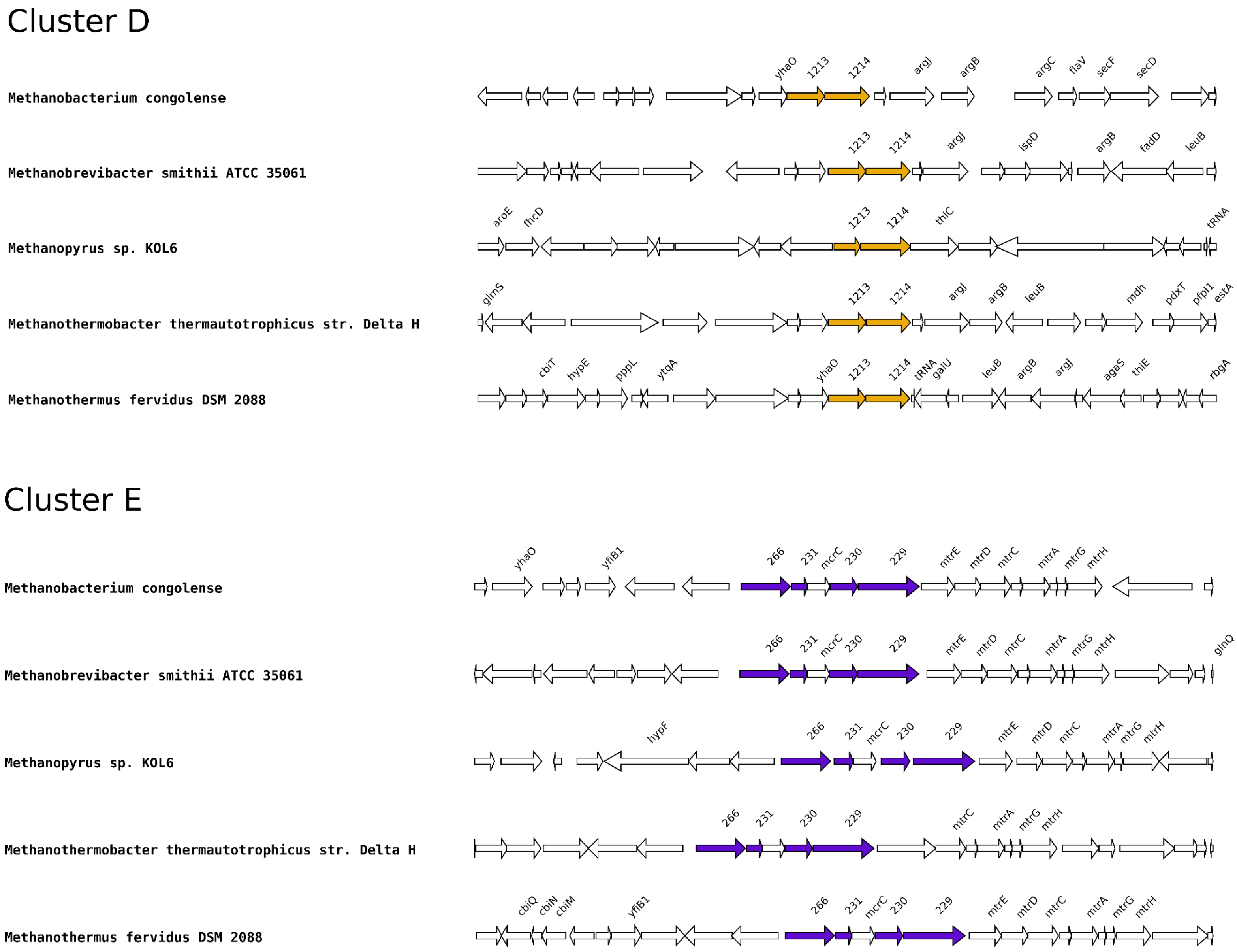


**Figure S2. Genetic organization of the five clusters.** Identified clustered genes are colored, whereas other genes are left in white. Above each identified gene is indicated its corresponding orthologous group (OG) number (without 'OG' and the following 0s). Intergenic regions used during the regulon pipeline (see Supplemental data) are indicated with a lowercase letter (Table S4). The intergenic regions "b" and "c" of *Methanopyrus* sp. are not shown in this figure and correspond to the upstream region of the OG0001150 and OG0001147, genes respectively.

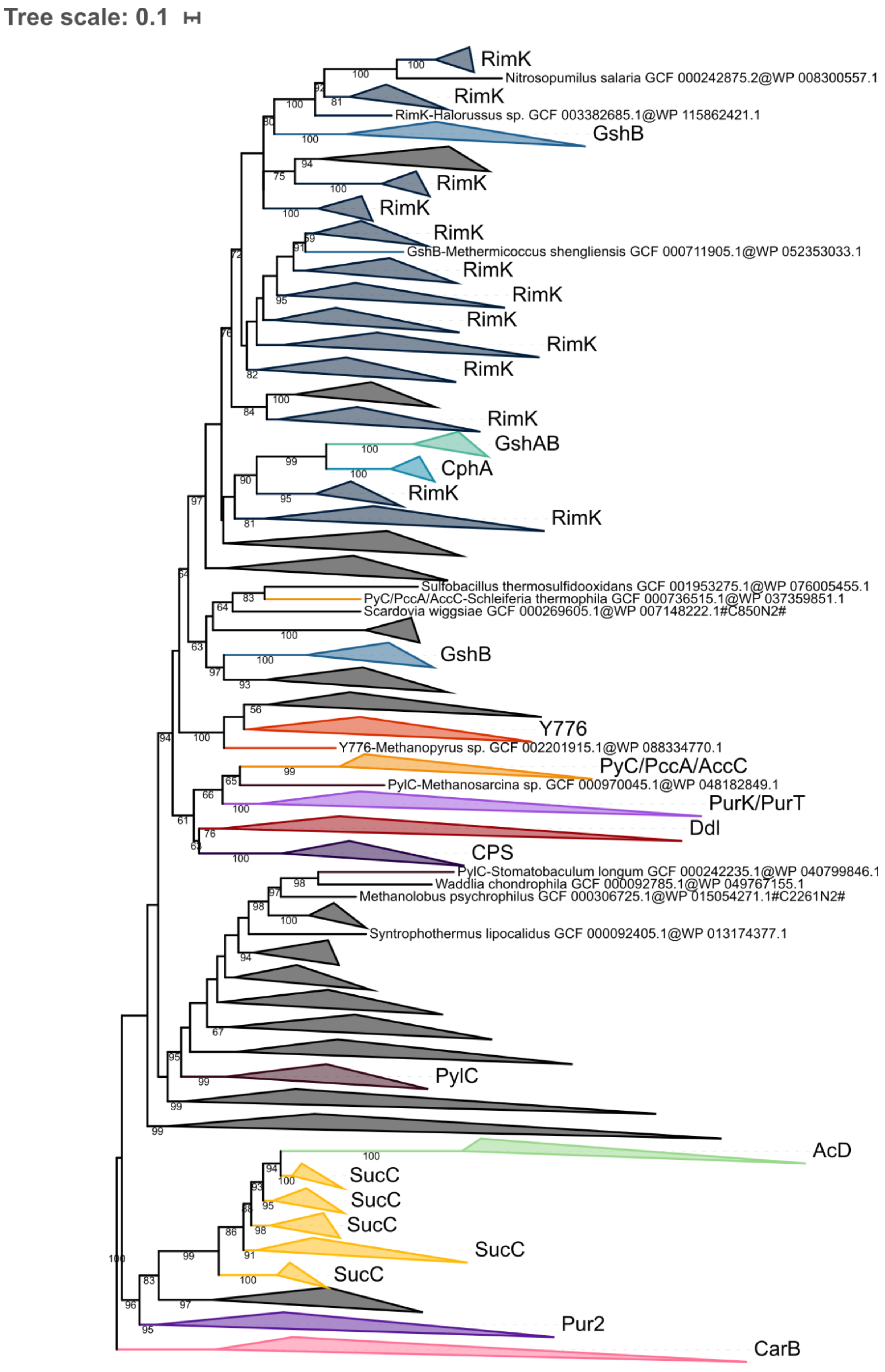
Tree scale: 0.1
RimK
Nitrosopumilus salaria GCF 000242875.2@WP 008300557.1
RimK
RimK-Halorussus sp. GCF 003382685.1@WP 115862421.1
GshB
RimK
RimK
RimK
GshB-Methermicoccus shengliensis GCF 000711905.1@WP 052353033.1
RimK
RimK
RimK
RimK
RimK
RimK
GshAB
CphA
RimK
RimK
Sulfobacillus thermosulfidooxidans GCF 001953275.1@WP 076005455.1
PyC/PccA/AccC-Schleiferia thermophila GCF 000736515.1@WP 037359851.1
Scardovia wiggsiae GCF 000269605.1@WP 007148222.1#C850N2#
GshB
Y776
Y776-Methanopyrus sp. GCF 002201915.1@WP 088334770.1
PyC/PccA/AccC
PylC-Methanosarcina sp. GCF 000970045.1@WP 048182849.1
PurK/PurT
Ddl
CPS
PylC-Stomatobaculum longum GCF 000242235.1@WP 040799846.1
Waddlia chondrophila GCF 000092785.1@WP 049767155.1
Methanolobus psychrophilus GCF 000306725.1@WP 015054271.1#C2261N2#
Syntrophothermus lipocalidus GCF 000092405.1@WP 013174377.1
PylC
AcD
SucC
SucC
SucC
SucC
SucC
Pur2
CarB

**Figure S3. Phylogenetic tree of the ATP-grasp superfamily rooted on CarB.** The tree was inferred from a matrix of 2,194 sequences x 180 unambiguously aligned AAs using IQ-TREE **under the C20+G4 model with 1000 iterations**. Tree visualization was performed using iTOL. Bootstrap support values are shown if greater or equal to 50. Branches were collapsed on homogeneous sequence annotation based on reference sequences. Black collapsed branches correspond to unannotated sequences.

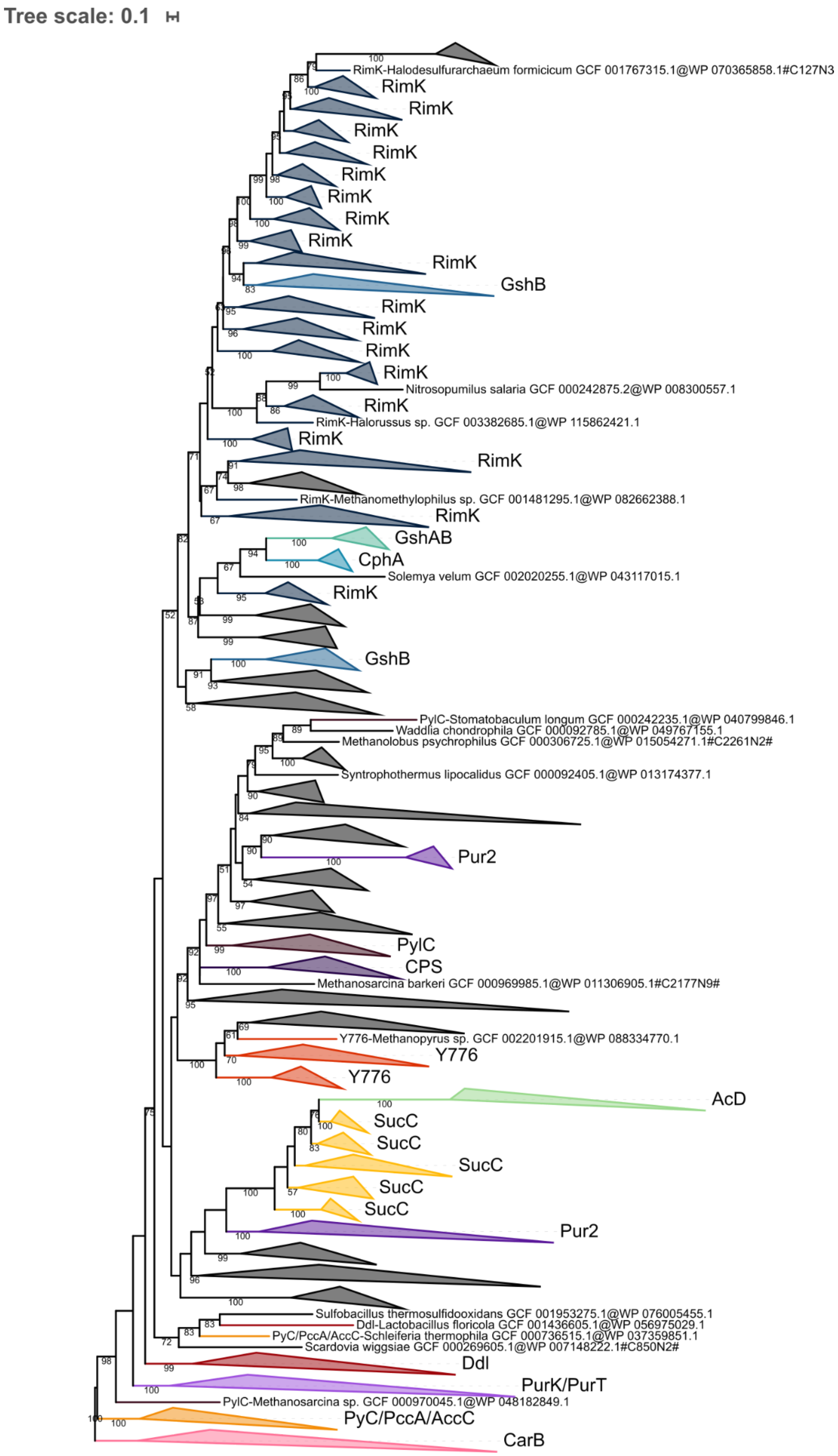
Tree scale: 0.1
RimK-Halodesulfurarchaeum formicicum GCF 001767315.1@WP 070365858.1#C127N3
RimK
RimK
RimK
RimK
RimK
RimK
RimK
RimK
RimK
GshB
RimK
RimK
RimK
RimK
Nitrosopumilus salaria GCF 000242875.2@WP 008300557.1
RimK
RimK-Halorussus sp. GCF 003382685.1@WP 115862421.1
RimK
RimK
RimK-Methanomethylophilus sp. GCF 001481295.1@WP 082662388.1
RimK
GshAB
CphA
Solemya velum GCF 002020255.1@WP 043117015.1
RimK
GshB
PylC-Stomatobaculum longum GCF 000242235.1@WP 040799846.1
Waddlia chondrophila GCF 000092785.1@WP 049767155.1
Methanolobus psychrophilus GCF 000306725.1@WP 015054271.1#C2261N2#
Syntrophothermus lipocalidus GCF 000092405.1@WP 013174377.1
Pur2
PylC
CPS
Methanosarcina barkeri GCF 000969985.1@WP 011306905.1#C2177N9#
Y776-Methanopyrus sp. GCF 002201915.1@WP 088334770.1
Y776
Y776
AcD
SucC
SucC
SucC
SucC
SucC
Pur2
Sulfobacillus thermosulfidooxidans GCF 001953275.1@WP 076005455.1
Ddl-Lactobacillus floricola GCF 001436605.1@WP 056975029.1
PyC/PccA/AccC-Schleiferia thermophila GCF 000736515.1@WP 037359851.1
Scardovia wiggsiae GCF 000269605.1@WP 007148222.1#C850N2#
Ddl
PurK/PurT
PylC-Methanosarcina sp. GCF 000970045.1@WP 048182849.1
PyC/PccA/AccC
CarB

**Figure S4. Phylogenetic tree of the ATP-grasp superfamily rooted on CarB.** The tree was inferred from a matrix of 2,194 sequences x 180 unambiguously aligned AAs using IQ-TREE **under the C20+G4 model with 3000 iterations**. Tree visualization was performed using iTOL. Bootstrap support values are shown if greater or equal to 50. Branches were collapsed on homogeneous sequence annotation based on reference sequences. Black collapsed branches correspond to unannotated sequences.

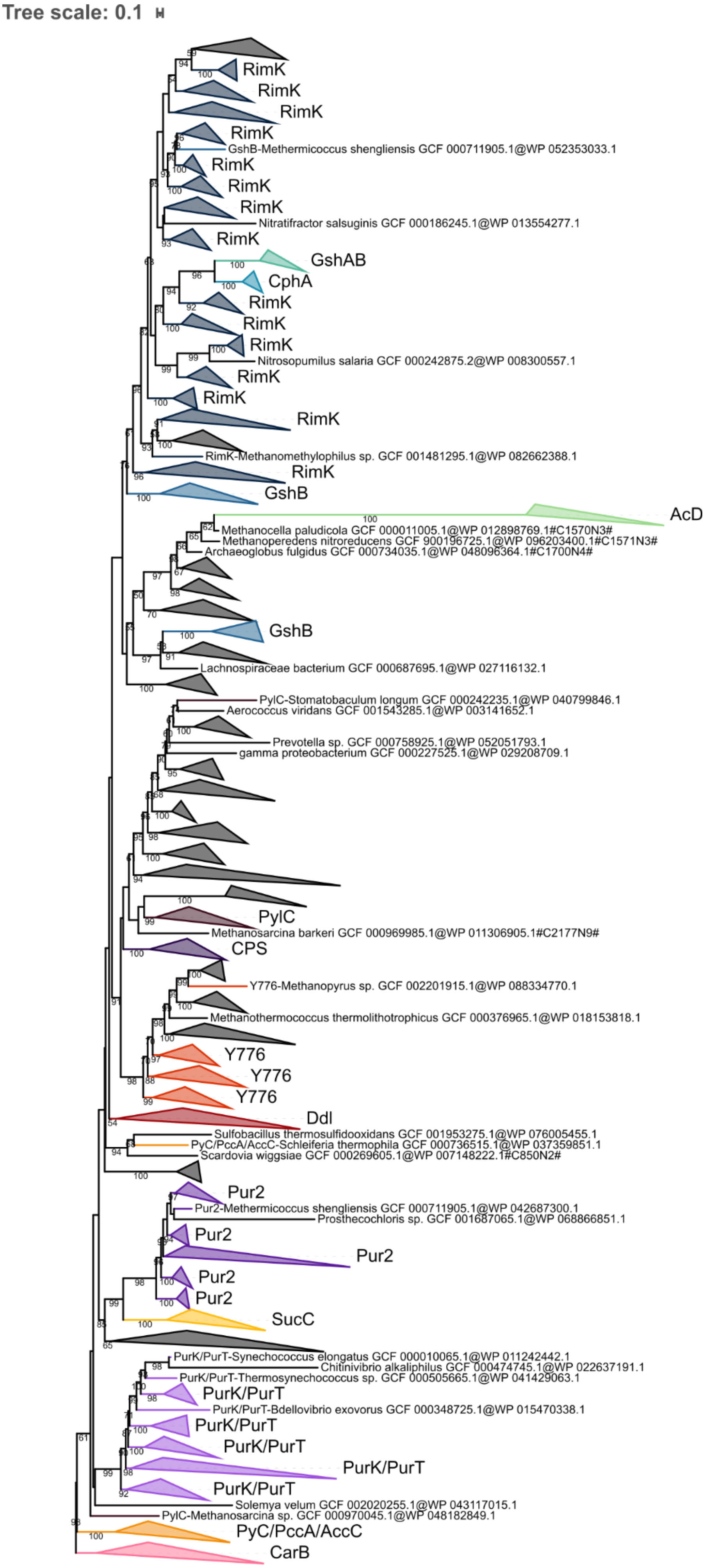
Tree scale: 0.1
RimK
RimK
RimK
RimK
GshB-Methermicoccus shengliensis GCF 000711905.1@WP 052353033.1
RimK
RimK
RimK
Nitratifractor salsuginis GCF 000186245.1@WP 013554277.1
RimK
GshAB
CphA
RimK
RimK
RimK
Nitrosopumilus salaria GCF 000242875.2@WP 008300557.1
RimK
RimK
RimK
RimK-Methanomethylophilus sp. GCF 001481295.1@WP 082662388.1
RimK
GshB
AcD
Methanocella paludicola GCF 000011005.1@WP 012898769.1#C1570N3#
Methanoperedens nitroreducens GCF 900196725.1@WP 096203400.1#C1571N3#
Archaeoglobus fulgidus GCF 000734035.1@WP 048096364.1#C1700N4#
GshB
Lachnospiraceae bacterium GCF 000687695.1@WP 027116132.1
PylC-Stomatobaculum longum GCF 000242235.1@WP 040799846.1
Aerococcus viridans GCF 001543285.1@WP 003141652.1
Prevotella sp. GCF 000758925.1@WP 052051793.1
gamma proteobacterium GCF 000227525.1@WP 029208709.1
PylC
Methanosarcina barkeri GCF 000969985.1@WP 011306905.1#C2177N9#
CPS
Y776-Methanopyrus sp. GCF 002201915.1@WP 088334770.1
Methanothermococcus thermolithotrophicus GCF 000376965.1@WP 018153818.1
Y776
Y776
Y776
Ddl
Sulfobacillus thermosulfidooxidans GCF 001953275.1@WP 076005455.1
PyC/PccA/AccC-Schleiferia thermophila GCF 000736515.1@WP 037359851.1
Scardovia wiggsiae GCF 000269605.1@WP 007148222.1#C850N2#
Pur2
Pur2-Methermicoccus shengliensis GCF 000711905.1@WP 042687300.1
Prosthecochloris sp. GCF 001687065.1@WP 068866851.1
Pur2
Pur2
Pur2
Pur2
SucC
PurK/PurT-Synechococcus elongatus GCF 000010065.1@WP 011242442.1
Chitinivibrio alkaliphilus GCF 000474745.1@WP 022637191.1
PurK/PurT-Thermosynechococcus sp. GCF 000505665.1@WP 041429063.1
PurK/PurT
PurK/PurT-Bdellovibrio exovorus GCF 000348725.1@WP 015470338.1
PurK/PurT
PurK/PurT
PurK/PurT
PurK/PurT
Solemya velum GCF 002020255.1@WP 043117015.1
PylC-Methanosarcina sp. GCF 000970045.1@WP 048182849.1
PyC/PccA/AccC
CarB

**Figure S5. Phylogenetic tree of the ATP-grasp superfamily rooted on CarB.** The tree was inferred from a matrix of 2,194 sequences x 180 unambiguously aligned AAs using IQ-TREE **under the C40+G4 model with 1000 iterations**. Tree visualization was performed using iTOL. Bootstrap support values are shown if greater or equal to 50. Branches were collapsed on homogeneous sequence annotation based on reference sequences. Black collapsed branches correspond to unannotated sequences.

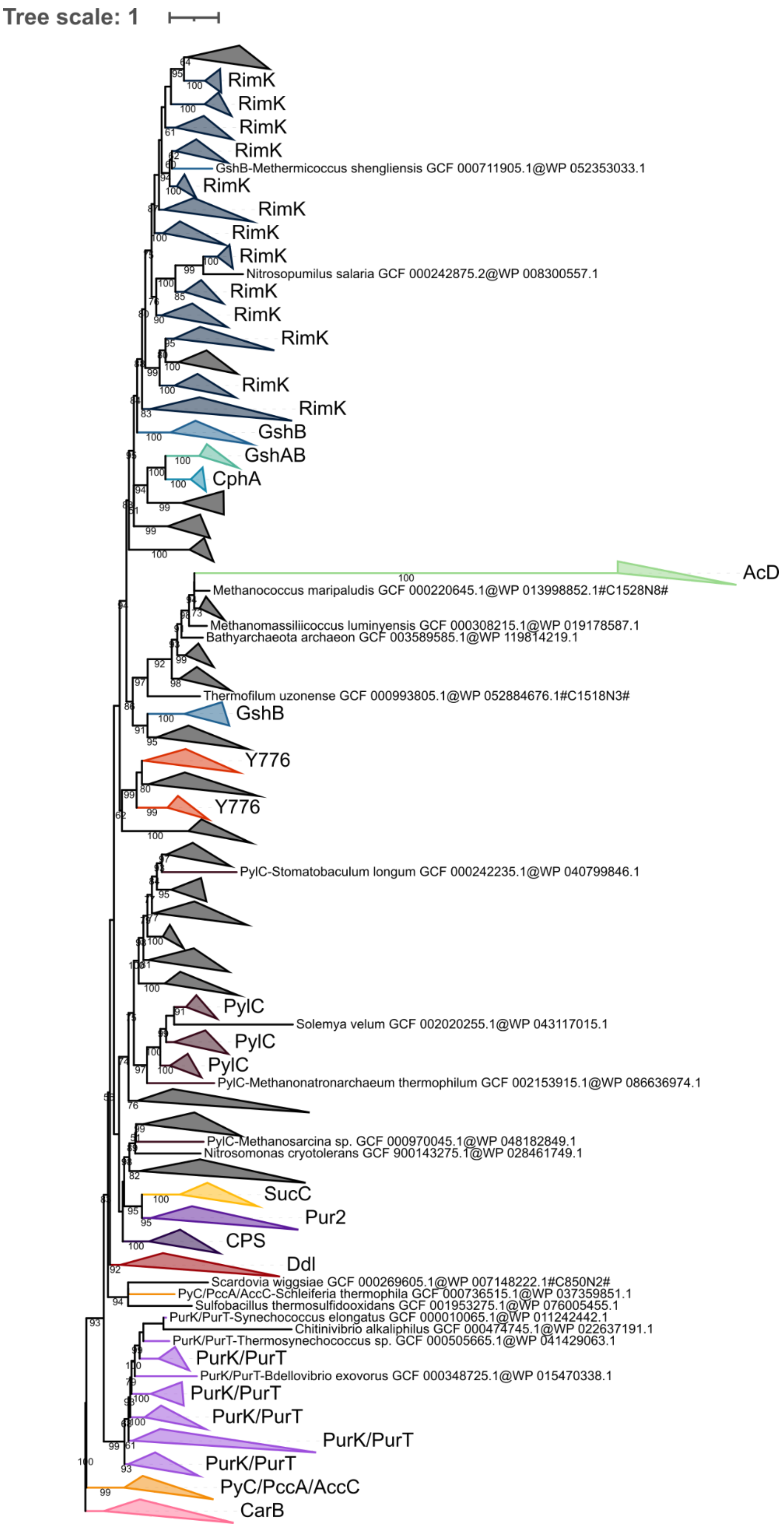

Tree scale: 1
RimK
RimK
RimK
RimK
GshB-Methermicoccus shengliensis GCF 000711905.1@WP 052353033.1
RimK
RimK
RimK
RimK
Nitrosopumilus salaria GCF 000242875.2@WP 008300557.1
RimK
RimK
RimK
RimK
RimK
GshB
GshAB
CphA
AcD
Methanococcus maripaludis GCF 000220645.1@WP 013998852.1#C1528N8#
Methanomassiliicoccus luminyensis GCF 000308215.1@WP 019178587.1
Bathyarchaeota archaeon GCF 003589585.1@WP 119814219.1
Thermofilum uzonense GCF 000993805.1@WP 052884676.1#C1518N3#
GshB
Y776
Y776
PylC-Stomatobaculum longum GCF 000242235.1@WP 040799846.1
PylC
Solemya velum GCF 002020255.1@WP 043117015.1
PylC
PylC
PylC-Methanonatronarchaeum thermophilum GCF 002153915.1@WP 086636974.1
PylC-Methanosarcina sp. GCF 000970045.1@WP 048182849.1
Nitrosomonas cryotolerans GCF 900143275.1@WP 028461749.1
SucC
Pur2
CPS
Ddl
Scardovia wiggsiae GCF 000269605.1@WP 007148222.1#C850N2#
PyC/PccA/AccC-Schleiferia thermophila GCF 000736515.1@WP 037359851.1
Sulfobacillus thermosulfidooxidans GCF 001953275.1@WP 076005455.1
PurK/PurT-Synechococcus elongatus GCF 000010065.1@WP 011242442.1
Chitinivibrio alkaliphilus GCF 000474745.1@WP 022637191.1
PurK/PurT-Thermosynechococcus sp. GCF 000505665.1@WP 041429063.1
PurK/PurT
PurK/PurT-Bdellovibrio exovorus GCF 000348725.1@WP 015470338.1
PurK/PurT
PurK/PurT
PurK/PurT
PurK/PurT
PyC/PccA/AccC
CarB

**Figure S6. Phylogenetic tree of the ATP-grasp superfamily rooted on CarB.** The tree was inferred from a matrix of 2,194 sequences x 180 unambiguously aligned AAs using IQ-TREE **under the C40+G4 model with 3000 iterations**. Tree visualization was performed using iTOL. Bootstrap support values are shown if greater or equal to 50. Branches were collapsed on homogeneous sequence annotation based on reference sequences. Black collapsed branches correspond to unannotated sequences.

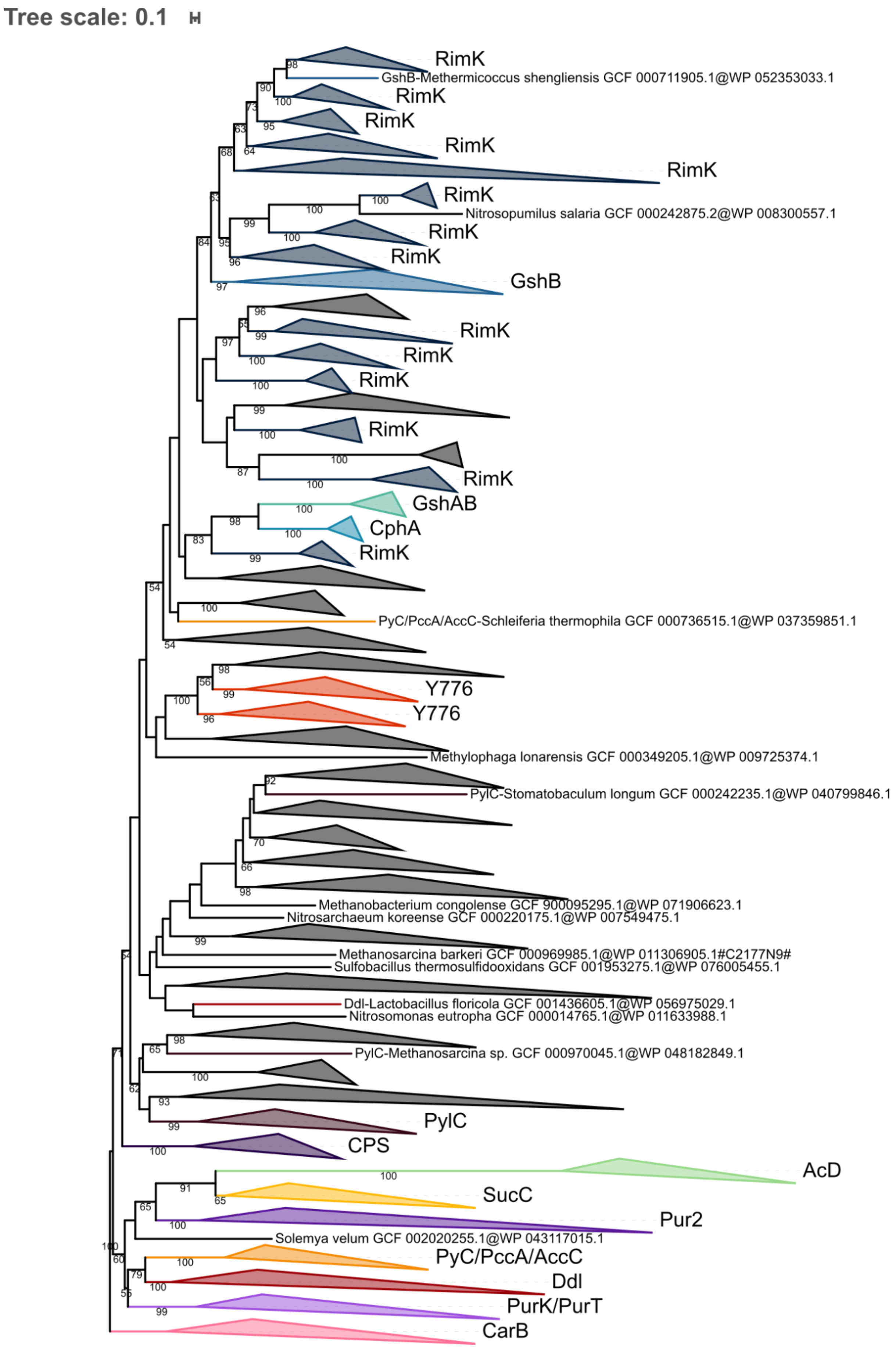
Tree scale: 0.1
RimK
GshB-Methermicoccus shengliensis GCF 000711905.1@WP 052353033.1
RimK
RimK
RimK
RimK
RimK
Nitrosopumilus salaria GCF 000242875.2@WP 008300557.1
RimK
RimK
GshB
RimK
RimK
RimK
RimK
RimK
GshAB
CphA
RimK
PyC/PccA/AccC-Schleiferia thermophila GCF 000736515.1@WP 037359851.1
Y776
Y776
Methylophaga lonarensis GCF 000349205.1@WP 009725374.1
PylC-Stomatobaculum longum GCF 000242235.1@WP 040799846.1
Methanobacterium congolense GCF 900095295.1@WP 071906623.1
Nitrosarchaeum koreense GCF 000220175.1@WP 007549475.1
Methanosarcina barkeri GCF 000969985.1@WP 011306905.1#C2177N9#
Sulfobacillus thermosulfidooxidans GCF 001953275.1@WP 076005455.1
Ddl-Lactobacillus floricola GCF 001436605.1@WP 056975029.1
Nitrosomonas eutropha GCF 000014765.1@WP 011633988.1
PylC-Methanosarcina sp. GCF 000970045.1@WP 048182849.1
PylC
CPS
AcD
SucC
Pur2
Solemya velum GCF 002020255.1@WP 043117015.1
PyC/PccA/AccC
Ddl
PurK/PurT
CarB

**Figure S7. Phylogenetic tree of the ATP-grasp superfamily rooted on CarB.** The tree was inferred from a matrix of 2,194 sequences x 180 unambiguously aligned AAs using IQ-TREE **under the PMSF LG+C60+G4 model with 3000 iterations**. Tree visualization was performed using iTOL. Bootstrap support values are shown if greater or equal to 50. Branches were collapsed on homogeneous sequence annotation based on reference sequences. Black collapsed branches correspond to unannotated sequences.

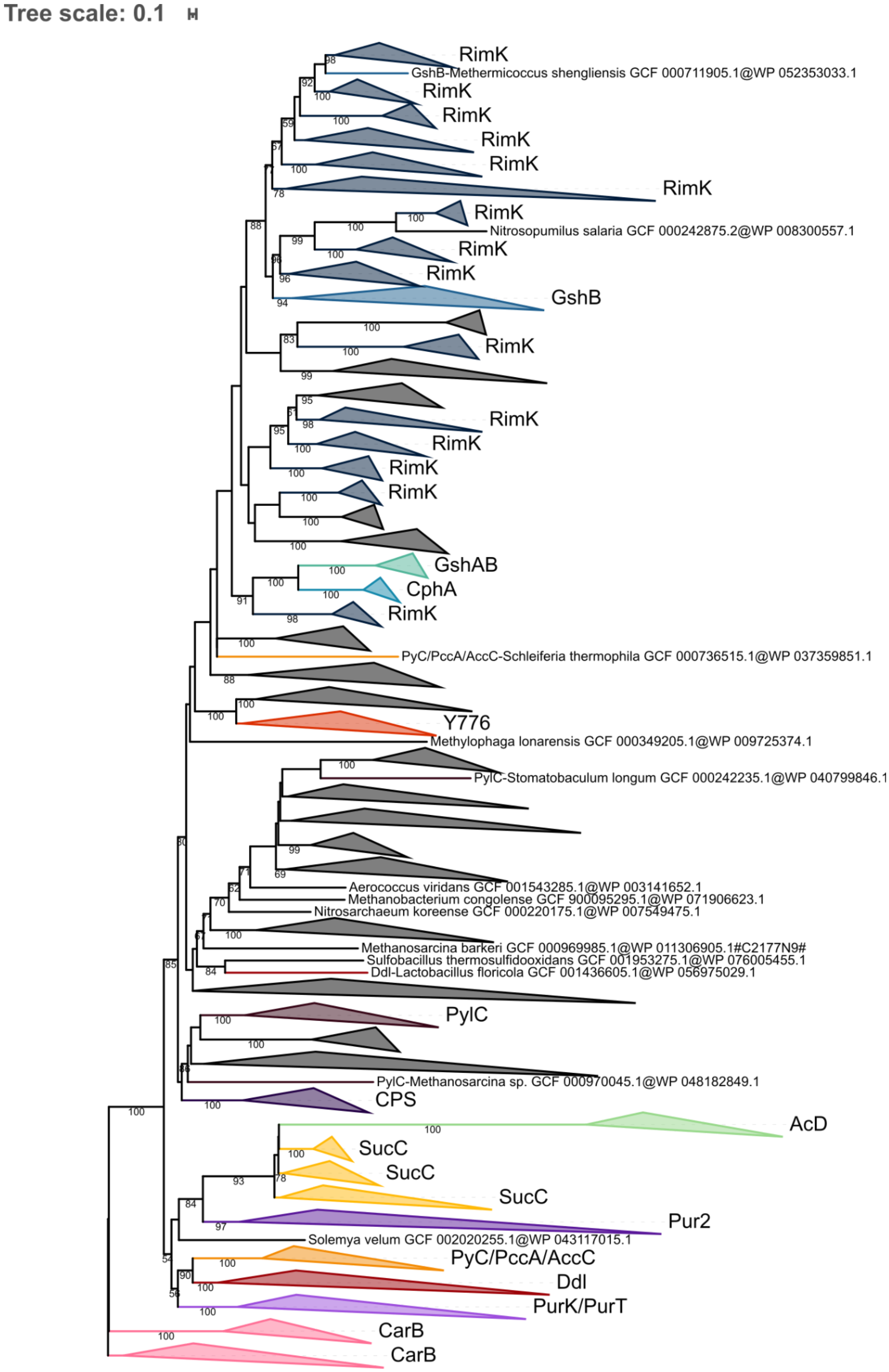
Tree scale: 0.1
RimK
GshB-Methermicoccus shengliensis GCF 000711905.1@WP 052353033.1
RimK
RimK
RimK
RimK
RimK
RimK
Nitrosopumilus salaria GCF 000242875.2@WP 008300557.1
RimK
RimK
GshB
RimK
RimK
RimK
RimK
RimK
GshAB
CphA
RimK
PyC/PccA/AccC-Schleiferia thermophila GCF 000736515.1@WP 037359851.1
Y776
Methylophaga lonarensis GCF 000349205.1@WP 009725374.1
PylC-Stomatobaculum longum GCF 000242235.1@WP 040799846.1
Aerococcus viridans GCF 001543285.1@WP 003141652.1
Methanobacterium congolense GCF 900095295.1@WP 071906623.1
Nitrosarchaeum koreense GCF 000220175.1@WP 007549475.1
Methanosarcina barkeri GCF 000969985.1@WP 011306905.1#C2177N9#
Sulfobacillus thermosulfidooxidans GCF 001953275.1@WP 076005455.1
Ddl-Lactobacillus floricola GCF 001436605.1@WP 056975029.1
PylC
PylC-Methanosarcina sp. GCF 000970045.1@WP 048182849.1
CPS
AcD
SucC
SucC
SucC
Pur2
Solemya velum GCF 002020255.1@WP 043117015.1
PyC/PccA/AccC
Ddl
PurK/PurT
CarB
CarB

**Figure S8. Phylogenetic tree of the ATP-grasp superfamily rooted on CarB.** The tree was inferred from a matrix of 2,194 sequences x 180 unambiguously aligned AAs using IQ-TREE **under the PMSF LG+C60+G4 model with 5000 iterations**. Tree visualization was performed using iTOL. Bootstrap support values are shown if greater or equal to 50. Branches were collapsed on homogeneous sequence annotation based on reference sequences. Black collapsed branches correspond to unannotated sequences.

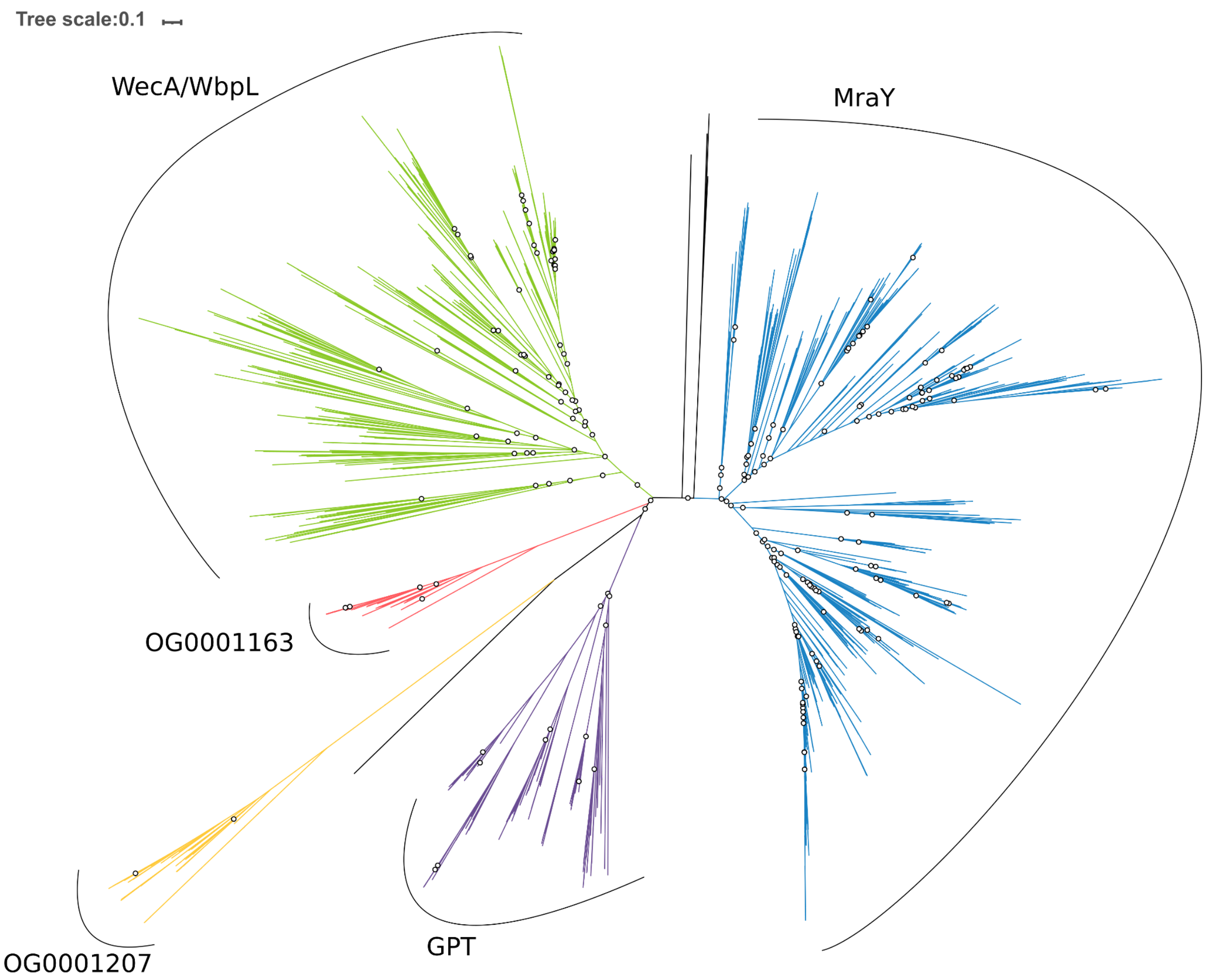


**Figure S9. Unrooted phylogenetic tree of the MraY-like family.** The tree was constructed from a matrix of 1,070 sequences x 408 unambiguously aligned AAs using IQ-TREE **under the LG4X+R4 model**. Open circles correspond to bootstrap support values lower than 90. Blue sequences correspond to a MraY annotation, green to WecA/WbpL, red to OG0001163 (MraY-like), yellow to OG0001207, purple to GPT, and black to unannotated bacterial sequences.

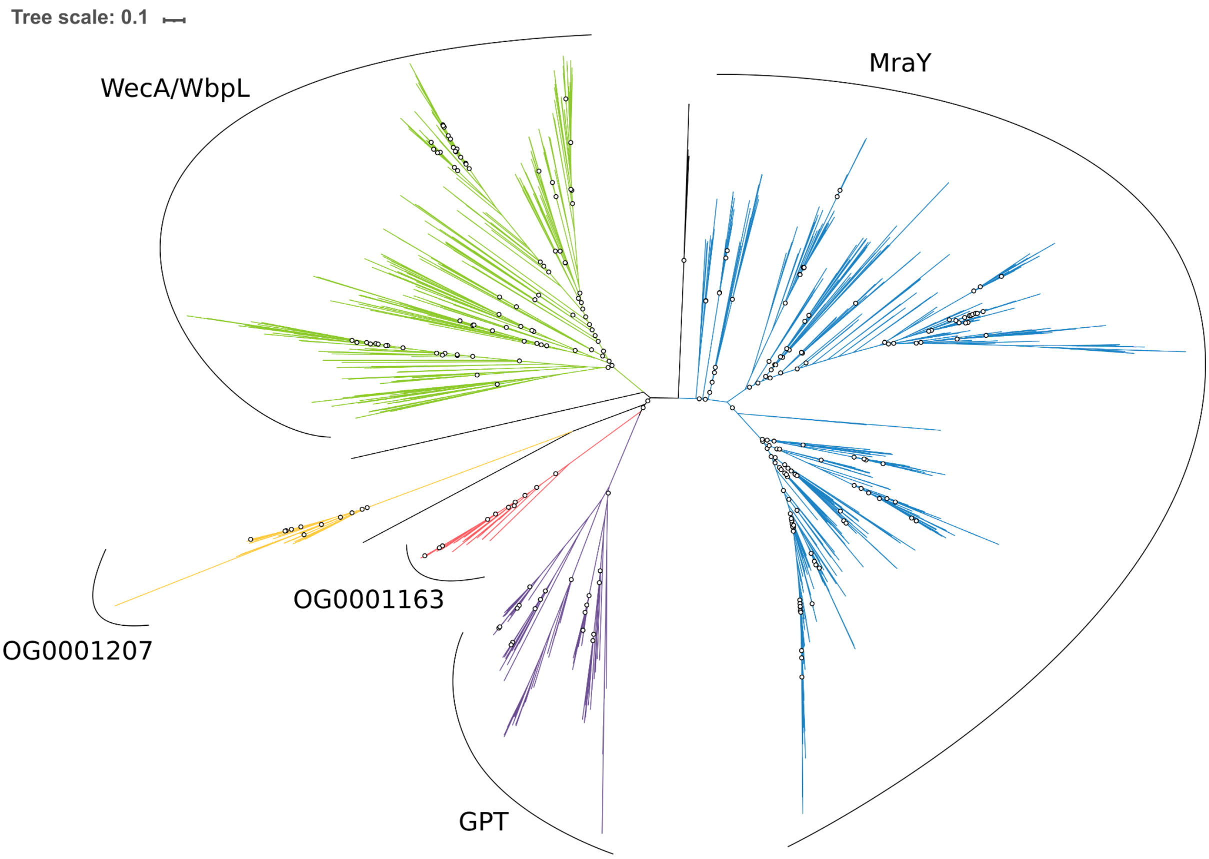


**Figure S10. Unrooted phylogenetic tree of the MraY-like family.** The tree was constructed from a matrix of 1,070 sequences x 408 unambiguously aligned AAs using IQ-TREE **under the C20+G4 model**. Open circles correspond to bootstrap support values lower than 90. Blue sequences correspond to a MraY annotation, green to WecA/WbpL, red to OG0001163 (MraY-like), yellow to OG0001207, purple to GPT, and black to unannotated bacterial sequences.

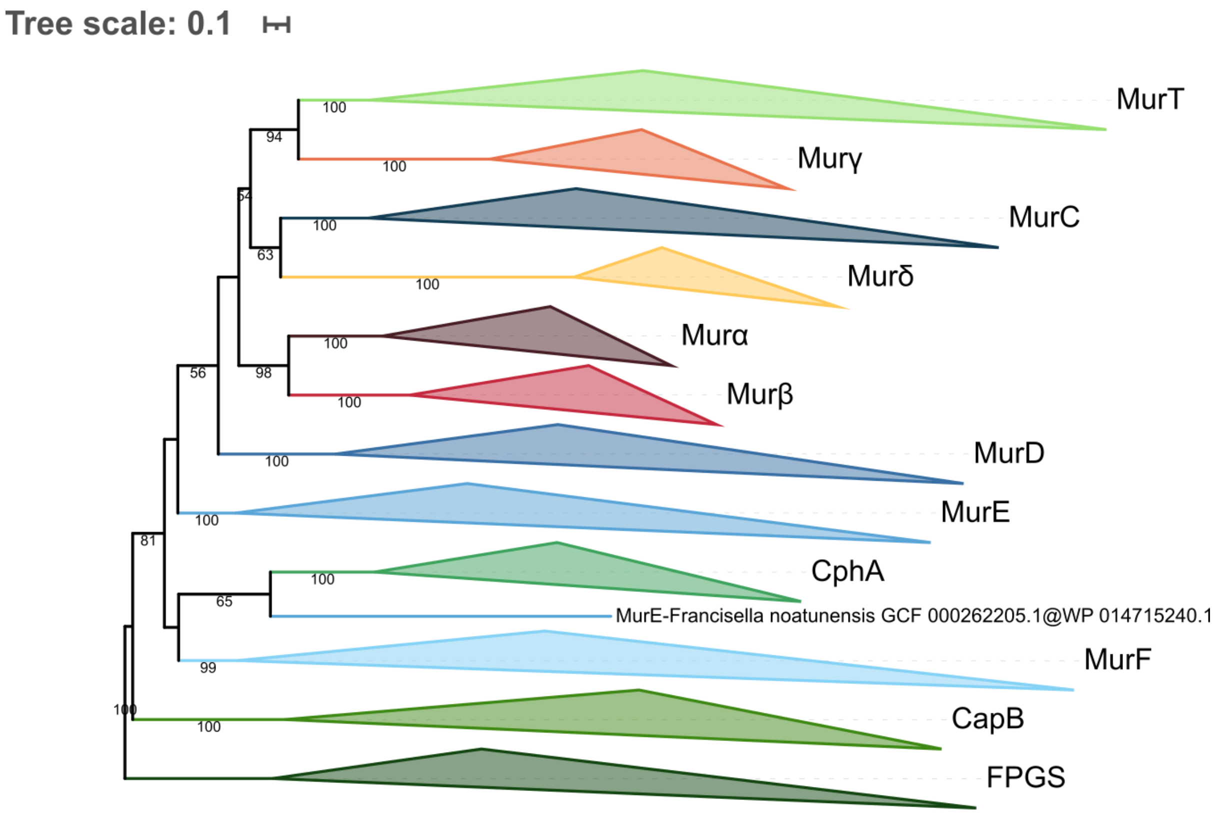


**Figure S11. Phylogenetic tree of the Mur domain-containing family rooted on FPGS.** The tree was inferred from a matrix of **3407 sequences x 550 unambiguously aligned AAs** using IQ-TREE **under the LG4X+R4 model**. Tree visualization was performed using iTOL. Bootstrap values are shown if greater or equal to 50. Branches were collapsed on homogeneous sequence annotation.

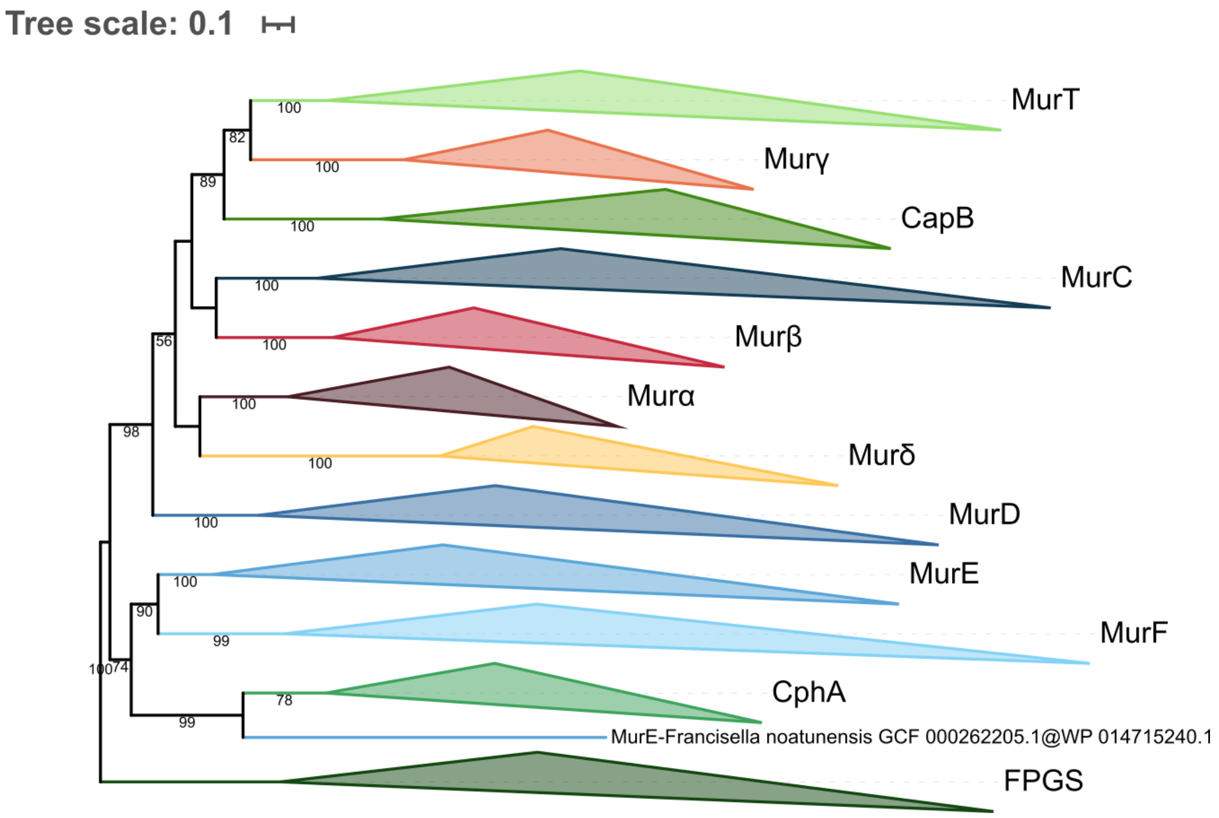


**Figure S12. Phylogenetic tree of the Mur domain-containing family rooted on FPGS.** The tree was inferred from a matrix of **3407 sequences x 550 unambiguously aligned AAs** using IQ-TREE **under the C20+G4 model**. Tree visualization was performed using iTOL. Bootstrap values are shown if greater or equal to 50. Branches were collapsed on homogeneous sequence annotation.

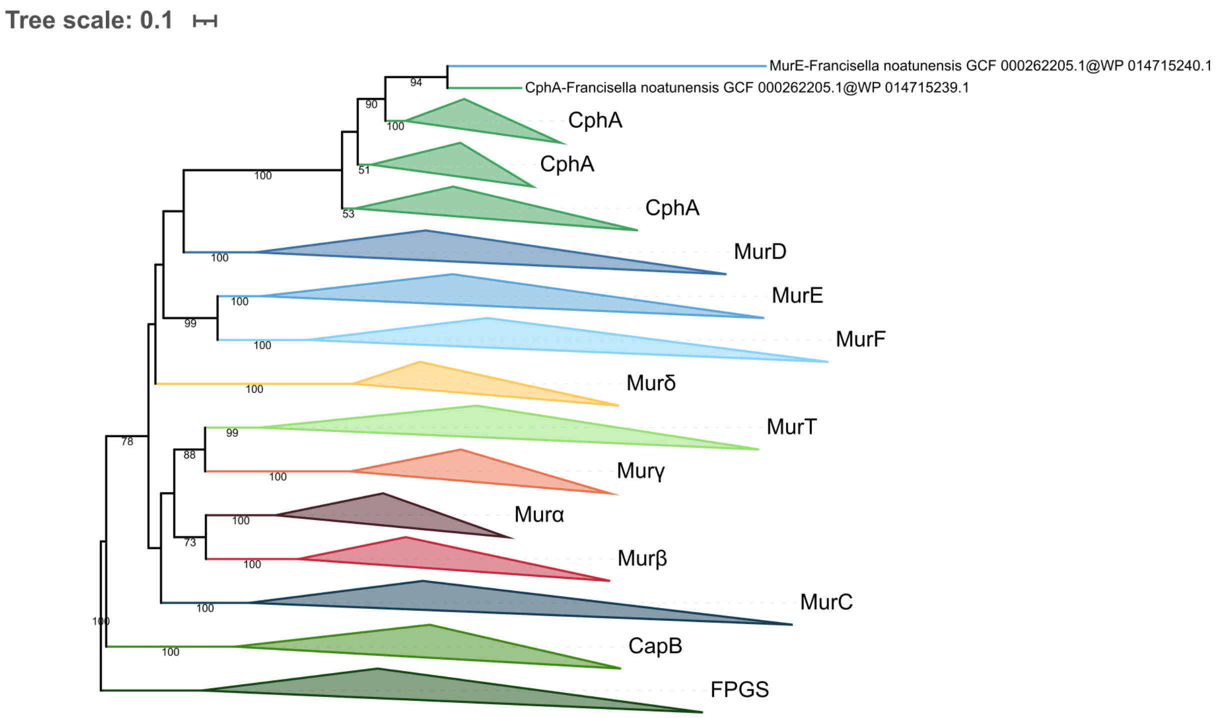


**Figure S13. Phylogenetic tree of the Mur domain-containing family rooted on FPGS.** The tree was inferred from a matrix of **3407 sequences x 550 unambiguously aligned AAs** using IQ-TREE **under the C40+G4 model**. Tree visualization was performed using iTOL. Bootstrap values are shown if greater or equal to 50. Branches were collapsed on homogeneous sequence annotation.

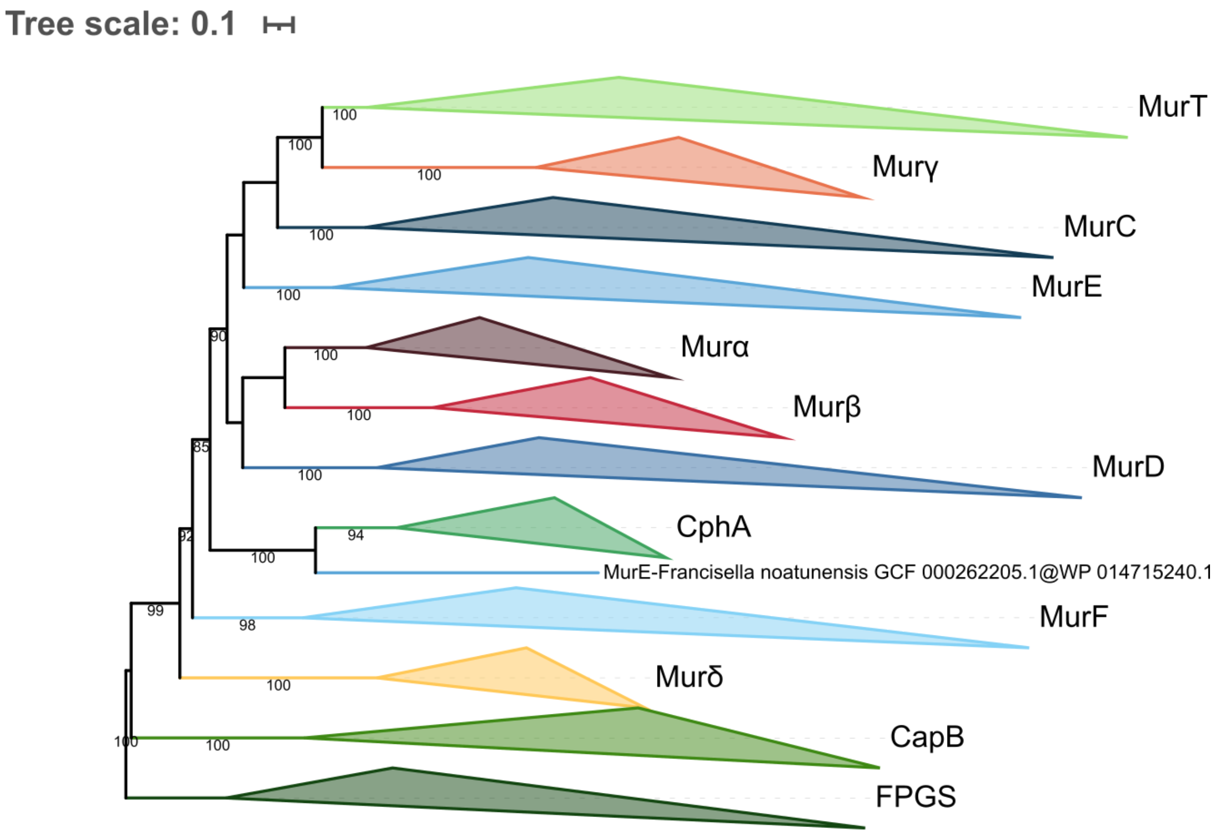


**Figure S14. Phylogenetic tree of the Mur domain-containing family rooted on FPGS.** The tree was inferred from a matrix of **3386 sequences x 228 unambiguously aligned AAs** using IQ-TREE **under the LG4X+R4 model**. Tree visualization was performed using iTOL. Bootstrap values are shown if greater or equal to 50. Branches were collapsed on homogeneous sequence annotation.

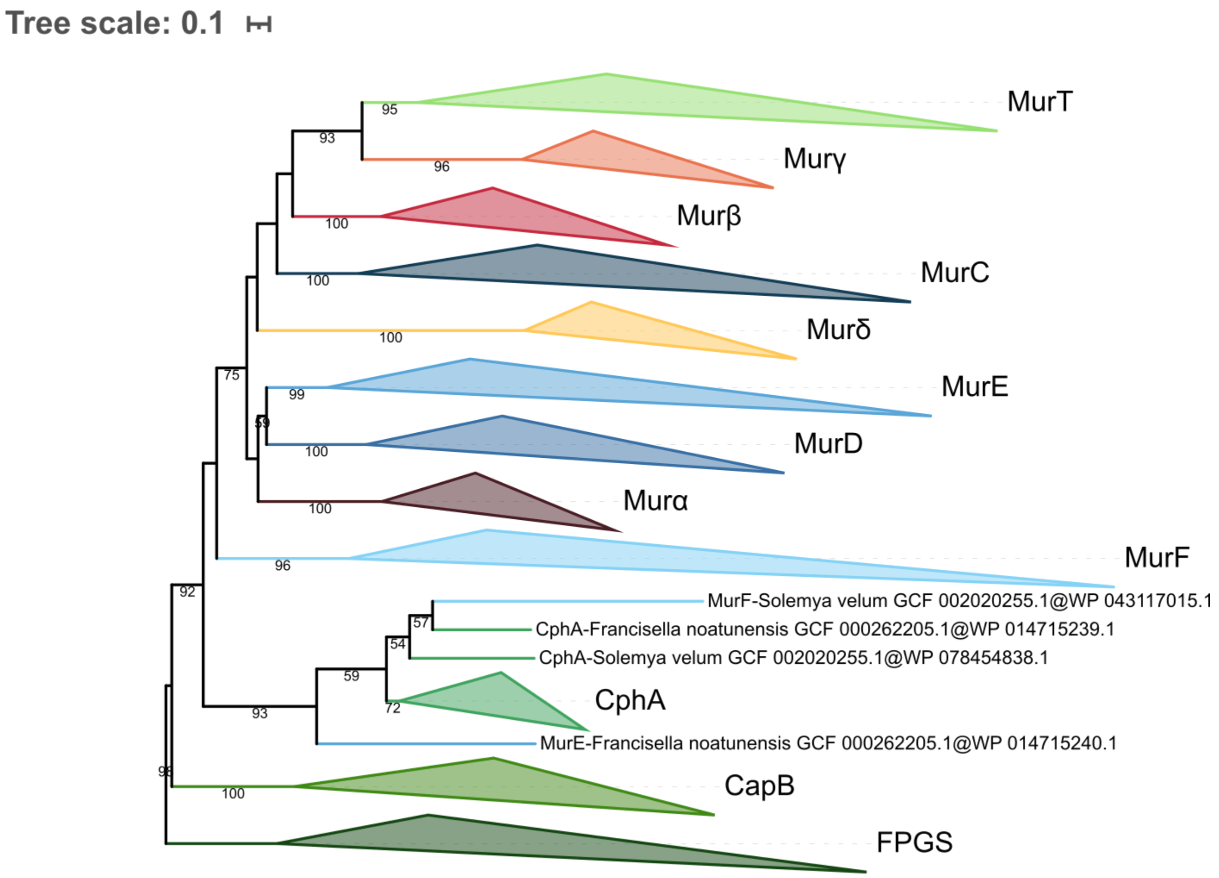


**Figure S15. Phylogenetic tree of the Mur domain-containing family rooted on FPGS.** The tree was inferred from a matrix of **3386 sequences x 228 unambiguously aligned AAs** using IQ-TREE **under the C20+G4 model**. Tree visualization was performed using iTOL. Bootstrap values are shown if greater or equal to 50. Branches were collapsed on homogeneous sequence annotation.

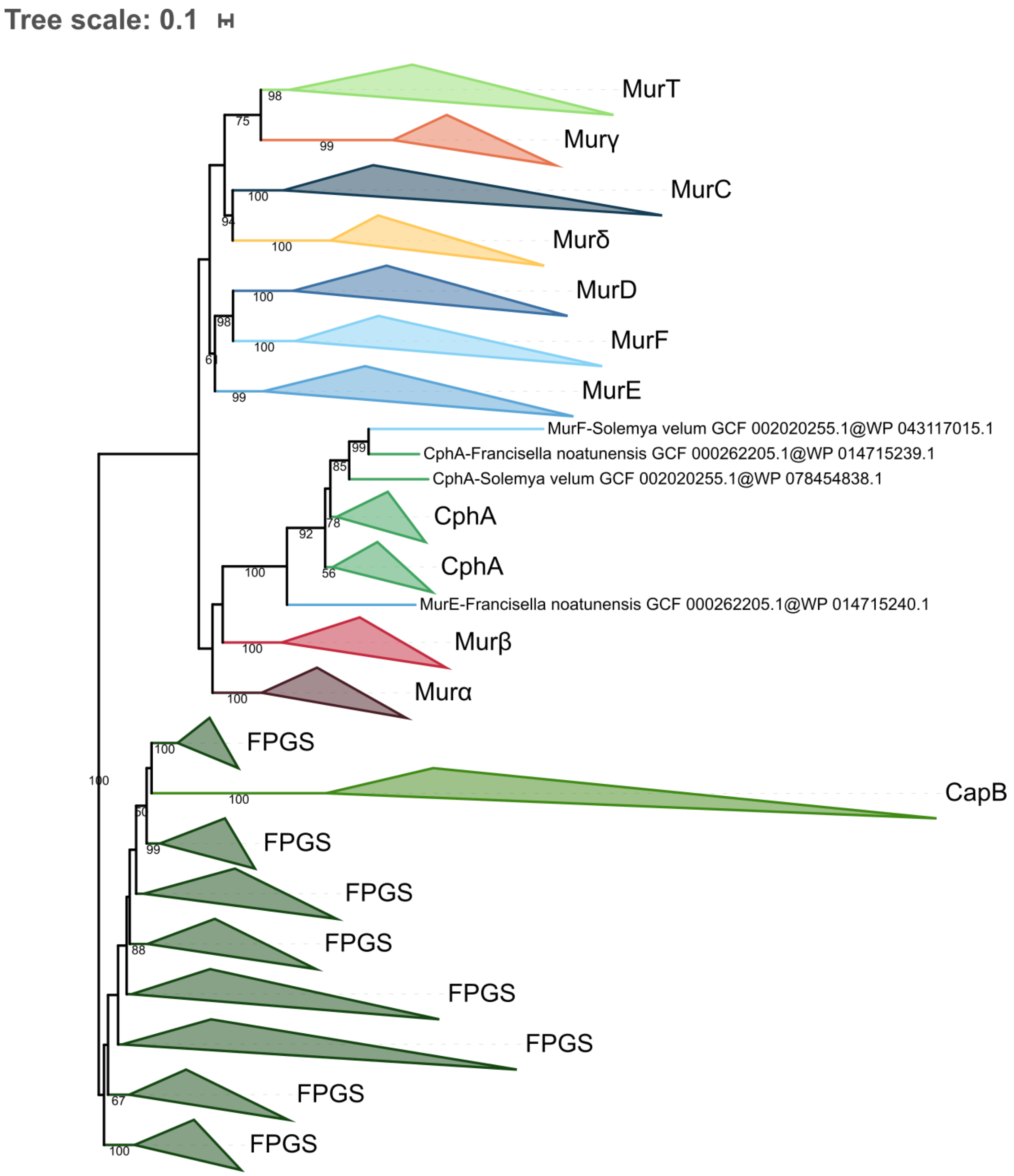


**Figure S16. Phylogenetic tree of the Mur domain-containing family rooted on FPGS.** The tree was inferred from a matrix of **3386 sequences x 228 unambiguously aligned AAs** using IQ-TREE **under the C40+G4 model**. Tree visualization was performed using iTOL. Bootstrap values are shown if greater or equal to 50. Branches were collapsed on homogeneous sequence annotation.

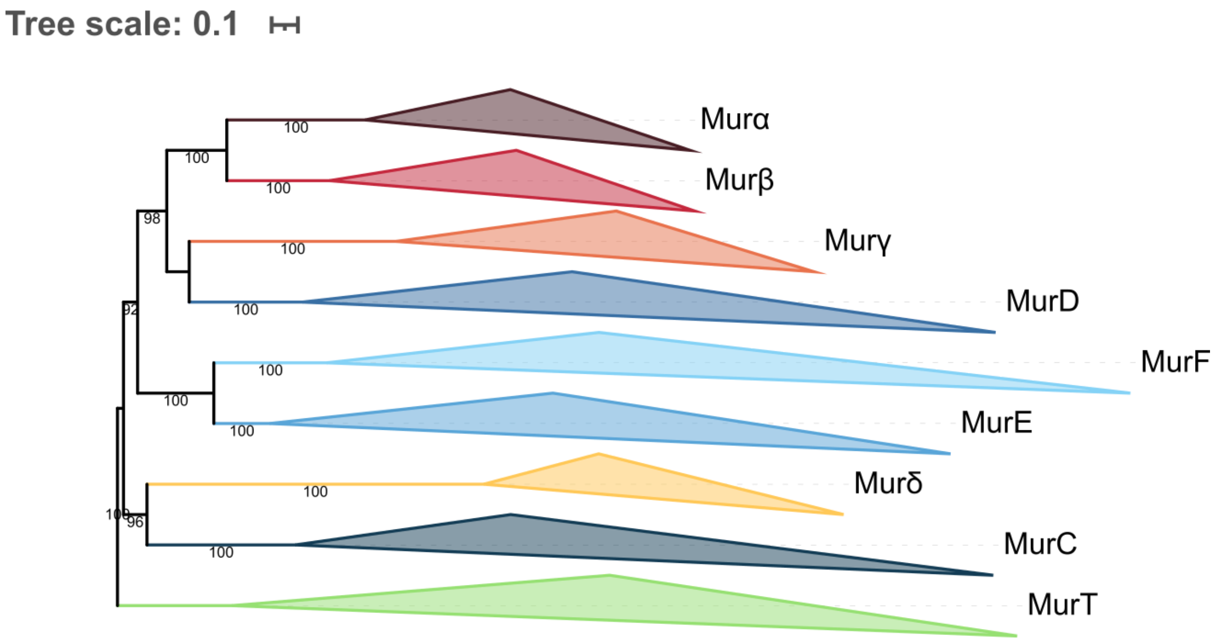


**Figure S17. Phylogenetic tree of the Mur domain-containing family rooted on MurT.** The tree was inferred from a matrix of **2677 sequences x 525 unambiguously aligned AAs** using IQ-TREE **under the LG4X+R4 model**. Tree visualization was performed using iTOL. Bootstrap values are shown if greater or equal to 50. Branches were collapsed on homogeneous sequence annotation.

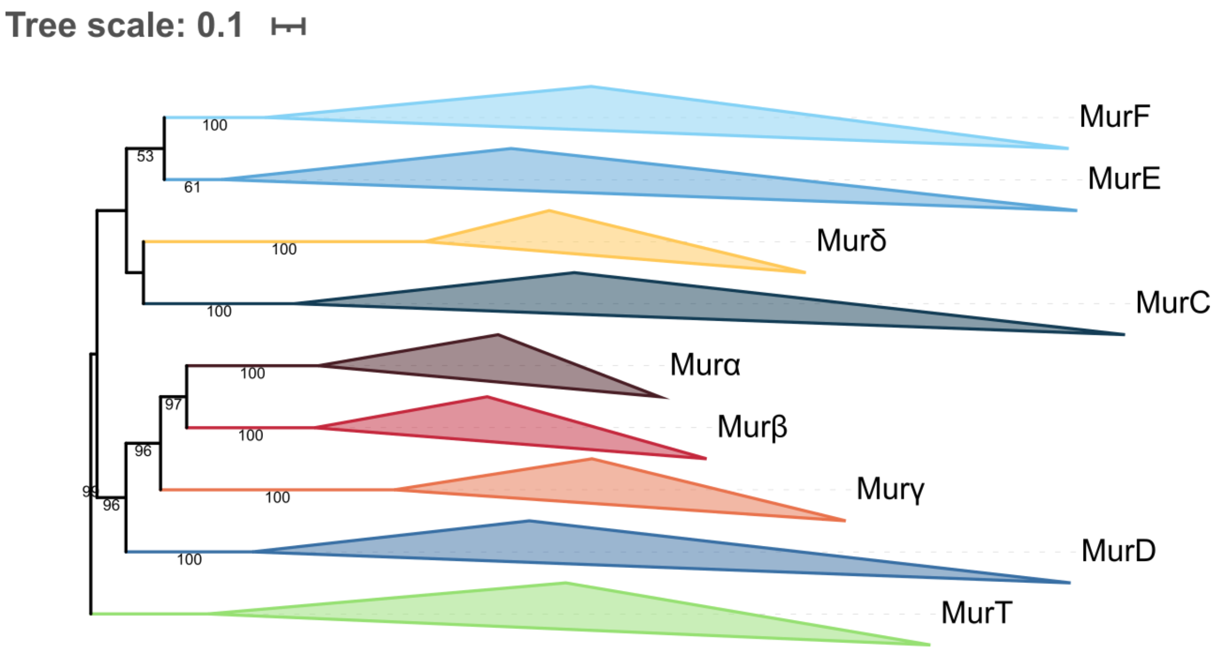


**Figure S18. Phylogenetic tree of the Mur domain-containing family rooted on MurT.** The tree was inferred from a matrix of **2677 sequences x 525 unambiguously aligned AAs** using IQ-TREE **under the C20+G4 model**. Tree visualization was performed using iTOL. Bootstrap values are shown if greater or equal to 50. Branches were collapsed on homogeneous sequence annotation.

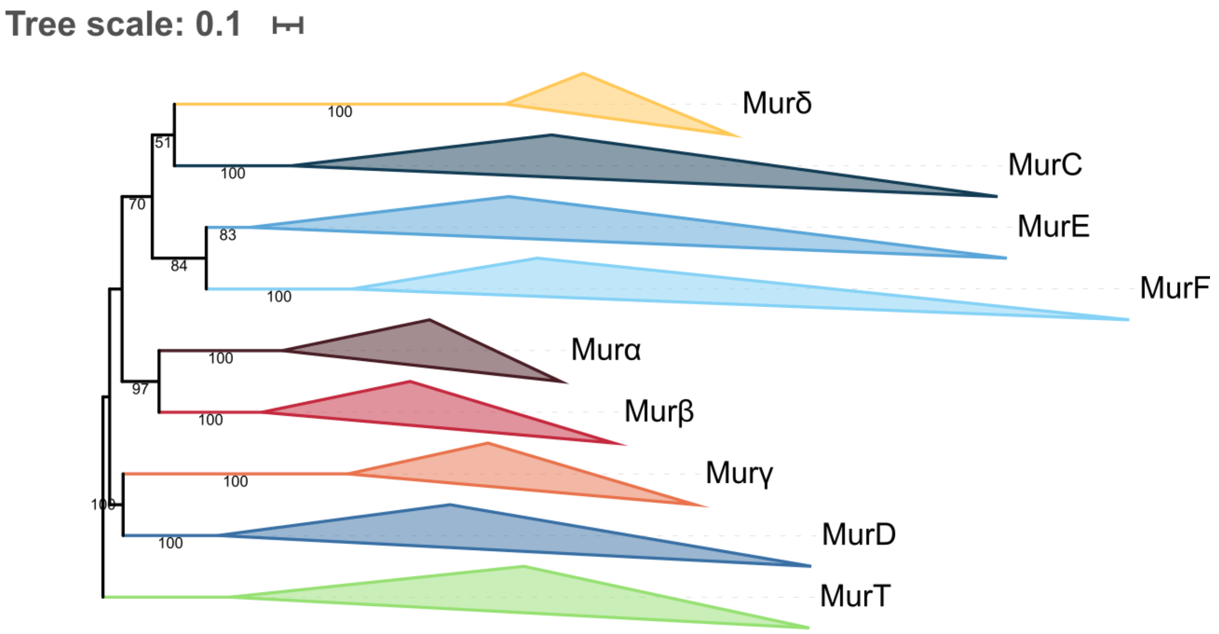


**Figure S19. Phylogenetic tree of the Mur domain-containing family rooted on MurT.** The tree was inferred from a matrix of **2677 sequences x 525 unambiguously aligned AAs** using IQ-TREE **under the C40+G4 model**. Tree visualization was performed using iTOL. Bootstrap values are shown if greater or equal to 50. Branches were collapsed on homogeneous sequence annotation.

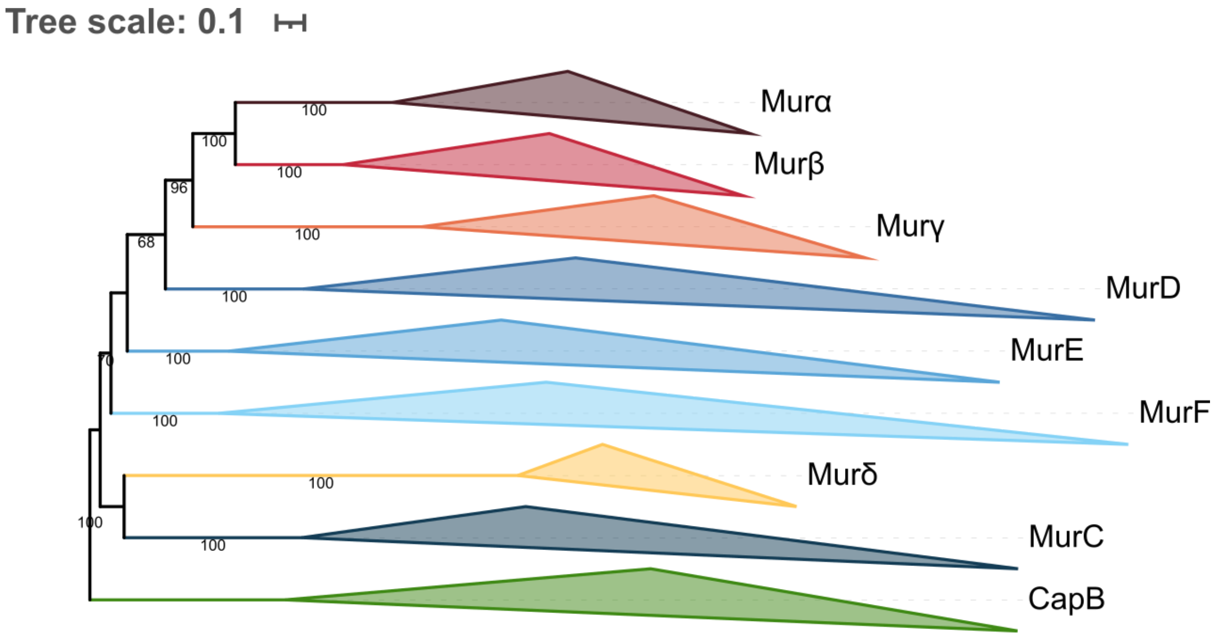


**Figure S20. Phylogenetic tree of the Mur domain-containing family rooted on CapB.** The tree was inferred from a matrix of **2519 sequences x 532 unambiguously aligned AAs** using IQ-TREE **under the LG4X+R4 model**. Tree visualization was performed using iTOL. Bootstrap values are shown if greater or equal to 50. Branches were collapsed on homogeneous sequence annotation.

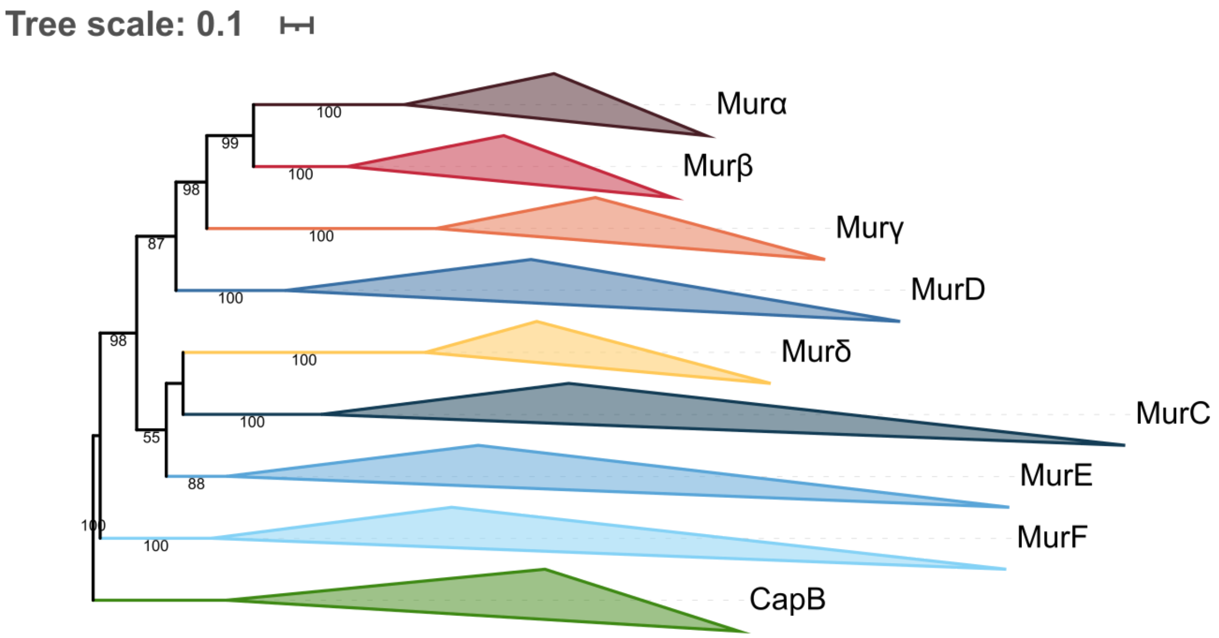


**Figure S21. Phylogenetic tree of the Mur domain-containing family rooted on CapB.** The tree was inferred from a matrix of **2519 sequences x 532 unambiguously aligned AAs** using IQ-TREE **under the C20+G4 model**. Tree visualization was performed using iTOL. Bootstrap values are shown if greater or equal to 50. Branches were collapsed on homogeneous sequence annotation.

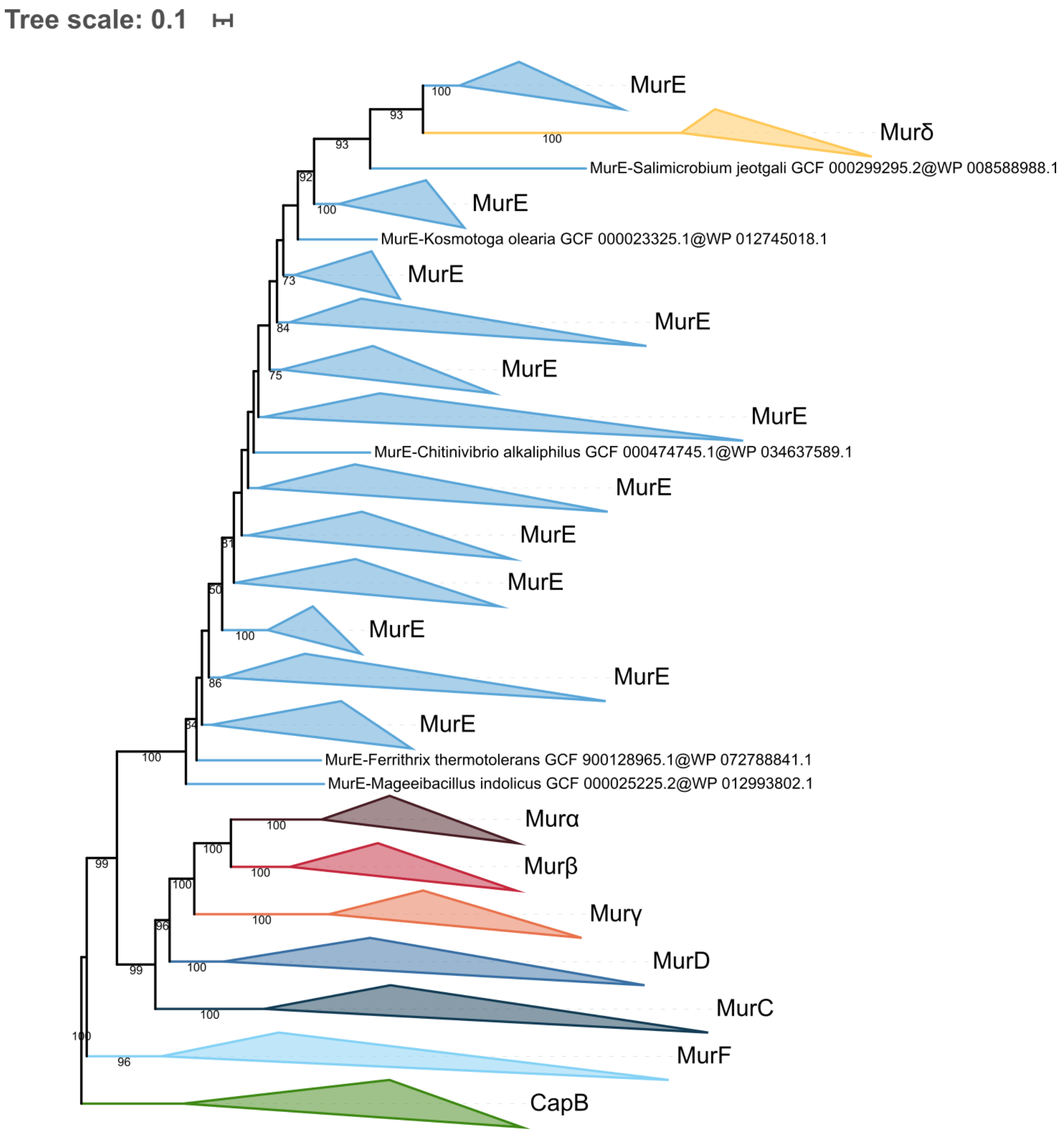


**Figure S22. Phylogenetic tree of the Mur domain-containing family rooted on CapB.** The tree was inferred from a matrix of **2519 sequences x 532 unambiguously aligned AAs** using IQ-TREE **under the C40+G4 model**. Tree visualization was performed using iTOL. Bootstrap values are shown if greater or equal to 50. Branches were collapsed on homogeneous sequence annotation.

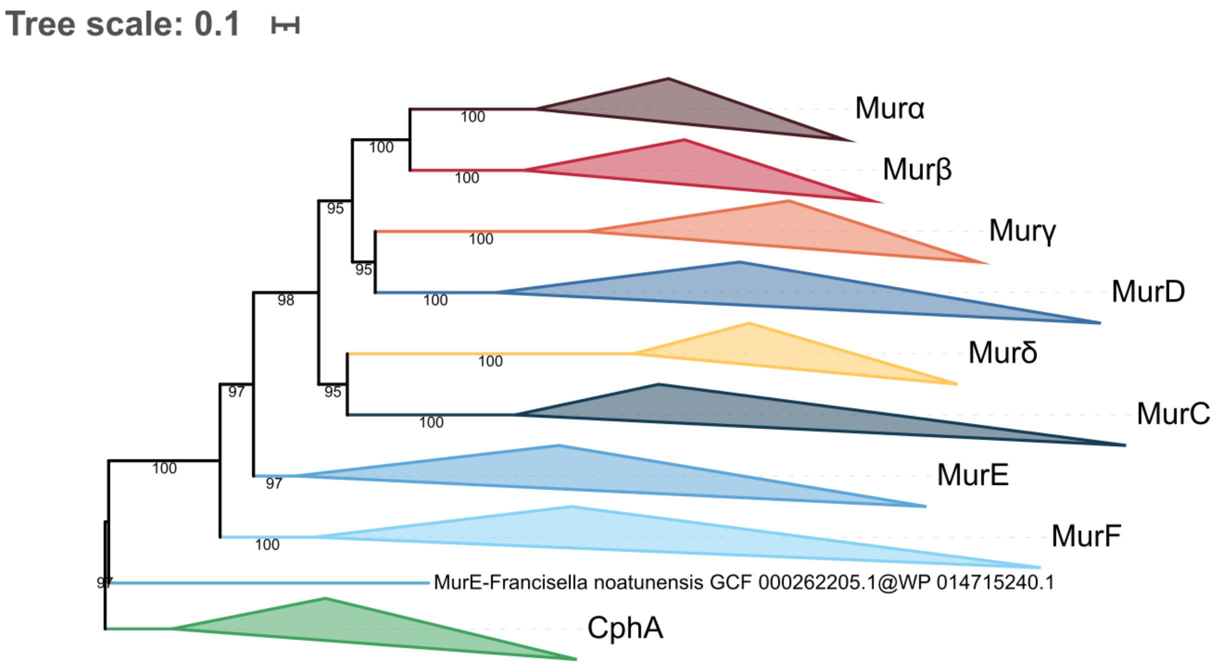


**Figure S23. Phylogenetic tree of the Mur domain-containing family rooted on CphA.** The tree was inferred from a matrix of **2461 sequences x 539 unambiguously aligned AAs** using IQ-TREE **under the LG4X+R4 model**. Tree visualization was performed using iTOL. Bootstrap values are shown if greater or equal to 50. Branches were collapsed on homogeneous sequence annotation.

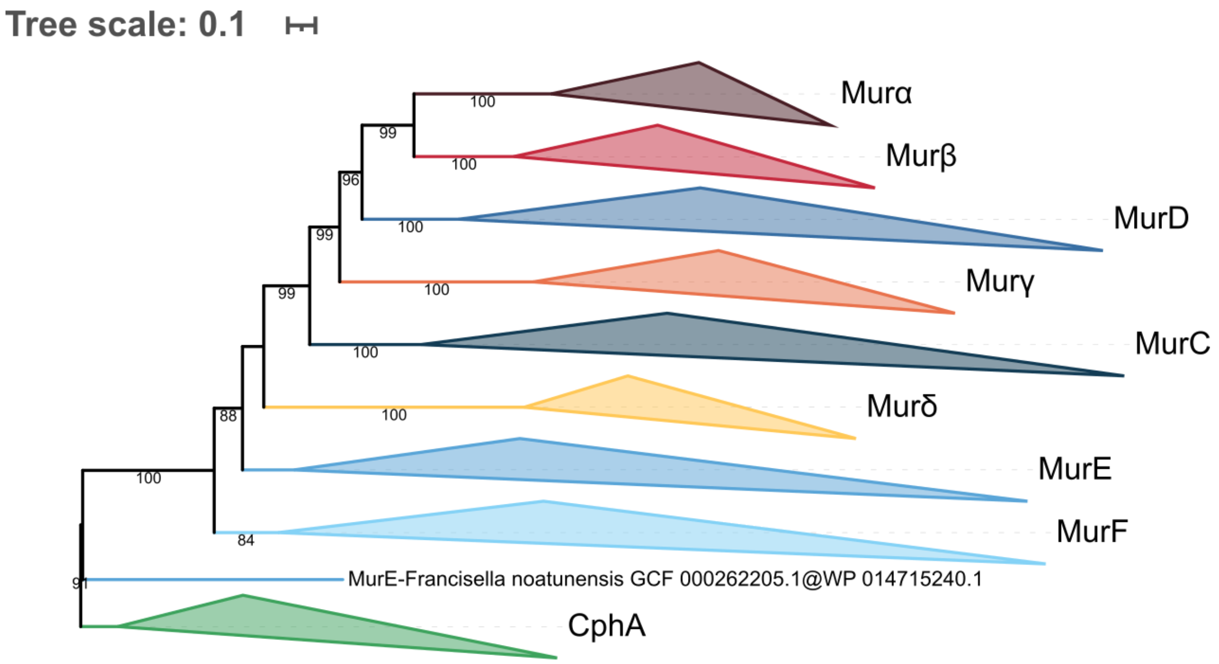


**Figure S24. Phylogenetic tree of the Mur domain-containing family rooted on CphA.** The tree was inferred from a matrix of **2461 sequences x 539 unambiguously aligned AAs** using IQ-TREE **under the C20+G4 model**. Tree visualization was performed using iTOL. Bootstrap values are shown if greater or equal to 50. Branches were collapsed on homogeneous sequence annotation.

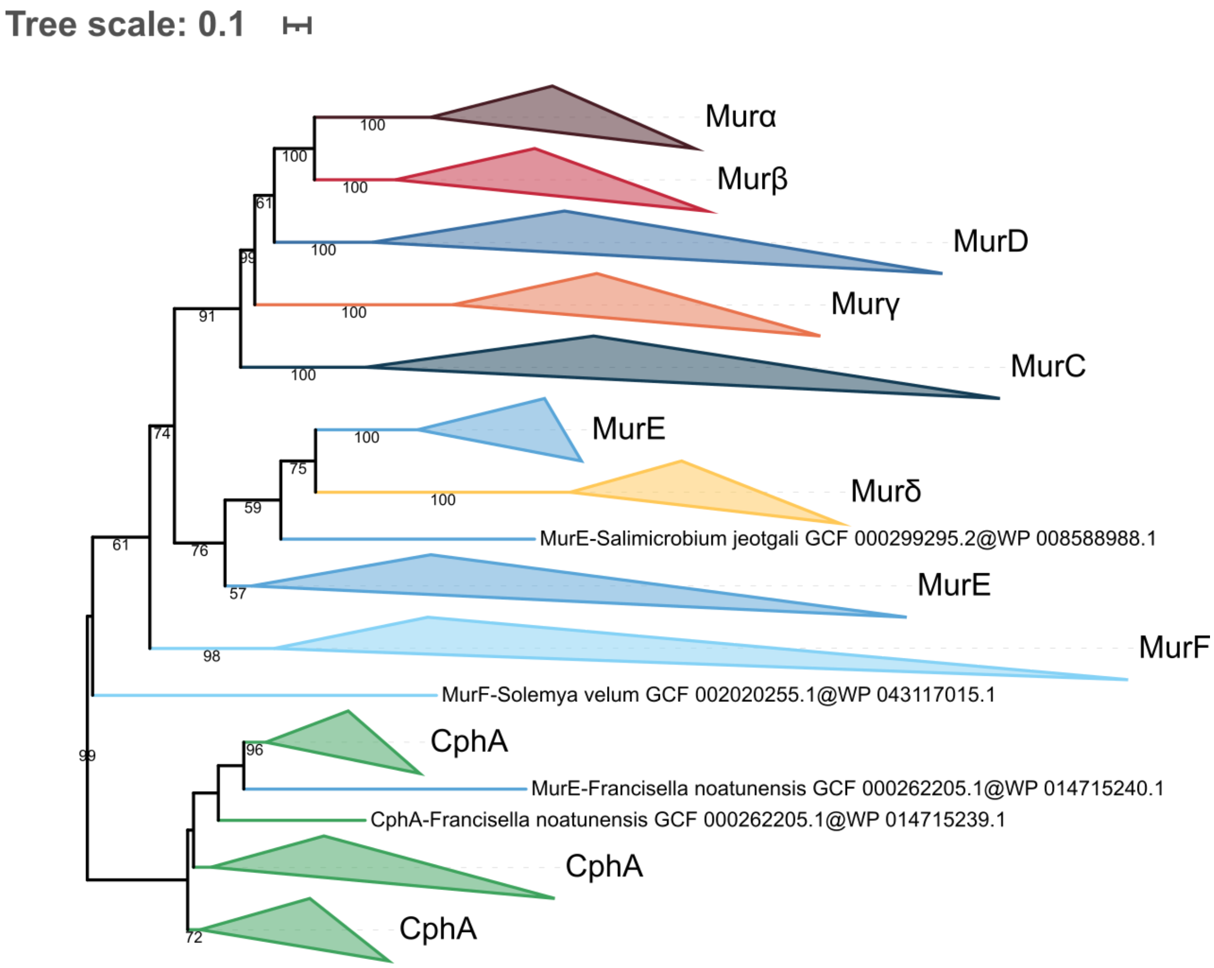


**Figure S25. Phylogenetic tree of the Mur domain-containing family rooted on CphA.** The tree was inferred from a matrix of **2461 sequences x 539 unambiguously aligned AAs** using IQ-TREE **under the C40+G4 model**. Tree visualization was performed using iTOL. Bootstrap values are shown if greater or equal to 50. Branches were collapsed on homogeneous sequence annotation.

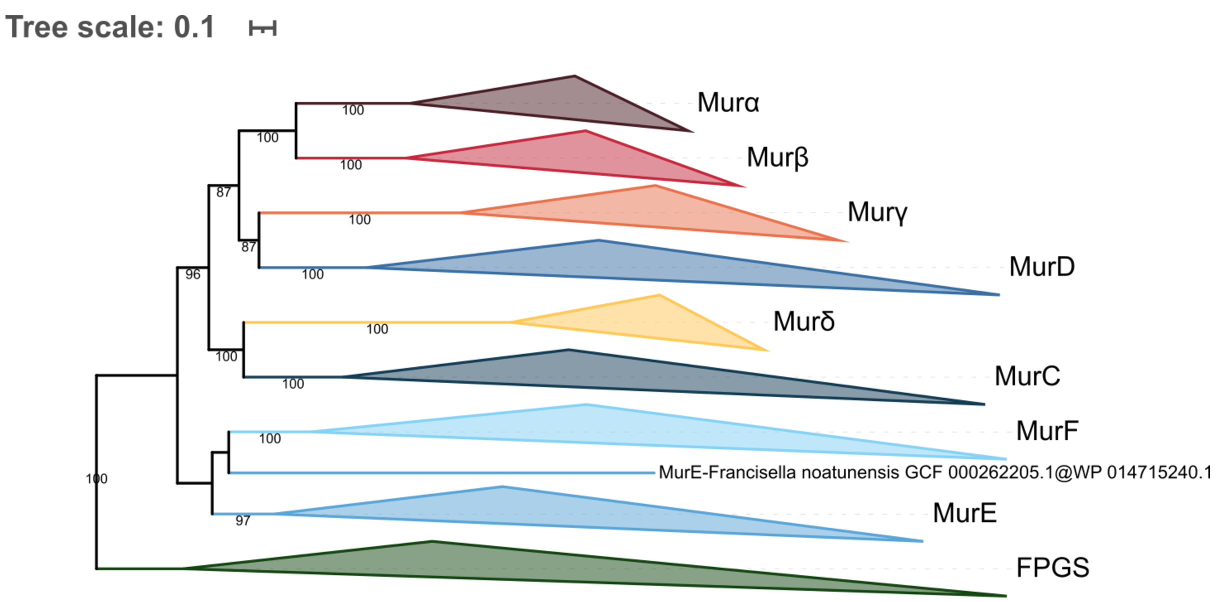


**Figure S26. Phylogenetic tree of the Mur domain-containing family rooted on FPGS.** The tree was inferred from a matrix of **3,046 sequences x 543 unambiguously aligned AAs** using IQ-TREE **under the LG4X+R4 model**. Tree visualization was performed using iTOL. Bootstrap values are shown if greater or equal to 50. Branches were collapsed on homogeneous sequence annotation.

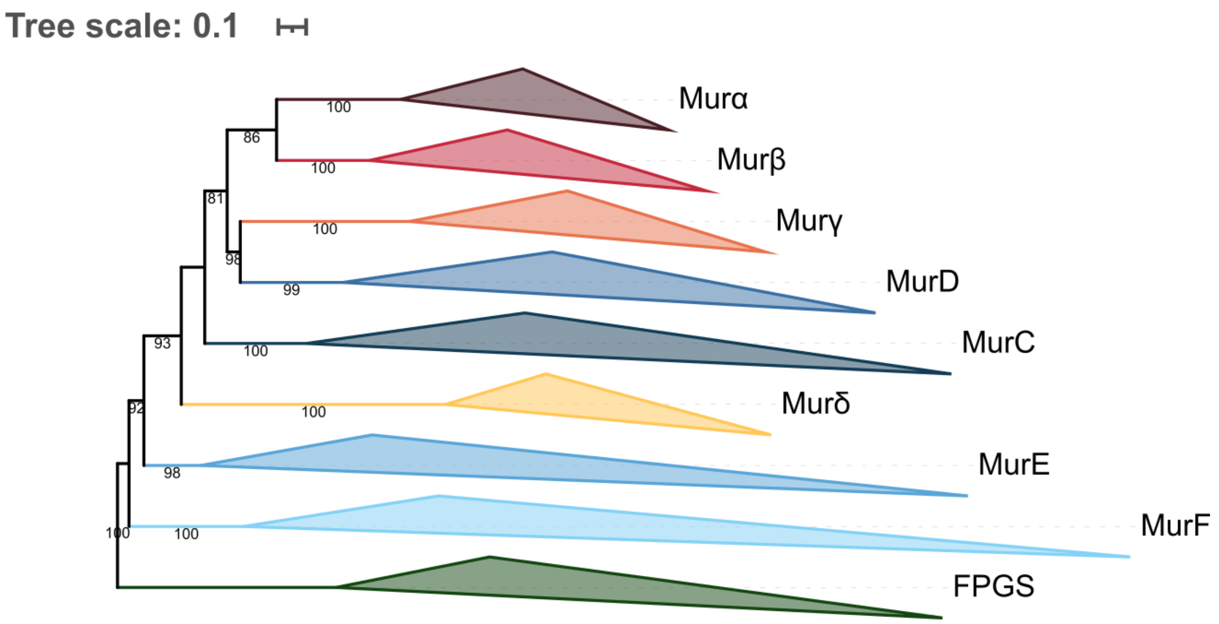


**Figure S27. Phylogenetic tree of the Mur domain-containing family rooted on FPGS.** The tree was inferred from a matrix of **3,046 sequences x 543 unambiguously aligned AAs** using IQ-TREE **under the C20+G4 model**. Tree visualization was performed using iTOL. Bootstrap values are shown if greater or equal to 50. Branches were collapsed on homogeneous sequence annotation.

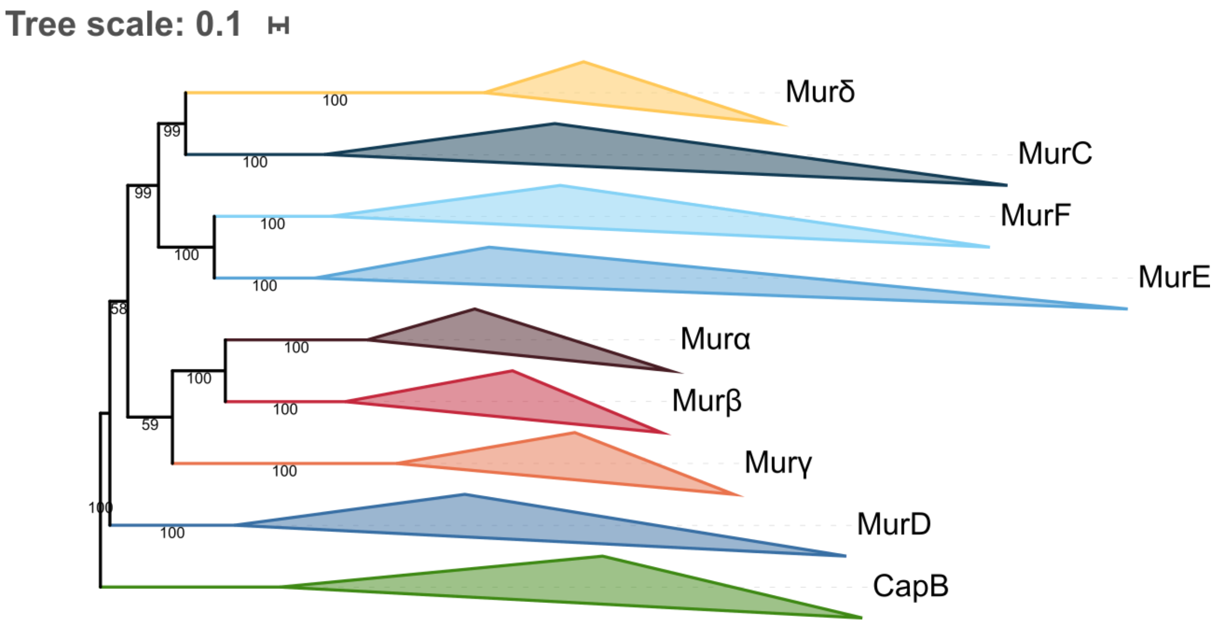


**Figure S28. Phylogenetic tree of the Mur domain-containing family rooted on CapB.** The tree was inferred from a matrix of **2519 sequences x 532 unambiguously aligned AAs** using IQ-TREE **under the PMSF LG+C60+G4 model** with 3000 iterations. The guide tree was the C40+G4 tree of Figure S22. Tree visualization was performed using iTOL. Bootstrap values are shown if greater or equal to 50. Branches were collapsed on homogeneous sequence annotation.

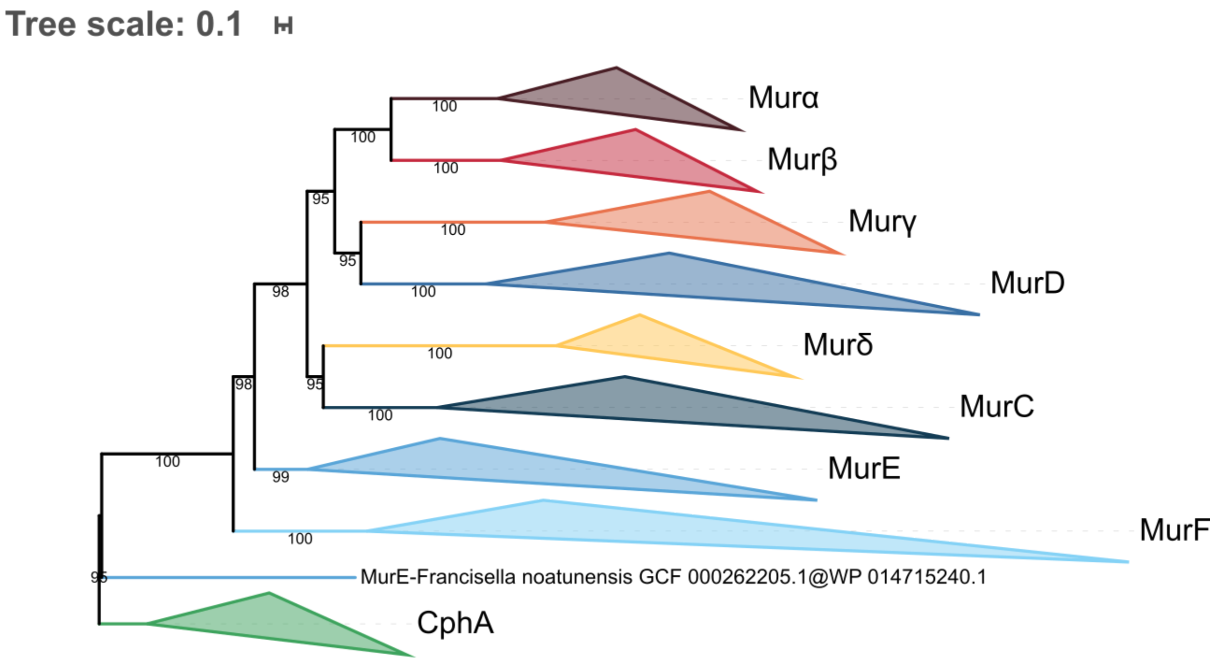


**Figure S29. Phylogenetic tree of the Mur domain-containing family rooted on CphA.** The tree was inferred from a matrix of **2461 sequences x 539 unambiguously aligned AAs** using IQ-TREE **under the PMSF LG+C60+G4 model** with 3000 iterations. The guide tree was the C40+G4 tree of Figure S25. Tree visualization was performed using iTOL. Bootstrap values are shown if greater or equal to 50. Branches were collapsed on homogeneous sequence annotation.

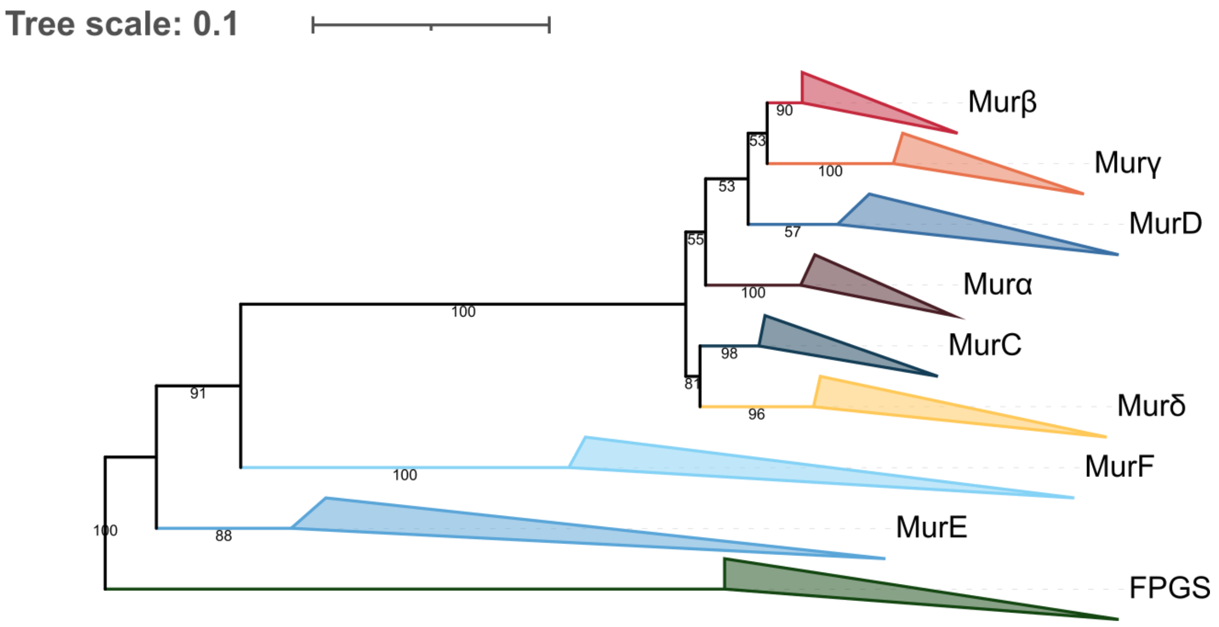


**Figure S30. Phylogenetic tree of the Mur domain-containing family rooted on FPGS. Indels tree** inferred from a matrix of **3001 sequences x 1799 unambiguously aligned AAs** using RAxML **under the BINGAMMAX model**. Tree visualization was performed using iTOL. Bootstrap values are shown if greater or equal to 50. Branches were collapsed on homogeneous sequence annotation.

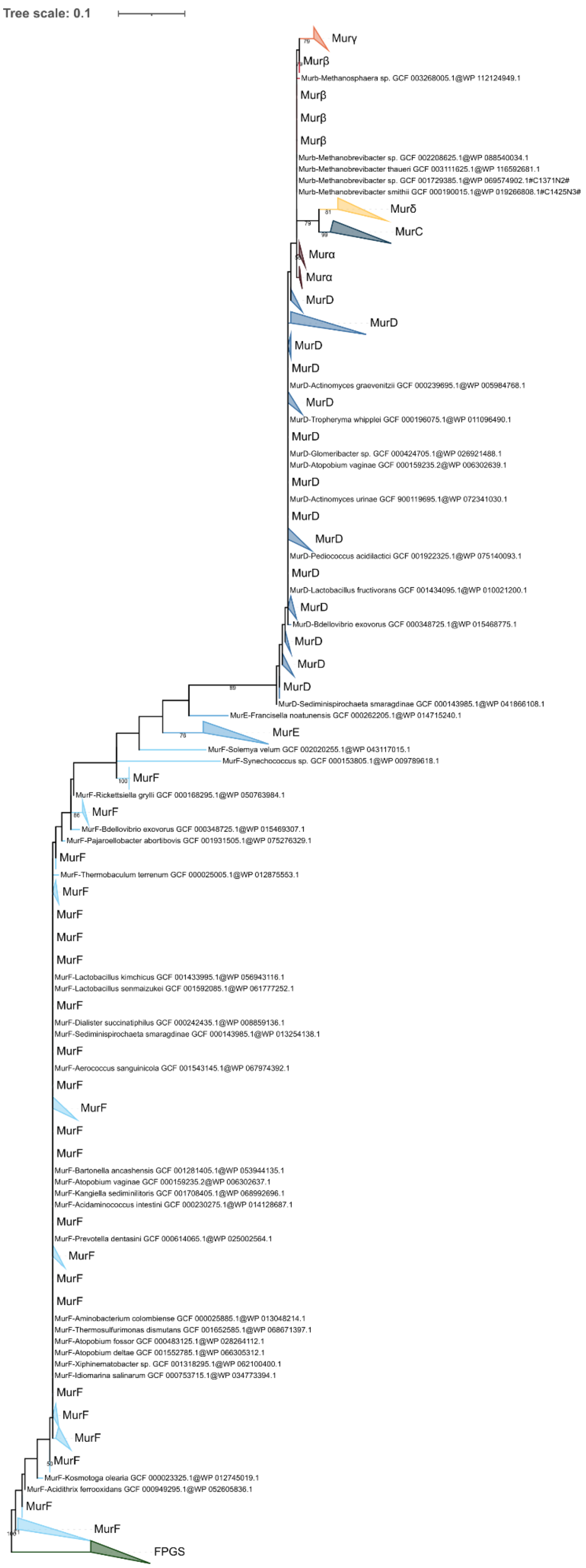
Tree scale: 0.1
Murγ
Murβ
Murb-Methanosphaera sp. GCF 003268005.1@WP 112124949.1
Murβ
Murβ
Murβ
Murb-Methanobrevibacter sp. GCF 002208625.1@WP 088540034.1
Murb-Methanobrevibacter thaueri GCF 003111625.1@WP 116592681.1
Murb-Methanobrevibacter sp. GCF 001729385.1@WP 069574902.1#C1371N2#
Murb-Methanobrevibacter smithii GCF 000190015.1@WP 019266808.1#C1425N3#
Murδ
MurC
Murα
Murα
MurD
MurD
MurD
MurD
MurD-Actinomyces graevenitzii GCF 000239695.1@WP 005984768.1
MurD
MurD-Tropheryma whipplei GCF 000196075.1@WP 011096490.1
MurD
MurD-Glomeribacter sp. GCF 000424705.1@WP 026921488.1
MurD-Atopobium vaginae GCF 000159235.2@WP 006302639.1
MurD
MurD-Actinomyces urinae GCF 900119695.1@WP 072341030.1
MurD
MurD
MurD-Pediococcus acidilactici GCF 001922325.1@WP 075140093.1
MurD
MurD-Lactobacillus fructivorans GCF 001434095.1@WP 010021200.1
MurD
MurD-Bdellovibrio exovorus GCF 000348725.1@WP 015468775.1
MurD
MurD
MurD
MurD-Sediminispirochaeta smaragdinae GCF 000143985.1@WP 041866108.1
MurE-Francisella noatunensis GCF 000262205.1@WP 014715240.1
MurE
MurF-Solemya velum GCF 002020255.1@WP 043117015.1
MurF-Synechococcus sp. GCF 000153805.1@WP 009789618.1
MurF
MurF-Rickettsiella grylli GCF 000168295.1@WP 050763984.1
MurF
MurF-Bdellovibrio exovorus GCF 000348725.1@WP 015469307.1
MurF-Pajaroellobacter abortibovis GCF 001931505.1@WP 075276329.1
MurF
MurF-Thermobaculum terrenum GCF 000025005.1@WP 012875553.1
MurF
MurF
MurF
MurF
MurF-Lactobacillus kimchicus GCF 001433995.1@WP 056943116.1
MurF-Lactobacillus senmaizukei GCF 001592085.1@WP 061777252.1
MurF
MurF-Dialister succinatiphilus GCF 000242435.1@WP 008859136.1
MurF-Sediminispirochaeta smaragdinae GCF 000143985.1@WP 013254138.1
MurF
MurF-Aerococcus sanguinicola GCF 001543145.1@WP 067974392.1
MurF
MurF
MurF
MurF
MurF-Bartonella ancashensis GCF 001281405.1@WP 053944135.1
MurF-Atopobium vaginae GCF 000159235.2@WP 006302637.1
MurF-Kangiella sediminilitoris GCF 001708405.1@WP 068992696.1
MurF-Acidaminococcus intestini GCF 000230275.1@WP 014128687.1
MurF
MurF-Prevotella dentasini GCF 000614065.1@WP 025002564.1
MurF
MurF
MurF
MurF-Aminobacterium colombiense GCF 000025885.1@WP 013048214.1
MurF-Thermosulfurimonas dismutans GCF 001652585.1@WP 068671397.1
MurF-Atopobium fossor GCF 000483125.1@WP 028264112.1
MurF-Atopobium deltae GCF 001552785.1@WP 066305312.1
MurF-Xiphinematobacter sp. GCF 001318295.1@WP 062100400.1
MurF-Idiomarina salinarum GCF 000753715.1@WP 034773394.1
MurF
MurF
MurF
MurF
MurF-Kosmotoga olearia GCF 000023325.1@WP 012745019.1
MurF-Acidithrix ferrooxidans GCF 000949295.1@WP 052605836.1
MurF
MurF
FPGS

**Figure S31. Phylogenetic tree of the Mur domain-containing family rooted on FPGS. Indels tree** inferred from a matrix of **3004 sequences x 281 unambiguously aligned AAs** using RAxML **under the BINGAMMAX model**. Tree visualization was performed using iTOL. Bootstrap values are shown if greater or equal to 50. Branches were collapsed on homogeneous sequence annotation.

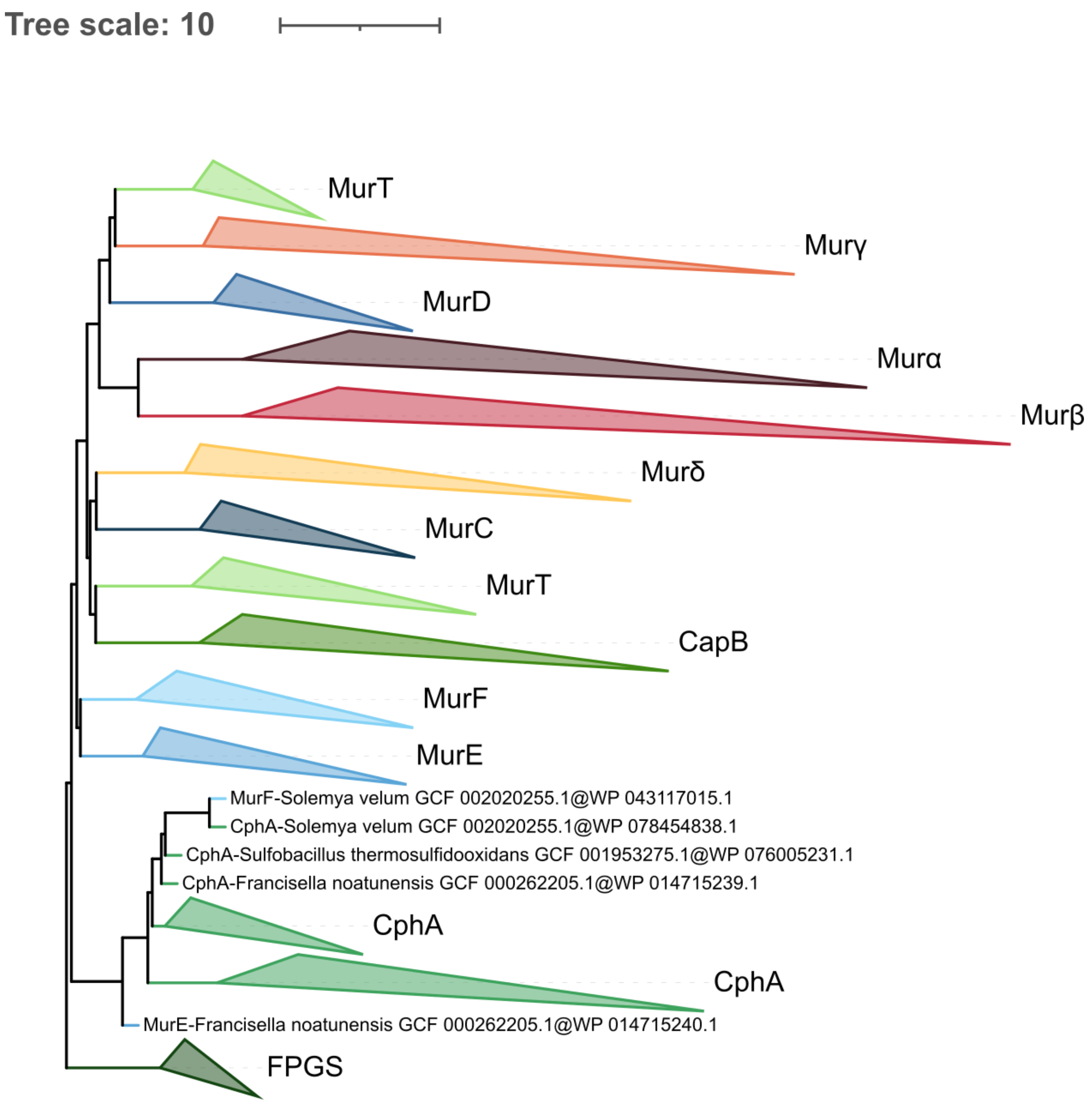


**Figure S32. Phylogenetic tree of the Mur domain-containing family rooted on FPGS. ASTRAL tree** inferred computed **from the 1000 LG4X+R4 species resampling trees**. Tree visualization was performed using iTOL. Branches were collapsed on homogeneous sequence annotation.

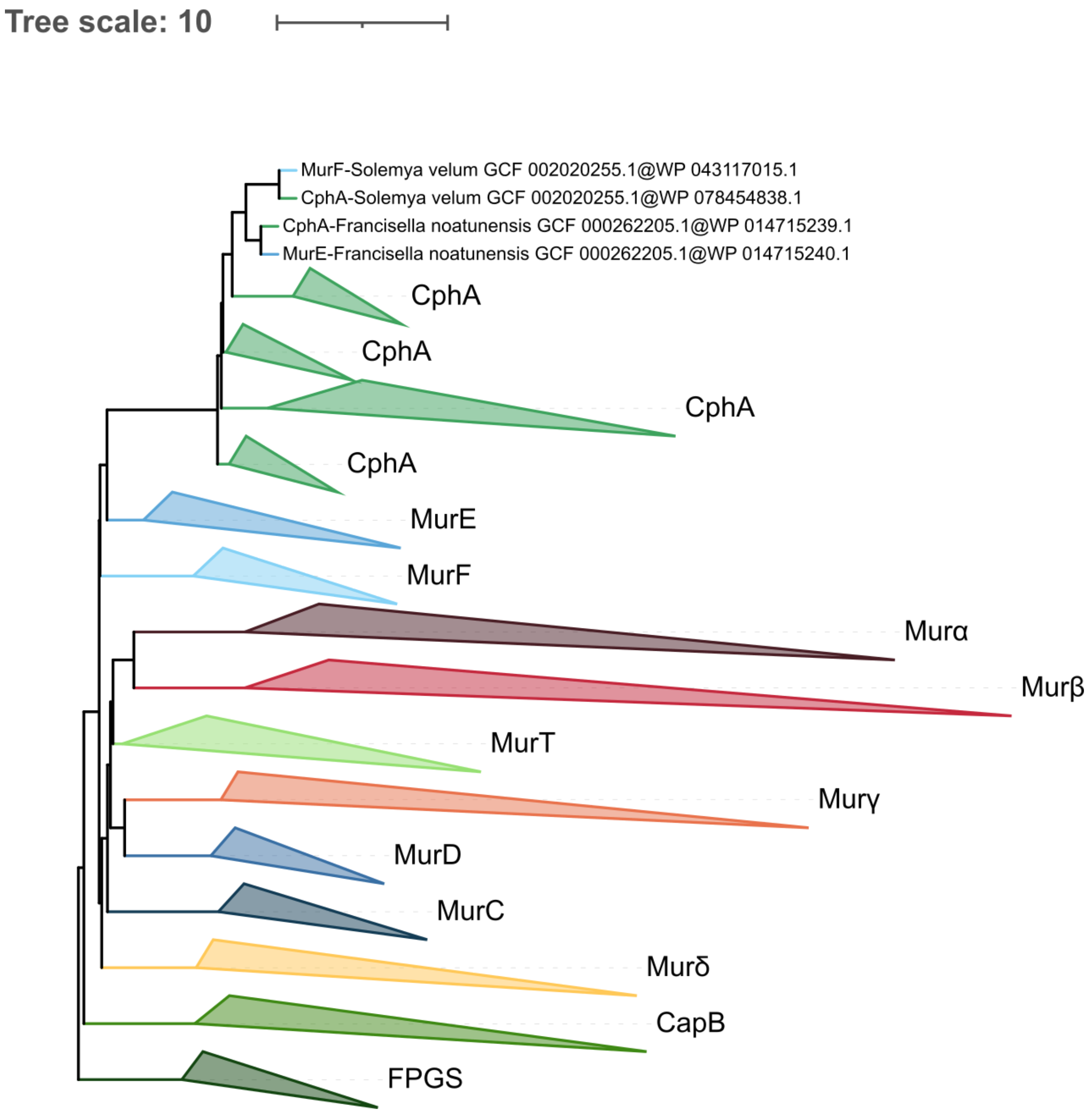


**Figure S33. Phylogenetic tree of the Mur domain-containing family rooted on FPGS. ASTRAL tree** computed **from the 1000 C20+G4 species resampling trees**. Tree visualization was performed using iTOL. Branches were collapsed on homogeneous sequence annotation.

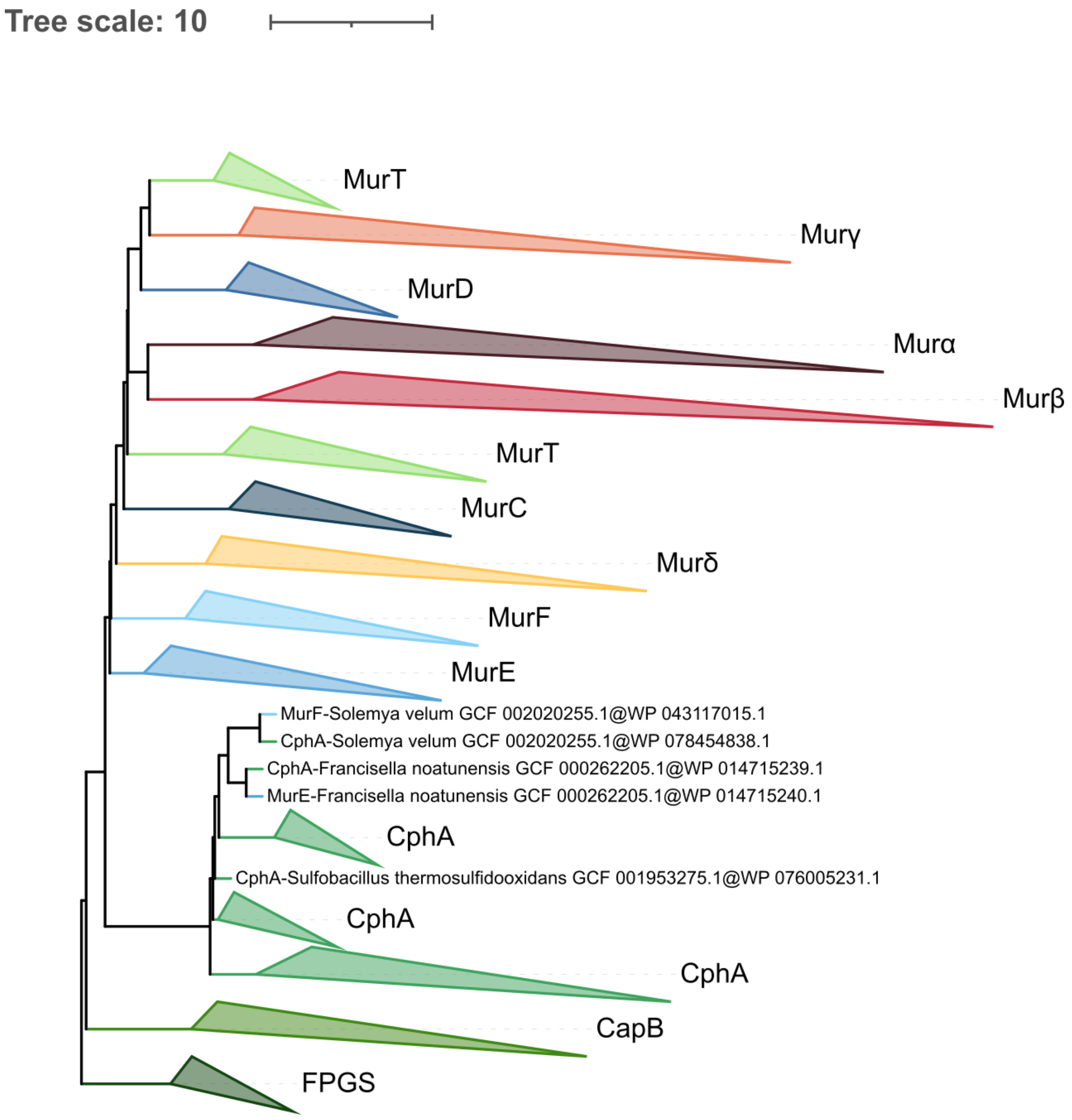


**Figure S34. Phylogenetic tree of the Mur domain-containing family rooted on FPGS. ASTRAL tree** computed **from the 1000 C40+G4 species resampling trees**. Tree visualization was performed using iTOL. Branches were collapsed on homogeneous sequence annotation.

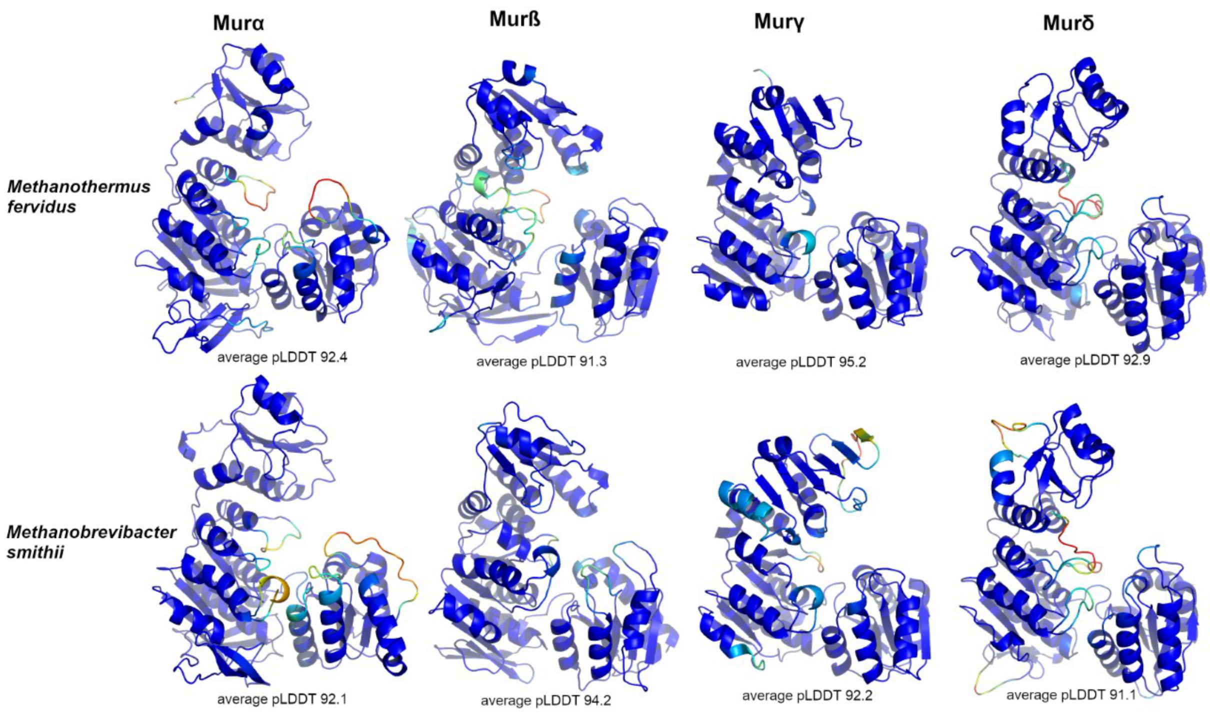


**Figure S35. Cartoon representation of the AlphaFold models of the Murαßγδ enzymes from *M. fervidus* and *M. smithii***. Amino acids are colored according to their pLDDT value (from red <50 to blue >90). The average pLDDT value calculated for the entire protein is indicated below each model. Because our analysis is centered on the C-terminal domain, we selected the model with the highest average pLDDT for this specific domain among the five models generated by AlphaFold. For Murα and Murß from *M. fervidus* and Murδ from *M. smithii*., these models had a slightly lower total average pLDDT (less than 1%) compared to the overall best.

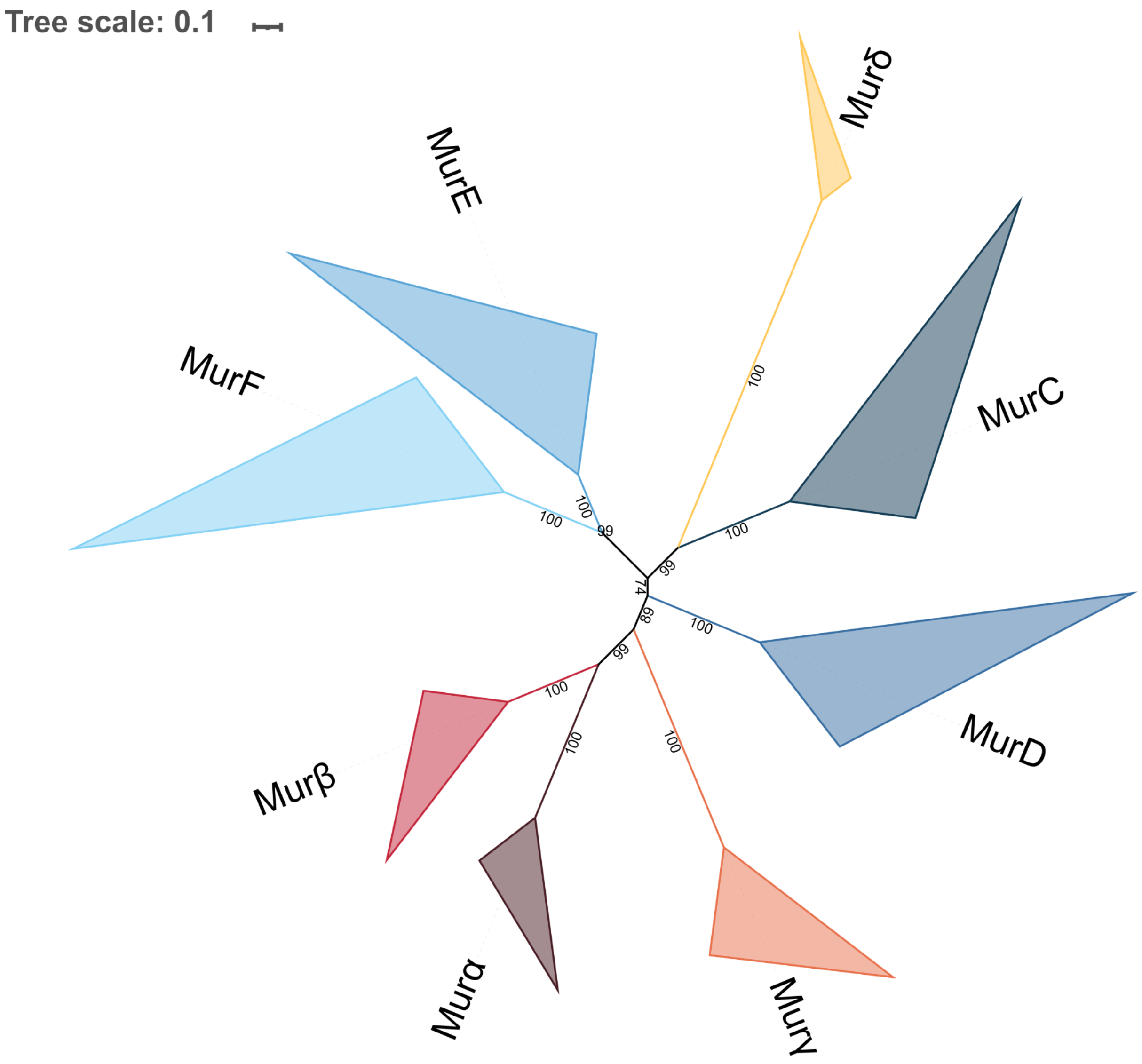


**Figure S36. Unrooted phylogenetic tree of the Mur domain-containing family.** The tree was inferred from a matrix of **2432 sequences x 528 unambiguously aligned AAs** using IQ-TREE **under the LG4X+R4 model**. Tree visualization was performed using iTOL. Bootstrap values are shown if greater or equal to 50. Branches were collapsed on homogeneous sequence annotation.

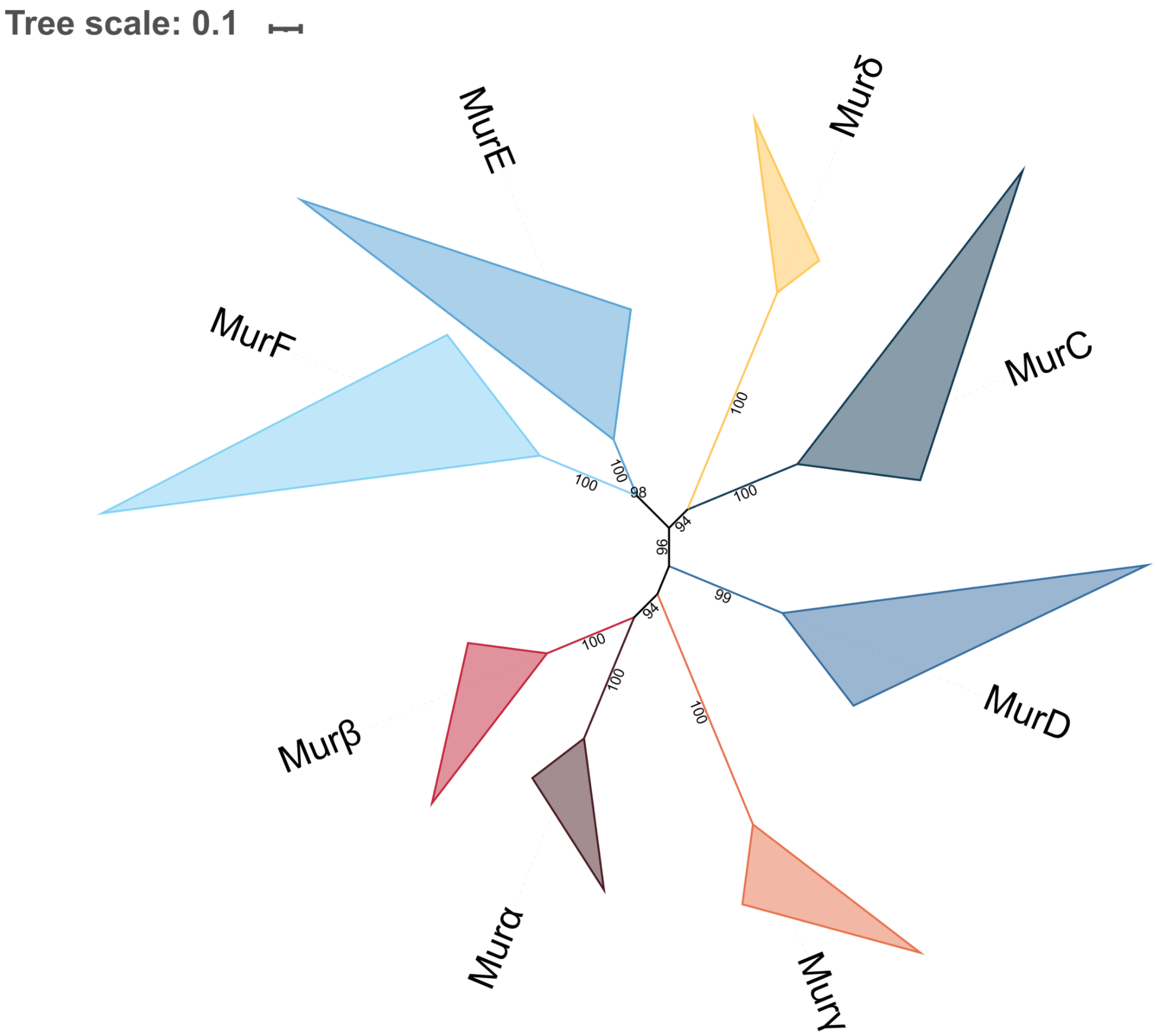


**Figure S37. Unrooted phylogenetic tree of the Mur domain-containing family.** The tree was inferred from a matrix of **2432 sequences x 528 unambiguously aligned AAs** using IQ-TREE **under the C20+G4 model**. Tree visualization was performed using iTOL. Bootstrap values are shown if greater or equal to 50. Branches were collapsed on homogeneous sequence annotation.

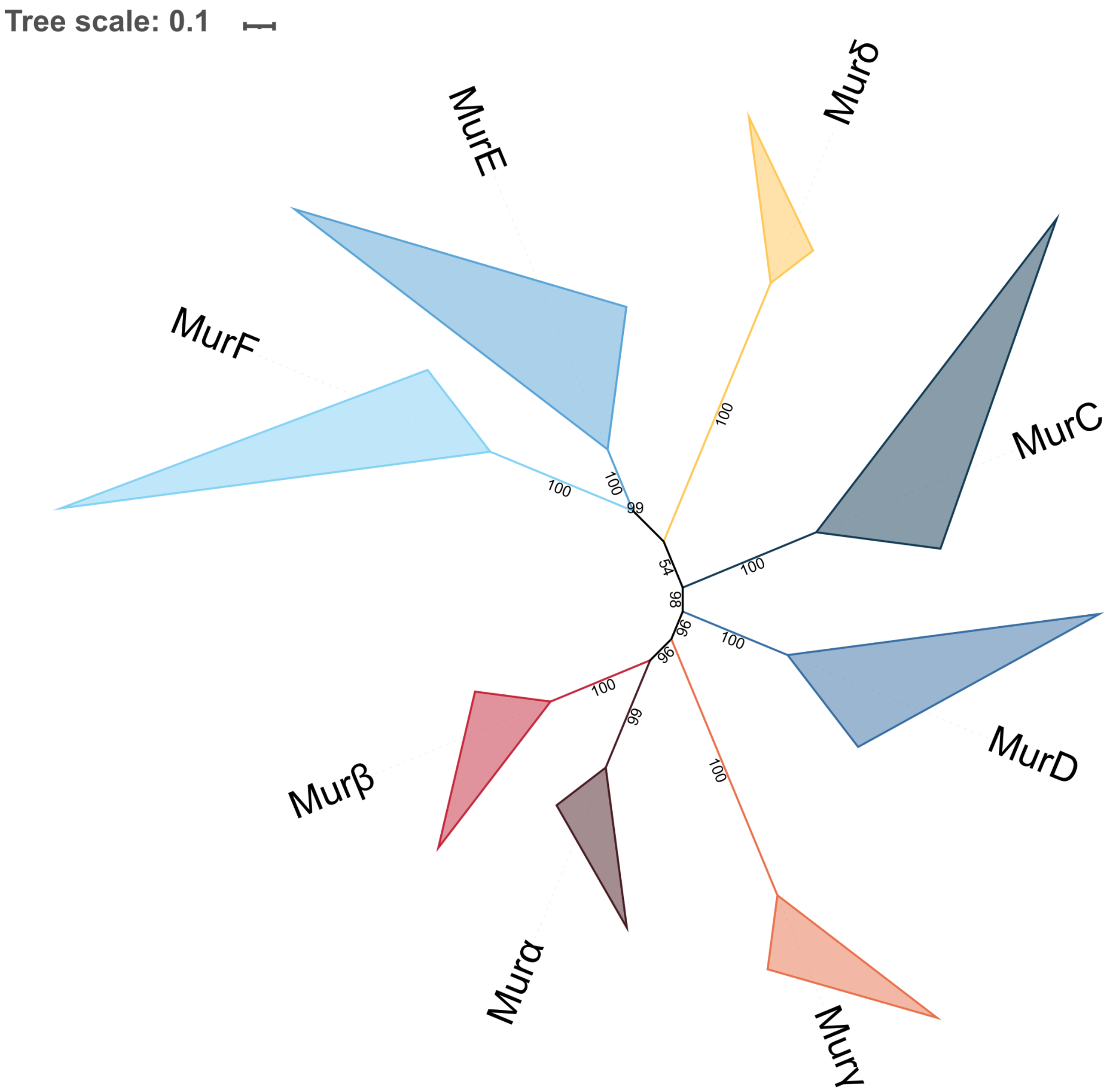


**Figure S38. Unrooted phylogenetic tree of the Mur domain-containing family.** The tree was inferred from a matrix of 2**432 sequences x 528 unambiguously aligned AAs** using IQ-TREE **under the C40+G4 model**. Tree visualization was performed using iTOL. Bootstrap values are shown if greater or equal to 50. Branches were collapsed on homogeneous sequence annotation.

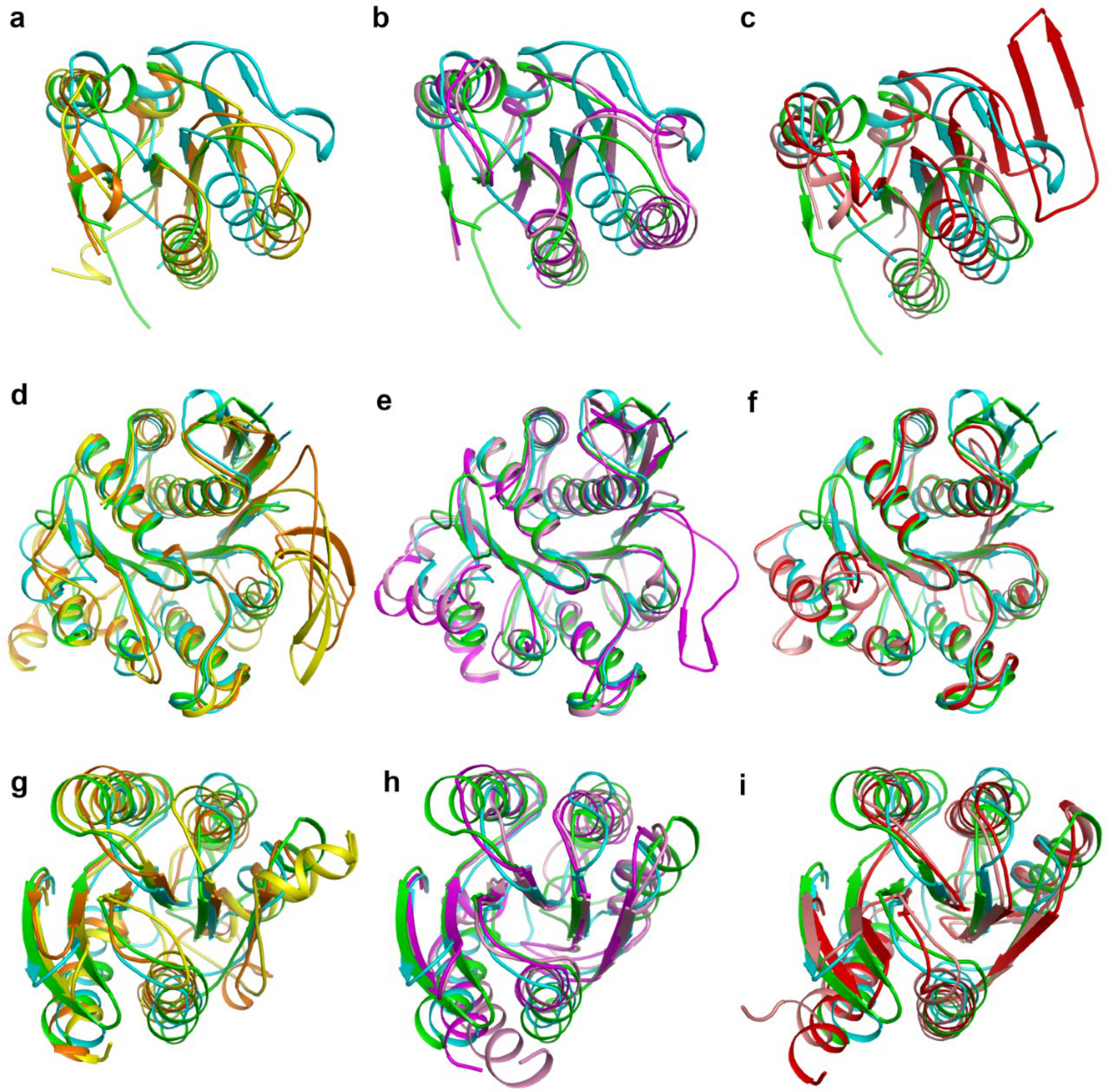


**Figure S39. Superimposition of the three domains of MurC, MurD, Murα, Murß and Murγ.** (**a**) Superimposition of the N-terminal domain of MurC from *Haemophilus influenzae* (PDB code 1P3D; green), MurD from *E. coli* (PDB code 4UAG; cyan), and Murα from *M. fervidus* (orange) and *M. smithii* (yellow) (**b**) same as (a) with Murß from *M. fervidus* (magenta) and *M. smithii* (pink) (**c**) same as (a) with Murγ from *M. fervidus* (salmon) and *M. smithii* (red). (**d, e, f**) Same as (a, b, c) for the middle domain. (**g, h, i**) Same as (a, b, c) for the C-terminal domain.
