## Supplemental Data for "Origin and Evolution of Pseudomurein Biosynthetic Gene Clusters"

#### Background

In this study, our bioinformatic pipeline (Fig. S1) allowed us to identify five syntenic regions (termed clusters A to E), which are conserved across the five pseudomurein (PM)-containing archaea (Fig. S2). Two of these regions, clusters A and B, are probably involved in PM biosynthesis, while cluster C seems to be involved in cell surface proteins (e.g., pili) and cell shape determination (or gene regulation). In contrast clusters D and E appear unrelated to such processes. Based on the genetic environment of clusters A to C, we attempted to identify a conserved regulon for PM biosynthesis and cell surface regulation. A regulon is a group of genes that are under the control of the same regulatory element (Cristianini and Hahn 2006; Anderssen et al. 2022).

#### Material and Methods

In order to determine whether the genes located in the three clusters are regulated by same transcription factors (TFs), we extracted the DNA sequences of the intergenic regions (IRs) if their length were at least 50 nucleotides (nt) long (or less if the direction of the upstream coding region was in reverse orientation compared to the considered gene). TATA-box and GpC island predictions were performed on IRs using respectively funzznuc (pattern ‘TATAWNNN’) and newcpgreport (window size of 50 and minimum length of 25) from EMBOSS package version 6.6.0.0 (Rice et al. 2000). Then, IRs between 50 to 75 nt with no TATA-box nor GpC island were discarded. MEME (from the MEME Suite; Bailey et al. 2009) was used with different combinations of IRs (Table S4) to identify DNA motifs that could be considered as TF binding sites. MEME was configured to find three motifs with a maximum length of 30 nt and using the DNA alphabet. Predicted DNA motifs were uploaded on the online tool PREDetector (Tocquin et al. 2016) with default parameters to identify new candidate genes with similar motifs in their regulatory regions in the genomes of the five PM-containing archaea. The resulting TSV files were filtered to retain only “upstream” and “regulatory” predictions (see PREDectector documentation) and the gene loci were used to fetch the corresponding protein accessions from GeneSpy (Garcia et al. 2019) GFM files.

### Results

DNA motifs identified with MEME did not show significant E-values (from 1.5e-002 to 7.1e+003), indicating that MEME struggled to discover reliable motifs. Nevertheless, we retained the five best motifs across all combinations to identify genes presenting potential TF-binding sites. Despite MEME not using all input sequences to discover motifs (Table S4), we searched for the discovered motifs in all PM-containing archaea. However, quite unsurprisingly, PREDectector predictions solely worked in organisms from which DNA sequences had been used for motif discovery, except in one case: while no sequence from *Methanobrevibacter smithii* had been used for motif prediction, some gene loci were identified in this organism using the second motif. The OGs of the corresponding protein products were then filtered using classify-ali.pl (see Material and Methods in the main text) keeping only those with proteins found in the five PM-containing archaea or at least four of them (one Methanopyrales and three Methanobacteriales), thereby allowing one missing gene in Methanobacteriales. In total, 112 OGs with at least five PM-containing archaea proteins and 19 with at least four were identified, of which 21 OGs had already been identified in the main pipeline for identifying PM biosynthesis candidate proteins (see main text). Among the new OGs, two were identified with different motifs (Table S1, sheet 5), (1) OG0000311 (a glutamate--tRNA ligase) with motifs 1, 4 and 5 in *Methanopyrus sp.*, yet its upstream region (**M-a**; see Figure S2 and Table S4) had been used for motif discovery, (2) OG0000359 with motif 2 in *Methanopyrus sp.* and with motif 3 in *Methanobacterium congolense*. It is a single-copy gene present in the ten archaea and it corresponds to the transcription factor Pcc1, which regulates the cell cycle and polar growth (Kisseleva-Romanova et al. 2006). To investigate whether OG0000359 could actually include the transcription factor regulating PM biosynthesis, we performed the same pipeline of analysis (see Material and Methods above). Due to the difficulty to define a DNA upstream sequence in *Methanothermus fervidus*, we did not include it for motif discovery. The best motif identified by MEME in the upstream region of OG0000359 genes is an 8-nt motif with an E-value of 9.0e-002. It allowed us to identify 185 OGs with at least five PM-containing archaea proteins and 40 with at least four of them. As above, out of these 225 OGs, 25 had already been identified (see main text). The newly identified OGs were then intersected with those selected using classify-ali.pl (i.e., multi-copy genes in PM-containing archaea (paralogs) but existing in a single copy in other archaea). Unfortunately, none of the new OGs passed this filter. Unlike bacteria (Anderssen et al. 2022), the obtained results showed that such a regulon-oriented pipeline is ineffective when predicting regulatory elements in archaea. However, it is not clear whether it is only ineffective in this specific case or if the approach cannot be applied to any extant archaea.
